# Single cell tri-channel-processing reveals structural variation landscapes and complex rearrangement processes

**DOI:** 10.1101/849604

**Authors:** Ashley D. Sanders, Sascha Meiers, Maryam Ghareghani, David Porubsky, Hyobin Jeong, M. Alexandra C.C. van Vliet, Tobias Rausch, Paulina Richter-Pechańska, Joachim B. Kunz, Silvia Jenni, Benjamin Raeder, Venla Kinanen, Jürgen Zimmermann, Vladimir Benes, Martin Schrappe, Balca R. Mardin, Andreas Kulozik, Beat Bornhauser, Jean-Pierre Bourquin, Tobias Marschall, Jan O. Korbel

## Abstract

Structural variation (SV), where rearrangements delete, duplicate, invert or translocate DNA segments, is a major source of somatic cell variation. It can arise in rapid bursts, mediate genetic heterogenity, and dysregulate cancer-related pathways. The challenge to systematically discover SVs in single cells remains unsolved, with copy-neutral and complex variants typically escaping detection. We developed single cell tri-channel-processing (scTRIP), a computational framework that jointly integrates read depth, template strand and haplotype phase to comprehensively discover SVs in single cells. We surveyed SV landscapes of 565 single cell genomes, including transformed epithelial cells and patient-derived leukemic samples, and discovered abundant SV classes including inversions, translocations and large-scale genomic rearrangements mediating oncogenic dysregulation. We dissected the ‘molecular karyotype’ of the leukemic samples and examined their clonal structure. Different from prior methods, scTRIP also enabled direct detection and discrimination of SV mutational processes in individual cells, including breakage-fusion-bridge cycles. scTRIP will facilitate studies of clonal evolution, genetic mosaicism and somatic SV formation, and could improve disease classification for precision medicine.

---

Cancer is largely a disease of the genome, which is caused by the clonal expansion of cells driven by mutation and selection. SVs represent the leading class of somatic driver mutation in different cancer types^1,2^. They comprise copy-number alterations (CNAs) and copy-balanced SVs mediating gene amplification, disruption, fusion, or enhancer hijacking^3–5^. SVs can arise in short evolutionary bursts through a variety of mechanisms, can precipitate further rearrangements during periods of extensive genomic instability, lead to vast genetic heterogeneity contributing to disease development and metastasis, and affect therapy responses^6–9^. A comprehensive understanding of the extent and nature of SVs in single cells is imperative to elucidate clonal evolution and mutational dynamics, and to unravel aberrant clonal expansions in normal and cancerous tissues^10,11^.

Important challenges have so far limited SV studies in genetically heterogeneous contexts. Methods for discovering different SV classes depend on the detection of discordantly aligned paired-end or split reads^12^. These methods require ≥20-fold genome coverage for clonal, and vastly higher coverage for subclonal SV discovery^13^. The exception is read-depth analysis, which can be pursued at lower coverage^10^, but which is restricted to detecting only CNAs. Translocations, inversions and complex rearrangements therefore largely escape detection in subclones, despite their known relevance in cancer and the relationship between complex rearrangement formation and disease prognosis^2,5,14^. While single cell analyses could help overcome these limitations^15^, scalable methods for single cell SV detection are likewise only suited for CNAs^16–18^. The systematic discovery of additional SV classes has been limited by the requirement to obtain uniformly high coverage in each cell using whole genome amplification (WGA)^17^, which can lead to read chimeras confounding SV calling. And while read chimera can be filtered when sequencing single cells to deep coverage^19,20^, SV surveys across tens to hundreds of cells using such deep sequencing approach would lead to prohibitive costs.

Here we describe a computational approach termed scTRIP (<u>s</u>ingle <u>c</u>ell <u>tri</u>-channel <u>p</u>rocessing) for systematic SV discovery in single cells. scTRIP leverages Strand-seq, a preamplification-free single cell technique based on labeling non-template strands during DNA replication^21^, which generates strand-specific read data and is suitable for SNP haplotype phasing^22^. While Strand-seq has been used to identify polymorphic (germline) inversions based on changes in read orientation^21,23^, efforts to exploit strand-specific reads for characterizing additional somatic SV classes and to deconvolve subclonal SV heterogeneity have been lacking. scTRIP now unlocks the full potential of Strand-seq, rendering a wide variety of SV classes accessible to systematic single cell studies. It does so using a joint calling framework that integrates three data layers - depth-of-coverage, read orientation, and haplotype phase - from strand-specific data. This enabled us to build complete SV landscapes in single cells, measure SV mutational processes and characterize somatic heterogeneity in leukemic samples. Distinct from prior approaches that characterize somatic SVs, scTRIP detects variants without requiring SV breakpoint-traversing paired or split reads, which enables robust discovery of somatic SVs at unprecedented scale using low-pass single cell sequencing.

## Results

### scTRIP enables systematic discovery of a wide variety of SV classes in single cells

The underlying rationale of scTRIP is that each class of SV can be identified via a specific ‘diagnostic footprint’. These diagnostic footprints capture the co-segregation patterns of rearranged DNA segments made visible by sequencing single strands of each chromosome in a cell, as follows: During S-phase, the DNA double strand unwinds, and the two resulting single strands (Watson [‘W’] and Crick [‘C’]) act as templates for DNA replication. In Strand-seq, newly replicated strands incorporate Bromodeoxyuridine (BrdU)^21^, which acts as a traceable label for these non-template strands (see **Fig. 1A** depicting the Strand-seq protocol)^24^. During mitosis, each of the two daughter cells receive one copy of each chromosomal homolog through independent and random chromatid segregation^21^. The labeled nascent strand is then removed, and the segregation pattern of each chromosomal segment is analyzed following strand-specific sequencing (**Fig. 1B**). scTRIP combines this strand-specific segregation information with read depth and haplotype phase information to capture newly-defined diagnostic footprints that characterize each SV class (**Fig. 1C-F**).

**Fig. 1.**
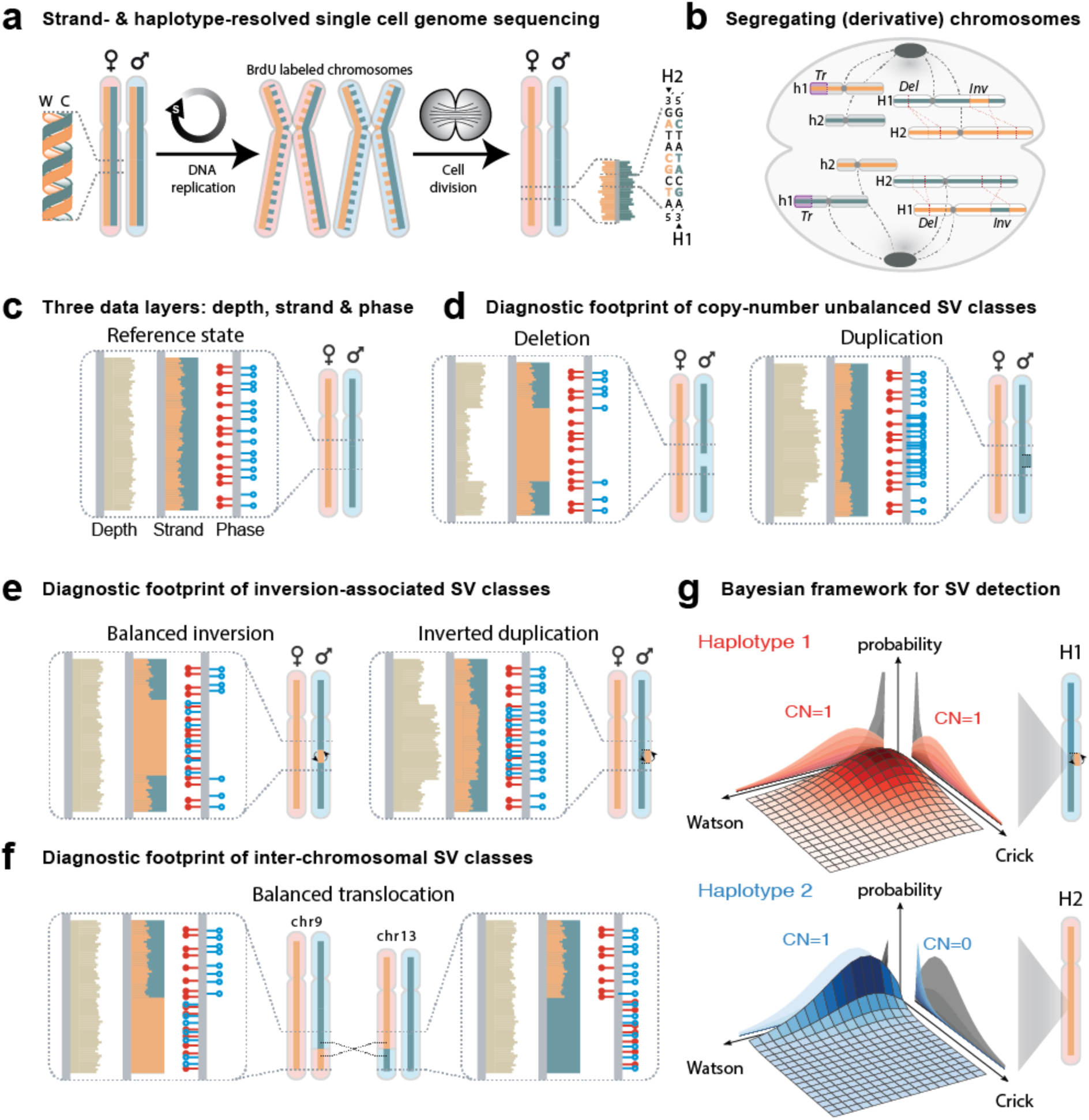
Haplotype-aware discovery of SVs in single cells. (**a**) Overview of the Strand-seq sequencing protocol. Strand-seq involves incorporating BrdU into dividing cells, followed by removal of the BrdU containing strands through nicking, and short read sequencing of the remaining strand^21^. Strand-seq libraries preserve strand-orientation and chromosomal homolog (haplotype) identity. Dashed line: strand (BrdU) label. W, Watson strand (orange); C, Crick (green); H, haplotype. (**b**) Scheme depicting how template strand co-segregation patterns during mitosis reveal SVs in single cells. *Del*, deletion; *Inv*, inversion; *Tr*, translocation. Segments of derivative chromosomes share the same template strand during DNA replication. H1/H2, haplotype 1 and 2 of a chromosome; h1/h2, haplotype 1 and 2 of another chromosome. (**c**) The scTRIP computational approach exploits three data layers: read depth, strand ratio, and chromosome-length haplotype phase. Red lollipops: reads assigned to H1 based on overlapping SNPs; blue lollipops: reads assigned to H2. Haplotype phase is assessed in a strand-aware fashion, with phased W reads shown as lollipops on left of ideogram and phased C reads shown to right. In contrast to prior SV detection approaches, scTRIP does not depend on discordant or split reads, the scalable detection of which has been considered infeasible in single cells. Panels **d-f** depict diagnostic footprints for chromosomes where both haplotypes are labeled on different strands (‘WC/CW chromosomes’). Our framework also detects and scores equivalent footprints on CC and WW chromosomes (see **Table S1**). (**d**) Del, detected as losses in read depth affecting a single haplotype, combined with unaltered read orientation. Dup, detected as a haplotype-specific gain in depth with unaltered read orientation. (**e**) Balanced Inv, identified as haplotype-phased read orientation ‘flips’ with unaltered depth. InvDup, characterized by inverted reads detected for one haplotype coinciding with a read depth gain of the same haplotype. (**f**) Balanced translocation, detected as correlated template strand switches affecting the same paired genomic regions in cells harboring the SV. (**g**) Bayesian framework for SV discovery. The probability distributions depicted represent an InvDup on H1 (segments on both strands are seen for haplotype 1 (H1), whereas H2 is represented on the W strand only).

The diagnostic footprint of deletions (Del) is defined by read depth losses affecting a single haplotype, coupled with unaltered read orientation (**Fig. 1B,D** and **Table S1**). Duplications (Dup) are characterized by haplotype-specific gains with unaltered orientation (**Fig. 1D, right panel**). In the case of balanced inversions (Inv), read orientation is altered with the re-oriented reads mapping to a single haplotype at constant read depth (**Fig. 1B,E**). Re-oriented reads co-locating with a read depth gain on the re-oriented haplotype signify an inverted duplication (InvDup; **Fig. 1E, right panel**). In the case of inter-chromosomal SVs, physically connected segments will co-segregate during mitosis, allowing the discovery of translocations. This is because segments originating from different chromosomes will now be adjacent to each other and thus will receive the same non-template strand-label during replication (**Fig. 1B**). Segments showing correlating strand states in different cells without a change in read depth characterize balanced translocations (**Fig. 1F**), whereas unbalanced translocations exhibit a similar footprint in conjunction with a read depth gain of the affected haplotype (**Fig. S1)**. Finally, altered cellular ploidy states also exhibit their own diagnostic footprints (**Fig. S2**).

To exploit these diagnostic footprints we developed a joint calling framework enabling the systematic discovery of SVs on a cell-by-cell basis. Described in detail in the **Supplementary Information** file, the framework first aligns, normalizes and places strand-specific read data into genomic bins, and assigns template strand states and chromosome-scale haplotypes for all cells (**Fig. S3**). It then identifies putative SVs by segmentation (**Methods**), and using a Bayesian model estimates genotype likelihoods for each segment and each single cell (**Fig. 1G**). The model integrates read depth, strand and haplotype phase signals to predict the most probable SV class described by our diagnostic footprints (**Fig. S4**). By performing SV discovery in a haplotype-aware manner, our joint calling framework also combines signals across cells (**Methods**) to sensitively detect subclonal SVs in a heterogeneous cell population. Finally, by analyzing adjacent SVs arising on the same haplotype it can unravel complex rearrangements, an abundant class of somatic structural variation in cancer^25,26^. As a first benchmark, we performed simulation experiments and observed excellent recall and precision after randomly placing SVs into cell populations *in silico*, even down to a single cell (**Fig. S5**).

### SV landscapes of RPE cells uncovered by scTRIP

To investigate single cell SV landscapes using scTRIP we next generated strand-specific DNA sequencing libraries from telomerase-immortalized retinal pigment epithelial (RPE) cells. We used hTERT RPE cells (RPE-1), commonly used to study patterns of genomic instability^20,27–29^, and additionally C7 RPE cells, which show anchorage-independent growth used as an indicator for cellular transformation^30^. Both RPE-1 and C7 cells originate from the same anonymous female donor. We sequenced 80 and 154 single cells for RPE-1 and C7, respectively, to a median depth of 387,000 mapped non-duplicate fragments (**Table S2** and **Methods**). This amounts to only 0.01X genomic coverage per cell.

We first searched for Dels, Dups, Invs and InvDups. Following read normalization (**Fig. S6**), we identified 54 SVs in RPE-1, and 53 in C7 cells (**Table S3**). 22 SVs were present only in RPE-1, and 21 were present only in C7, and thus likely correspond to sample-specific SVs (*i*.*e*. SVs that formed somatically or in the cultured cells rather than corresponding to germline variants; hereafter simply referred to as ‘somatic SVs’). Two representative SVs are shown in **Fig. 2A**, including a 1.4 Megabase (Mb) somatic Dup seen in RPE-1, and a 800 kb somatic Del detected in C7. While all but one of the Dels and Dups events were unique to RPE-1 and C7, Inv and InvDup events, including an Inv on 17p shown in **Fig. 2B**, were largely shared between both. These variants mapped to sites of known inversion polymorphisms^23^ (**Table S3**). We also identified chromosome arm-level CNAs, including deletion of 13q in C7, and duplication of a large 10q region in RPE-1. The 13q-arm showed a 1:0 strand ratio diagnostic for monosomy (**Fig. S2** and **Fig. 2C**), whereas the gained 10q region exhibited 2:1 and 3:0 strand ratios diagnostic for trisomic regions (**Fig. 2D** and **Table S4**).

**Fig. 2.**
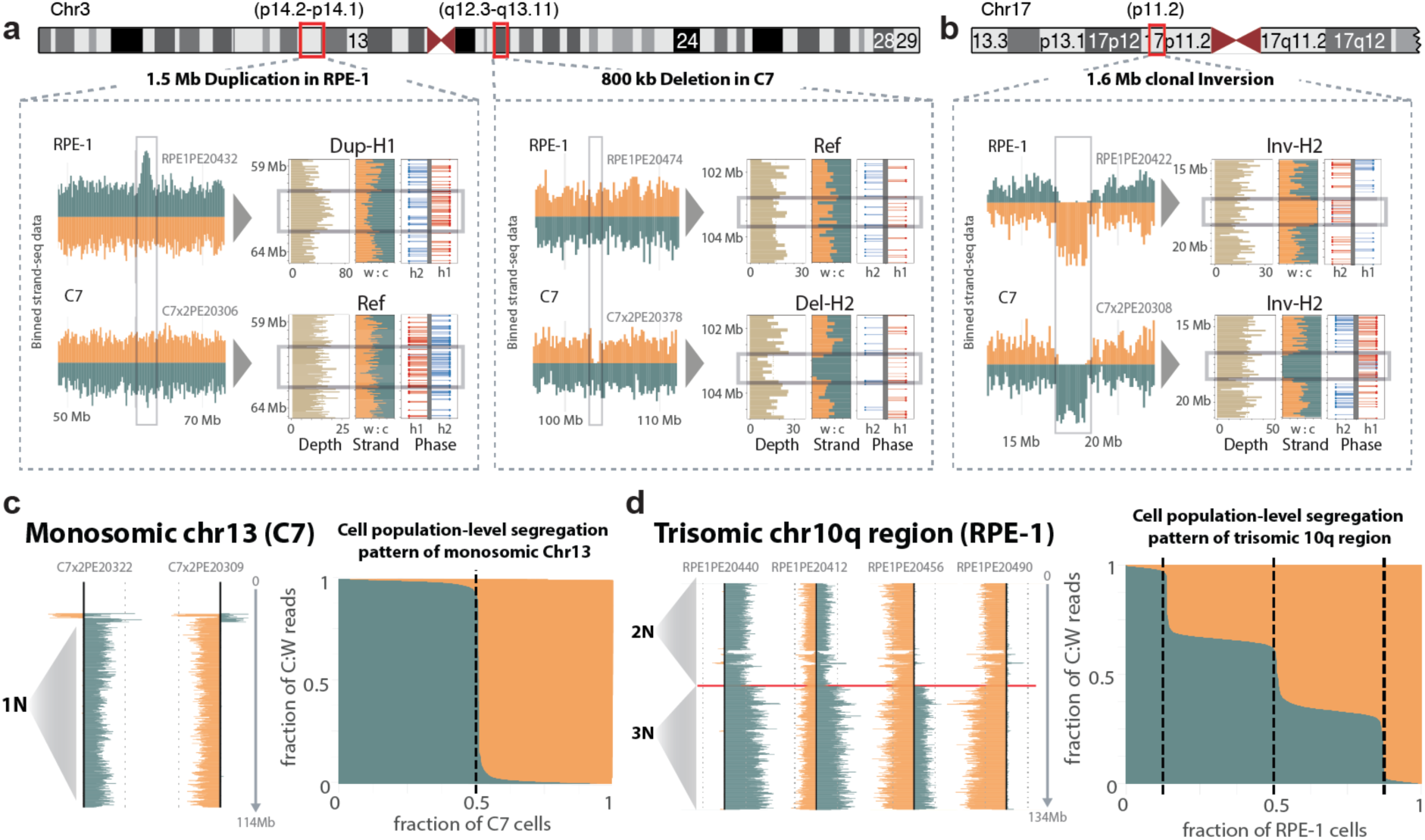
scTRIP reveals deletions, duplications, inversions and chromosome aneuploidies in epithelial cells. (**a**) Binned read counts separated by DNA strand and haplotype reveal the presence of SVs in single cells (W, Watson strand (orange); C, Crick (green)). Left panel: haplotype-resolved duplication (Dup) on 3p, which is present in RPE-1 but absent in C7. Right panel: haplotype-resolved deletion (Del) on 3q present in C7 and absent in RPE-1. The box ‘Depth’ depicts read counts; ‘Strands’ depicts the W:C fraction; ‘Phase’ shows the location of haplotype-phased SNPs, with lollipop orientation reflecting the strand state of the read containing the SNP (W on the left, C on the right, of the ideogram). (**b**) Chromosome 17p haplotype-resolved inversion (Inv) shared across both C7 and RPE-1. (**c**) Diagnostic footprint of a monosomic chromosome. Template strand state patterns depicted are from C7, which has a karyotypically defined^30^ monosomy 13. The left panel shows chromosome 13 strand-patterns from two single cells, with a visible 1:0 pattern characteristic for monosomy (1N). The right panel summarizes the fraction of observed W and C reads across 154 sequenced cells. (**d**) Diagnostic footprint of a trisomic region. Template strand state patterns depicted are from RPE-1 cells exhibiting a karyotypically defined 10q trisomic region^27^. The left panel shows chromosome 10 strand-patterns from four single cells. The right panel summarizes the fraction of observed W and C reads for the trisomic (3N) 10q region across 80 sequenced cells, revealing 2:1 and 3:0 strand ratios characteristic for trisomy (**Table S4**).

We benchmarked scTRIP by different means. First, we attempted to verify SVs present with ≥30% variant allele frequency (VAF) using bulk genome sequencing, which confirmed 12/12 (100%) of somatic SVs (≥30% VAF) in C7, and 15/16 (94%) in RPE-1 (**Table S5**). Second, we examined sensitivity, and successfully identified 85% of all SVs ≥200kb in size out of those detected when employing the DELLY SV caller^31^ on the bulk genomic sequence data (**Table S3**). Third, we used an *in silico* cell mixing approach, focusing on validated SVs that are near clonally present (VAF>90%) in C7 or RPE-1 and absent in the other, to assess the ability of scTRIP to identify subclonal SVs. This analysis showed that scTRIP is able to detect SVs at very low levels of VAF (<1% VAF), down to individual cells (**Fig. S7**). Fourth, we compared our approach with a prior computational method for CNA-profiling in single cells^18^. scTRIP outperformed this method even when limiting analyses to CNAs (**Fig. S8**).

### Dissecting complex cancer-related translocations in single cells

To assess the ability of scTRIP to detect a wider diversity of SV classes, we subjected RPE-1 cells to the CAST protocol^28^: we silenced the mitotic spindle machinery (**Supplementary Information**) to construct an anchorage-independent line (BM510) likely to exhibit genome instability. We sequenced 145 single BM510 cells detecting overall 67 SVs when searching for Dels, Dups, Invs and InvDups events (**Table S3**). Additionally, several DNA segments did not segregate with the respective chromosomes they originated from, indicating inter-chromosomal SV formation (**Fig. 3A**). We performed translocation detection with scTRIP searching for diagnostic co-segregation footprints (**Fig. 3B**), and identified four translocations in BM510 (**Fig. 3B,C**). We additionally subjected RPE-1 and C7 to translocation detection, identifying one translocation each (**Fig. 3D** and **Table S6**).

**Fig. 3.**
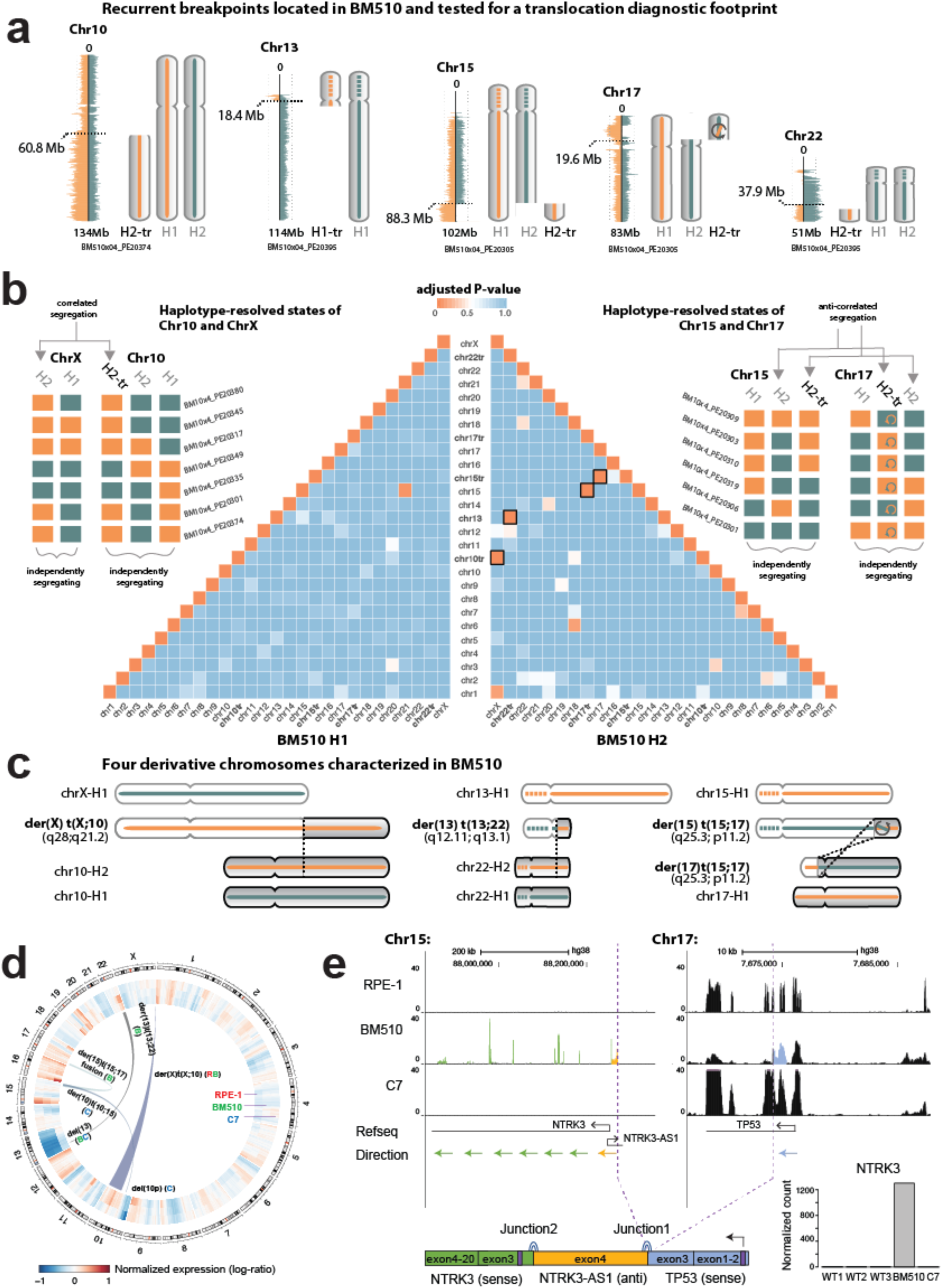
Translocation discovery in single cells. **(a)** In BM510, segments from chromosomes 10, 13, 15, 17 and 22 failed to co-segregate with the respective chromosomes they originated from, suggesting putative involvement in translocations (use of ‘tr’, as in “H2-tr’ or ‘chr10tr’, denotes the candidate translocation status of these segments). **(b)** Pyramid in the center: Unbiased analysis of translocations in BM510. Pairwise heatmap depicting segmental template-strand correlation values for each haplotype, highlighting the segment co-segregation diagnostic footprint (**Fig. 1F**) of translocations (correlation values are here expressed as Benjamini-Hochberg adjusted P-values). Orange boxes with black outline depict significant correlations (*P*<0.01; Fisher’s exact test) in four cases - corresponding to four derivative chromosomes we discovered in BM510. Schemes to the left and right: Colored boxes exemplify haplotype-resolved template strand state of segments for the non-reciprocal der(X) t(X;10) translocation and the t(15;17) reciprocal translocations. (In each case only a few cells are depicted for visualisation purposes.) Box colours: W (orange); C (green). Grey arrows highlight pairwise correlations between segments, where the paired segments always exhibit the same strand state (*e*.*g*. chrX and chr10tr), or always exhibit inverse strand states (e.g. chr15tr and chr17; reflecting inverted orientations of these translocation partners). Inversion within the translocated portion of 17p is denoted with a circular arrow. (**c**) Center: cartoon representation of four inferred derivative chromosomes. Dashed lines correspond to unassembled regions at acrocentric chromosomes 13 and 15. (**d**) Circos plot depicting translocations and averaged gene expression values across genomic windows^77^, computed from RNA-seq data generated for BM510 (here denoted *‘B*), RPE-1 (*‘R*’) and C7 (*‘C’)*. **Fig. S11** resolves expression by haplotype. (**e**) Validation of gene fusion in BM510. RNA-seq based read depth for *NTRK3* (green), *NTRK3-AS1* (yellow) and *TP53* (blue) is depicted for C7, RPE-1 and BM510. Purple dashed lines: detected fusion junctions. Lower left corner: inferred fusion transcript. Purple boxes show start codon locations. Lower right corner: *NTRK3* dysregulation in BM510. R1-3, RNA-seq replicates of RPE-1. Ex., exon.

One translocation was shared between RPE-1 and BM510 and involved the aforementioned gained 10q segment that underwent unbalanced translocation with a chromosome X haplotype (**Fig. 3B** and **Fig. S9**). We leveraged the footprints of sister chromatid exchange events^21^ to orient and order the segment (**Supplementary Information**), which placed the 10q gain to the telomeric tip of Xq, consistent with the published RPE-1 spectral karyotype^27^ (**Fig. 3C**). In BM510, scTRIP also uncovered balanced reciprocal translocations involving 15q and 17p (**Fig. 3B,C**). Notably, a *de novo* somatic inversion was additionally detected on the same 17p haplotype, which shared one of its breakpoints with the reciprocal translocation (**Fig. 3C** and **Supplementary Information**). Since these SVs shared one of their breakpoints, it is likely that both arose jointly, potentially involving a complex rearrangement process. Analysis of the locus revealed that the inversion encompassed the *TP53* gene, and upon translocating fused the 5’ exons of *TP53* to coding regions of the *NTRK3* oncogene^32^ (**Fig. 3E**). This suggests that scTRIP can reveal fusion genes using single cell sequence data.

Bulk whole genome sequencing (WGS) and RNA-Seq (**Methods**) analyses revealed excellent accuracy and specificity of our framework. We validated all translocations (100%), with 4/5 recapitulated by WGS and the remaining der(X) t(X;10) event by the existing karyotype data^27^. No additional translocation was detected in the deep sequencing data, indicating excellent sensitivity of scTRIP. WGS failed to verify the der(X) t(X;10) unbalanced translocation because the chrX breakpoint resides in highly repetitive telomeric DNA (resulting in ambiguous alignments hampering read pair analysis), whereas scTRIP uses mitotic co-segregation patterns not affected by repetitive breakpoints (**Fig. S10**). We also observed increased expression of the duplicated haplotype in the context of the der(X) t(X;10) event, corroborating our haplotype placements (**Fig. S11**). Finally, we verified the presence of the complex rearrangement at 17p, and uncovered expressed *NTRK* gene fusion transcripts exclusive to BM510 (**Fig. 3D,E**). Thus, scTRIP enables the haplotype-resolved discovery of translocations by single cell sequencing with high accuracy and sensitivity, which included detection of a translocation missed by bulk WGS.

### Single cell dissection of a complex DNA rearrangement process

Cancer genomes frequently harbor clustered SVs arising via complex rearrangements, which facilitate accelerated cancer evolution^33^. One process leading to such SVs are breakage-fusion-bridge cycles (BFBs)^34–39^. BFBs are initiated by the loss of a terminal chromosome segment, which causes newly replicated sister chromatids to fuse. The resulting dicentric chromosome will lead to a chromosomal bridge, the resolution of which via DNA breakage can initiate a new BFB^14^. Thus BFBs successively duplicate DNA segments in inverted orientation (*i*.*e*. generate InvDups), typically with an adjacent deletion of the terminal chromosome segment of the same haplotype (*i*.*e*. terminal deletion, here referred to as ‘DelTer’). BFBs rising to high VAF can be inferred from bulk WGS by analyzing ‘fold-back inversions’ (read-pairs aligning close to one another in inverted orientation)^34^. Owing to high coverage requirements, fold-back inversions cannot be systematically tracked in single cells. But we reasoned that scTRIP could provide the opportunity to directly study BFB formation in single cells.

To investigate BFBs we first turned to C7, in which fold-back inversions were previously described^28^. scTRIP located clustered InvDups on the 10p-arm in 152 out of 154 sequenced cells (**Fig. 4**). Closer analysis of 10p showed an amplicon with ‘stepwise’ InvDup events with an adjacent DelTer on the same haplotype, consistent with BFBs (**Fig. 4A-C** and **Fig. S12**). The remaining two cells lacking InvDups, notably, showed a larger DelTer affecting the same 10p segment (**Fig. 4C**). By aggregating sequencing reads across cells, we identified 8 discernable segments along chromosome 10, which included the 10p amplicon (comprising six copy-number segments) and its adjacent regions (the 10p terminal region, and the centromere-proximal region) (**Fig. 4B**). To further characterize the genetic heterogeneity seen at 10p, we inferred the cell-specific copy-number of all 8 segments (**Fig. 4D**). This revealed at least three distinct groups of cells with respect to 10p copy-number: (i) A large group presenting ‘intermediate’ copy-number with 100-130 copies detected for the highest copy-number segment (referred to as the ‘major clone’). (ii) Two cells that lost the corresponding 10p region through a DelTer, (iii) A single cell exhibiting vastly higher copy-number (∼440 copies), which may have undergone additional BFB cycles (**Fig. 4C** and **Fig. S13**).

**Fig. 4.**
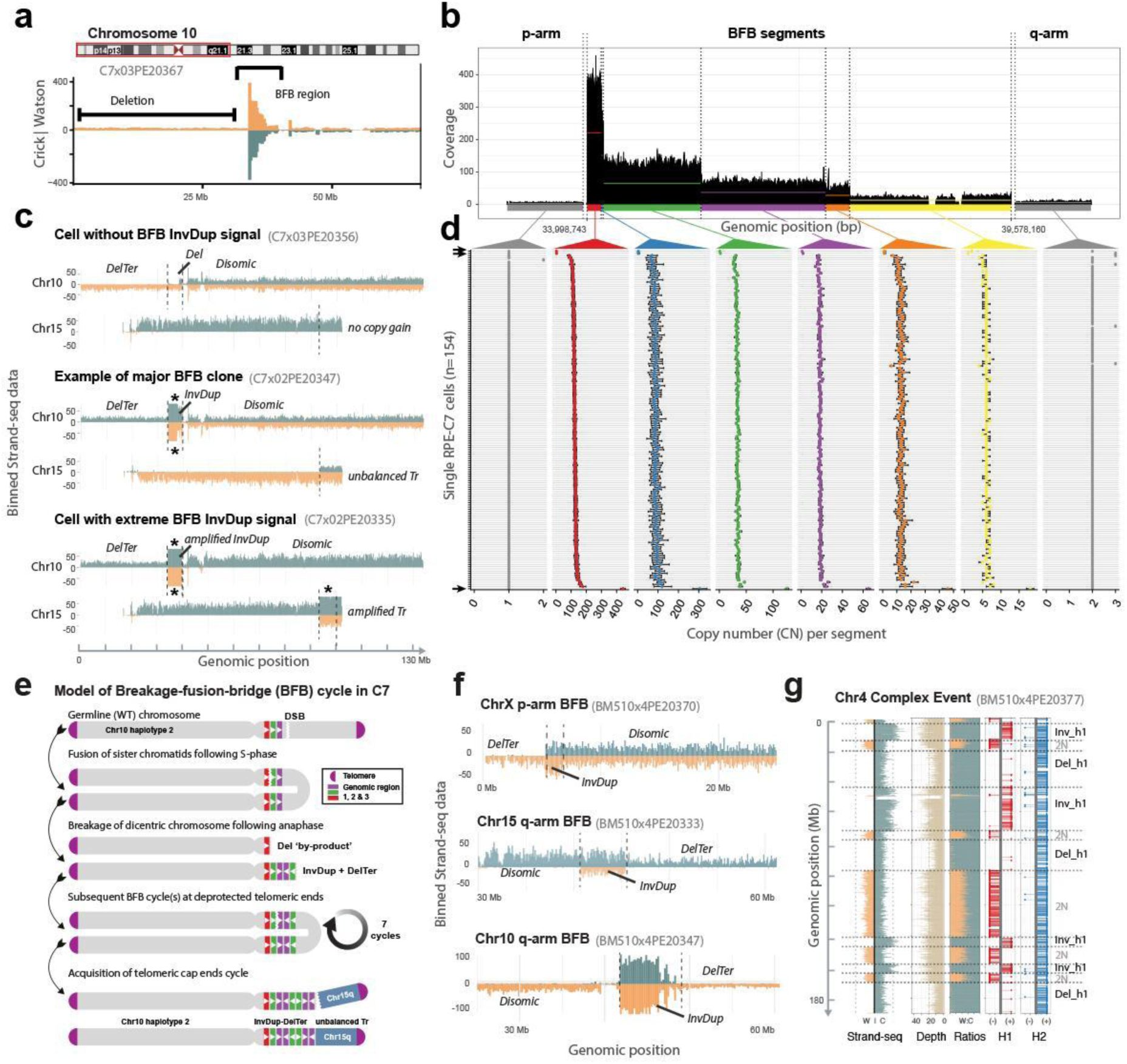
Single cell characterization of complex rearrangement processes. (**a**) Strand-specific read depth of C7 cell with region of InvDup mediated amplification on 10p, with adjacent terminal deletion (DelTer) of the same haplotype, resulting from BFB cycles. (**b**) Aggregated read data from 154 C7 cells. Colours indicate six copy number segments (red, blue, green, purple, orange and yellow) identified within the amplicon. Grey: regions flanking the amplicon. (**c**) Depiction of three C7 cells, with estimated maximum copy-number (CN) of 1 (upper panel), CN of ∼110 (middle panel), and CN of ∼440 (lower panel) at the 10p amplicon region indicated in red. A gained segment on 15q, which scTRIP inferred to have undergone unbalanced translocation with the amplicon region, is shown beneath (this SV is absent from cells lacking the amplicon; upper panel). Read counts for W (green) and C (orange) are capped at 50 (*, saturated read counts; see also **Fig. S13**, which uses a different y-axis scale). Tr, translocation. (**d**) Genetic diversity at 10p. CN (x-axis) is shown across 154 sequenced C7 cells (y-axis), providing cell-by-cell estimates of CN for each segment in (b). At least 3 different groups are readily discernible: high CN, intermediate CN, and loss of the 10p region (compare with panel (c)). Error bars reflect 95% confidence intervals. Arrows denote cells with CN=1 and CN of ∼440 at the 10p amplicon. (**e**) Model of sSVs leading to the observed structures seen for the ‘major clone’. Amplification via BFB cycles typically proceeds in 2^*n*^ copy-number steps, suggesting ∼7 successive BFB cycles occurred. According to our model, translocation of 15q terminal sequence stabilized 10p. DSB, double strand break. (**f**) The scar of BFBs, corresponding to InvDups flanked by DelTer on the same haplotype, identified in single BM510 cells. (**g**) Clustered rearrangements involving Dels and Invs in a single BM510 cell. Shown is the binned read data (left) seperated into the three data channels typical to scTRIP. All clustered SVs affect a single haplotype (H1, red).

Additional SVs identified in C7 provided further insights into the rearrangements occurring in the major clone: Namely, we detected an unbalanced translocation stitching a duplicated 15q segment to the 10p amplicon (**Fig. 4C** and **Table S6**). The duplicated segment encompassed the 15q telomere (**Fig. 4C**), which may have stabilized the amplicon to terminate the BFB process. In further support of C7 containing at least three groups of cells with respect to 10p structure, the unbalanced translocation was absent from the two cells harbouring the extended DelTer, whereas the translocated region became further amplified in the cell with excessive 10p copy-number (**Fig. 4C**). A model of the temporal sequence of rearrangements leading to the major clone is shown in **Fig. 4E**. These data underscore the ability of scTRIP to characterize BFB cycles for which direct measurement by single cell sequencing was not possible previously.

### Abundant BFB formation in anchorage-independent RPE cells

The frequency of BFB-mediated SV formation in somatic cells is unknown. Since scTRIP can systematically detect InvDup and DelTer footprints, we searched all sequenced RPE cells (379 in total) (**Methods**) and identified 15 additional cells exhibiting a BFB formation signature. Out of these, 11 displayed the ‘classical’ BFB footprint - an InvDup flanked by DelTer on the same homolog with no other SV present on the homolog (**Fig. 4F** and **Fig. S14**). The remaining four instances showed additional rearrangements on the same homolog as the BFB-associated SV. We tested whether InvDup-DelTer footprints coincided by chance by searching for structures where an InvDup on one haplotype was flanked by a DelTer on the other haplotype. Amongst the 379 cells, InvDup-DelTer footprints always occurred on the same haplotype, consistent with the well-known BFB model^38^. 11 out of the 15 InvDup-DelTer events occurred in BM510 affecting 8% (11/145) of the sequenced cells and 4 occurred in C7 affecting 3% (4/154) of the cells. No InvDup-DelTer footprints occurred in RPE-1 cells (0%; 0/80) and thus BFBs occurred exclusively in the transformed, anchorage-independently growing cells. Copy-number estimates of the InvDup regions ranged from 3 to 9, indicating that up to three BFB cycles occurred in these cells (**Fig. 4F**).

Interestingly, all of these 15 InvDup-DelTer footprints were singleton events detected in isolated cells (*i*.*e*. none were shared across more than one cell) and hence are likely to represent chromosomes with sporadically formed and potentially ongoing BFB cycles (**Fig. S14**). We reasoned that SVs identified in individual cells can serve as a proxy for currently active mutational processes. Using scTRIP, we systematically searched for other abundant SV mutational patterns in the RPE cell line in which we had induced genomic instability (BM510). We located 60 chromosomes with evidence of mitotic errors causing large (megabase-scale) deletion or duplication. Of these, 35/60 (58%) affected an entire homolog arm, 17/60 (28%) involved the tip of a homolog (terminal loss or gain) but not the whole arm, and 7/60 (12%) corresponded to whole homolog aneuploidy (monosomy or trisomy). The unifying characteristic of these abundant SV classes is that they can all result from mitotic segregation errors and reflect ongoing chromosome instability^40^.

Further underscoring this, nine cells showed multiple clustered SVs affecting the same haplotype. This included those four cells showing the InvDup-DelTer footprint and at least one additional SV. By employing the infinite sites assumption^37^, we inferred the relative ordering of SVs occurring on the same haplotype in these cases, identifying instances where the formation of additional SV preceded BFB formation, as well as such were the formation of additional SV succeeded BFB formation (**Fig. S15** and **SupplementaryMaterial**). This analysis also revealed a single cell exhibiting multiple reoriented and lost fragments, all on the same haplotype, resulting in 12 SV breakpoints affecting a single homolog. This rearrangement potentially resulted from a one-off rearrangement burst (chromothripsis)^41,42^ (**Fig. 4G**). Therefore, scTRIP enables systematic detection of *de novo* SV formation and discrimination of SV mutational processes, including BFBs and other complex rearrangements, in single cells.

### Constructing the karyotype of a PDX-derived T-ALL sample from 41 single cells

To evaluate the potential diagnostic value of scTRIP, we next analyzed patient-derived leukemic cells. Both balanced and complex SVs are abundant in leukemia, but largely escape detection in single cell studies geared towards CNAs^26,41,43^. We characterized PDX-derived^44^ samples from two T-cell acute lymphoblastic leukaemia (T-ALL) patients, to investigate the utility of scTRIP to characterize leukemic samples. We first focused on P33, a PDX-derived T-ALL relapse from a juvenile patient with Klinefelter Syndrome. We sequenced 41 single cells (**Table S2**) and used these data to reconstruct the haplotype-resolved karyotype of the major clone at 200kb resolution (**Fig. 5A**). While most chromosomes were disomic, we identified the typical XXY karyotype (Klinefelter Syndrome) and observed trisomies of chromosomes 7, 8, and 9. We further detected 3 regions of CNN-LOH characterized by haplotype-losses in the presence of constant read depth and orientation (**Fig. S16** and **Table S7**). Furthermore, we observed 6 focal CNAs, 5 of which affected genes previously reported to be genetically altered in and/or ‘driving’ T-ALL^43,45–47^, including deletions of *PHF6, RPL2* and *CTCF* sized 300kb and larger, as well as homozygous deletions of *CDKN2A* and *CDKN2B* (**Fig. 5A, Fig. S17** and **Table S3**). We also identified a t(5;14)(q35;q32) balanced translocation (**Fig 5A** and **Table S6**) - a recurrent rearrangement in T-ALL known to target *TLX3* for oncogenic dysregulation^48^. While few individual cells exhibited karyotypic diversity (**Fig. S17**), the majority of cells supported the karyotype of the major clone (**Fig. 5B**).

**Fig. 5.**
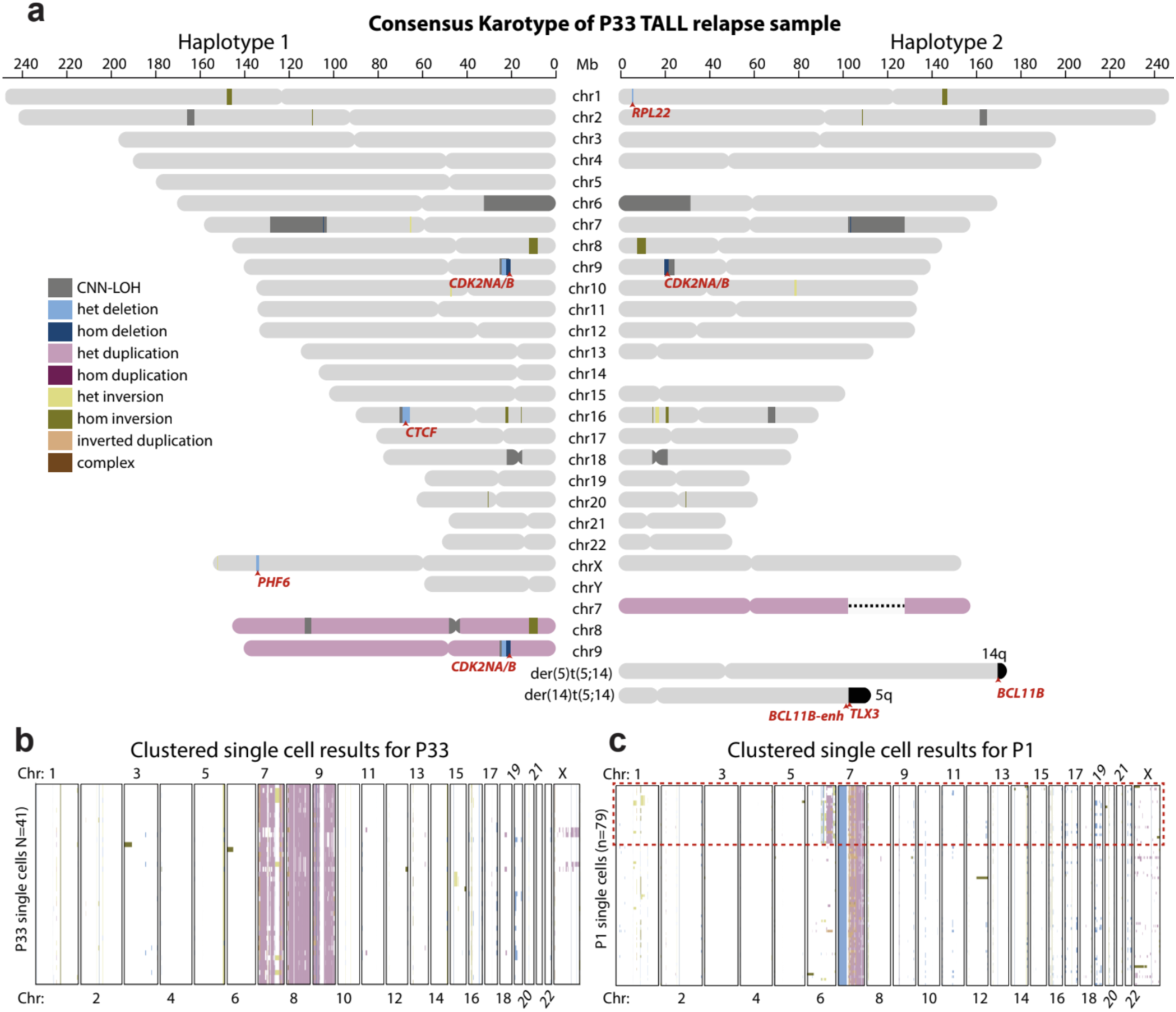
Single cell sequencing based karyotypes of PDX-derived T-ALL relapses. **(a)** Haplotype-resolved consensus P33 karyotype constructed from 41 sequenced cells, using single cell sequencing based SV calls generated by scTRIP. Heterozygous SVs are depicted only on the haplotype they have been mapped to. Homozygous SVs (by definition) appear on both haplotypes. CNN-LOH, copy-neutral loss in heterozygosity (shown on both haplotypes) ^78^. Chromosomes colored in pink reflect duplicated homologs. This T-ALL patient carries two chromosome X haplotypes (see also **Fig. S16**) as well as a Y chromosome, indicating transmission of an X and a Y chromosome from the father, whereas the mother contributed her X chromosome to the karyotype (Klinefelter or XXY syndrome). Affected leukemia-related genes are highlighted in red. ‘*BCL11B-enh*’ denotes a previously described enhancer region in 3’ of the *BCL11B* gene. **(b)** “Heatmap” of SVs arranged using Ward’s method for hierarchical clustering of SVs genotype likelihoods in P33, showing the presence of a single dominant clone and evidence of few additional somatic DNA alterations resulting in karyotypic diversity in this T-ALL relapse. **(c)** “Heatmap” of SV events called in an additional T-TALL sample, P1. Red dotted box outlines a clear subclonal population in the sample, represented by 25 cells.

We attempted verification of this karyotype with classical (cytogenetic) karyotyping obtained from primary T-ALL during diagnosis - the current clinical standard to genetically characterize T-ALL. Although this verified the duplications of chromosomes X, 7, 8 and 9, classical karyotyping failed to detect all the focal CNAs, and failed to capture the t(5;14)(q35;q32) translocation previously designed as ‘cryptic’ (*i*.*e*. ‘not detectable by karyotyping’)^49^. To verify the additional SVs detected by scTRIP, we next employed CNA profiling by bulk capture sequencing P33 at diagnosis, remission and relapse^50^ (**Supplementary Information**), as well as expression measurements. These experiments confirmed all (6/6, 100%) focal CNAs (**Table S3**), and verified *TLX3* dysregulation (**Fig. S18**) supporting the occurrence of a t(5;14)(q35;q32) balanced translocation. The haplotype-resolved karyotype inferred via scTRIP comprised SVs down to 200kb in size, located a ‘cryptic’ translocation missed by clinical karyotyping, and was built using sequence data from 41 cells, amounting to only ∼0.9× cumulative genomic coverage.

### scTRIP uncovers previously unrecognized DNA rearrangements in a PDX-derived T-ALL

We next turned to a second T-ALL relapse sample obtained from a juvenile female patient (P1). We sequenced 79 single cells of P1 (**Table S2**) and discovered two subclones, each represented by at least 25 cells (**Fig. 5C** and **Table S3)**. We first focused on the clonal SVs, which included a novel 2.6Mb balanced inversion at 14q32 (**Fig. 6A**). Interestingly, one of the inversion breakpoints fell into the exact same 14q region affected by the P33 t(5;14)(q35;q32) translocation (**Fig. 6B**). Prior studies have shown that, depending on their precise breakpoint locations, t(5;14) translocations can target *TLX3* and *NKX2-5* oncogenes at 5q35 by repositioning enhancer elements at 14q35 into the vicinity of these oncogenes^43,51^.

**Fig. 6.**
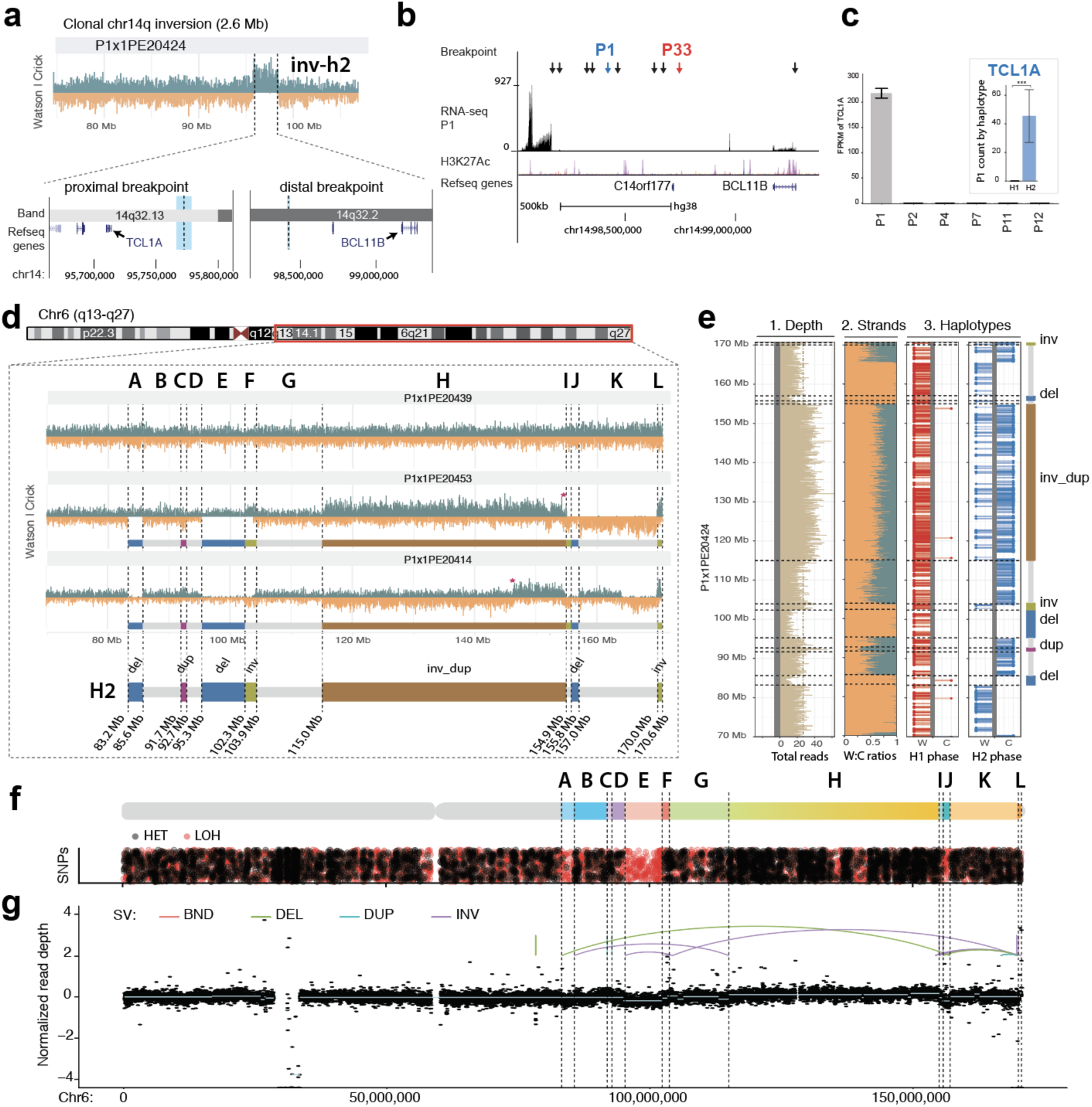
Single cell sequencing of PDX-derived T-ALL relapse P1 reveals previously unrecognized SVs. **(a)** Haplotype-resolved balanced 14q32 Inv inferred in P1 using scTRIP. The leftmost breakpoint (thick light blue line) resides close to *TCL1A*, whereas the rightmost breakpoint (thin light blue line) is in 3’ of *BCL11B*. (**b**) The rightmost Inv breakpoint falls into a “gene desert” region in 3’ *BCL11B* containing several enhancers. Black arrows show breakpoints of translocations resulting in T-ALL oncogene dysregulation from a recent study^45^. Colored arrows: SV breakpoints in T-ALL donors P1 and P33. (**c**) Dysregulation of *TCL1A* in conjunction with 14q32 Inv. Larger barplot shows *TCL1A* dysregulation in P1 compared to five arbitrarily chosen T-ALLs. Inset barplot shows allele-specific RNA-seq analysis demonstrating *TCL1A* dysregulation occurs only on the inverted (H2) haplotype. (**d**) Reconstruction of subclonal clustered DNA rearrangements at 6q via scTRIP. (**e**) Haplotype-resolved analysis of SVs clustered at 6q, all of which fall onto haplotype H2. (**f**) Detection of interspersed losses and retention of LOH in conjunction with the clustered SVs, indicative for a DNA rearrangements burst^41^. (LOH, signified by an abundance of red dots, was called as reported in the **Methods**. Regions with normal density of reference heterozygous SNPs (red), but with decreased density of additionally detected heterozygous SNPs (black), are indicative for LOH.) (**g**) Verification of subclonal clustered rearrangement burst at 6q, by bulk long-insert size paired-end sequencing^75^ to 165x physical coverage. Breakpoints inferred by scTRIP are shown as dotted lines, and scTRIP-inferred segments are denoted using the letters A to L. Colored breakpoint-connecting lines depict the paired-end mapping based rearrangement graph (*i*.*e*., deletion-type, tandem duplication-type, and inversion-type paired-ends). Using bulk whole-exome and mate-pair sequencing, read-depth shifts at these breakpoints were subtle and thus, this subclonal complex rearrangement escaped prior *de novo* SV detection efforts in bulk sequencing data.

The observation that both T-ALL patients showed balanced SVs affecting the same region motivated further analyses. This revealed that the novel 14q32 inversion we located in P1 juxtaposed the enhancer element-containing region 3’ of *BCL11B*^*48,51*^ into the immediate vicinity of the *T-cell leukemia/lymphoma 1A* (*TCL1A*) oncogene (**Fig. 6A** and **Fig. S18)**. Prior studies reported different enhancer-juxtaposing rearrangements in T-cell leukemia/lymphoma, as well as in T-ALL, resulting in *TCL1A* over-expression^52,53^, and we thus pursued RNA-seq to investigate differential expression in P1. This indeed confirmed *TCL1A* as the most highly overexpressed gene in P1 (>160-fold overexpression compared to five arbitrarily chosen T-ALLs; *P*=1.8E22 Wald test^54^, Benjamini-Hochberg correction; **Fig. 6C, left panel**). We reasoned that if *TCL1A* dysregulation had resulted as a consequence of the inversion, then *TCL1A* overexpression would be restricted to the rearranged haplotype. Leveraging scTRIP’s haplotype-resolved SV assignments, we performed allele-specific expression analyses, which showed that *TCL1A* overexpression indeed arose only from the haplotype carrying the inversion **(Fig. 6C, right panel**). These data implicate a novel inversion in driving oncogene expression. Further studies are needed to assess recurrence of this inversion in other T-ALL or T-cell malignancies, and to investigate the diversity of oncogene-dysregulating SVs involving the *BCL11B* enhancer. Due to its ability to perform scalable discovery of balanced SVs by shallow sequencing, scTRIP would be well-placed to investigate these questions in the context of larger patient cohorts.

We next analyzed subclonal SVs in P1, and discovered a series of highly clustered subclonal rearrangements affecting a single 6q haplotype (VAF=0.32). These rearrangements comprised two Invs, an InvDup, a Dup, and three Dels, resulting in overall 13 detectable breakpoints spanning nearly 90 Mb of 6q (**Fig. 6D,E)**. All cells exhibiting SVs at 6q showed evidence for the full set of 13 breakpoints. Furthermore, the copy-number profile oscillated between only three copy-number states^41^, and we observed islands of retention and loss in heterozygosity^41^ (**Fig. 6F**) - a rearrangement pattern that is reminiscent of chromothripsis^41,42^. To corroborate these data, we performed long (4.9kb) insert size mate-pair sequencing in bulk to deep (165x) physical coverage. While read depth alterations were barely discernible for this subclonal complex rearrangement, deep mate-pair sequencing confirmed all 13 subclonal SV breakpoints - thus verifying a subclonal rearrangement burst consistent with chromothripsis (**Fig. 6G**). These data underscore the ability of scTRIP to reveal subclonal complex SVs likely to be missed by standard bulk WGS^42^.

## Discussion

scTRIP enables the systematic detection of a wide variety of SVs in single cells using a joint calling framework that integrates read depth, strand, and haplotype phase. It can call subclonal SVs down to VAF<1% and identify SV formation processes acting in single cells, addressing unmet needs of SV detection methods^10,13,26,55,56^. Previous single cell studies investigating distinct SV classes have done so by sequencing only relatively few selected cells deeply after WGA^10,17,57^. And while prior SV detection efforts using Strand-seq were limited to germline inversions^23^, the computational advance presented here enables systematic discovery of CNAs, balanced and unbalanced translocations, inversions, inverted duplications and the outcomes of complex SV formation processes including BFBs and chromothripsis - all in single cells. Notably, scTRIP is further able to resolve repeat-embedded SVs (exemplified by an unbalanced translocation exhibiting a breakpoint in telomeric DNA), a class of SV that remains largely inaccessible to standard WGS in bulk. Moreover, SVs detected by scTRIP are haplotype-resolved, which helps to reduce false positive calls and allows integrating allele-specific gene expression data^57,58^.

We showcase the ability of scTRIP to measure SV formation processes by identifying BFB cycles in up to 8% of cells from transformed RPE cells, indicating that SV formation via BFB cycles is markedly abundant in these cells. Although initially described ∼80 years ago^38^, scTRIP now allows the direct and unbiased measurements of BFBs in individual somatic cells. BFB cycles were the most abundant SV formation process identified after chromosomal arm-level and terminal loss/gain events, all of which can result from chromosome bridges^40,59^. BFB cycles occur in a wide variety of cancers^14^, can precipitate other mutation processes such as chromothripsis^37^, and correlate with disease prognosis^60^. BFB cycles have also been reported outside of somatic cells, that is, in cleavage-stage embryos following *in vitro* fertilization, as revealed by hybridization-based single cell analysis^58^. It is estimated that 20% of all somatic deletions, and >50% of the entirety of somatic SVs in cancer genomes^25,26^, arise as complex DNA rearrangements. By enabling direct and robust measurement of these rearrangement processes in single cells, scTRIP will facilitate future investigations on the role of complex SVs in clonal evolution.

Our study also exemplified a potential value for disease classification by surveying balanced and unbalanced SVs, complex SVs, and karyotypic heterogeneity in patient-derived leukemic cells. We constructed the molecular karyotype of a T-ALL sample at 200kb resolution using 41 single cells, amounting to only 0.9x genomic coverage. This revealed submicroscopic CNAs and oncogenic DNA rearrangements invisible to cytogenetic methods currently used in the clinic. Classical cytogenetics is typically pursued for only a limited number of metaphase spreads per patient and normally fails to capture low levels of karyotypic heterogeneity accessible to scTRIP. In one of the T-ALL patients we discovered a subclonal chromothripsis event, highlighting potential utility for disease prognosis, since chromothripsis has been associated with dismal outcome in leukemia^61^. Studies of aberrant clonal expansions in healthy individuals^10^ and lineage tracing in cancer patients^62^ may also be facilitated by scTRIP in the future. Another potential application area is in rare disease genetics, where scTRIP may help resolve “unclear cases” by widening the spectrum of accessible SVs leading to somatic mosaicism^56^. Furthermore, our framework can be used to assess genome integrity in conjunction with cell therapy, gene therapy, and therapeutic CRISPR-Cas9 editing, which can result in unanticipated (potentially pathogenic) SVs^63,64^. The ability of scTRIP to generate high-resolution karyotypes could be employed to detect the presence of such unwanted SVs to address safety concerns pertaining to these future therapies.

scTRIP utilizes strand-specific data generated by Strand-seq, which requires labeling chromosomes during replication. Therefore, non-dividing, apoptotic, or fixed cells cannot be sequenced. However, many key cell types are naturally prone to divide or can be cultured, which for example includes fresh or frozen stem and progenitor cells, cancer cells, cells in regenerating or embryonic tissues, iPS cells and cells from diverse model systems including organoids. Moreover, in the future the computational framework underlying scTRIP could be used with strand-specific methods generating reads in the absence of cell division^65^.

Our approach enables systematic studies of somatic SV landscapes with much less sequence coverage than WGA-based single cell methods. We demonstrated robust SV discovery using ∼2000-fold less reads than required for prior read-pair or split-read based methods^12^. Single cell sequencing to deep coverage, using WGA, can enable mapping somatic SVs <200kb in size, and thus will remain useful for detecting small CNAs or retrotransposons. Compared to scTRIP, however, WGA-based single cell analyses are subject to the limitations of paired-end analyses, including susceptibility to allelic dropouts, difficulties in detecting repeat-embedded SVs, limited scalability and high costs^17^. The combined reagent costs for Strand-seq are ∼15$ per cell, and the protocol is readily scalable (see **Methods**) meaning scTRIP enables systematic studies of SV landscapes in hundreds of single cells. Low-depth methods for CNA profiling in single cells, for which scalable methods exist, detect CNAs of 1 to 5 Mb in size^16,18^. These methods show promise for investigating subclonal structure, particularly in cancers with abundant CNAs, but miss key SV classes and fail to identify or discriminate between different SV formation processes.

In conclusion, the joint calling framework of scTRIP enables systematic SV landscape studies in single cells to decipher derivative chromosomes, karyotypic diversity, and to directly investigate SV formation processes. This provides important value over existing methods, and opens up new possibilities in single cell sequencing and genetic heterogeneity studies.

## Acknowledgements

We thank Wolfgang Huber, Oliver Stegle, Francesco Marass, and Peter Lansdorp for valuable discussions, and Tania Christiansen for assistance with software documentation. We acknowledge the Human Genome Structural Variation Consortium (HGSVC) for providing early access to Strand-seq data from normal lymphoblastoid (1000 Genomes Project / HapMap) cell lines used during the development of scTRIP. We further thank the Flow Cytometry Core Facility, particularly Malte Paulsen, for valuable assistance in automating single cell library production, and Cornelia Eckert for the provision of primary T-ALL relapse samples for engraftment. JOK acknowledges funding from an ERC Starting Grant (336045), and from the National Institutes of Health (3U41HG007497-04S1). TM acknowledges funding from the German Research Foundation (DFG) (391137747 and 395192176). Funding was also provided by the José Carreras Foundation (DJCLS 06R/2016) to JOK, AEK and JBK. AS and HY received funding through Humboldt postdoctoral fellowships.

## Author contributions

*Study conception*: A.D.S., T.M., J.O.K. *Development of SV diagnostic footprints by scTRIP*: A.D.S., S.M., M.G., D.P., T.M., J.O.K. *Strand-seq single cell library preparation workflow:* A.D.S., B.R., J.Z., V.B., J.O.K. *Generation of BM510 using the CAST protocol:* B.R.M., J.O.K. *Single cell sequencing library preparation in PDX-derived T-ALL samples*. A.D.S., S.J., B.R., B.B., J.-P.B., J.O.K. *Development of computational framework (named MosaiCatcher) for analysing scTRIP data:* S.M., M.G., D.P., A.D.S., T.R, T.M., J.O.K. *Development of Bayesian framework for SV classification:* M.G., S.M., D.P., T.R., J.O.K., T.M. *Cell mixing experiments and simulations:* S.M., T.R., T.M. *Translocation analysis:* A.v.V., A.D.S., D.P., J.O.K. *Analysis of clustered rearrangement processes:* A.D.S., D.P., M.G., T.R., T.M., J.O.K. *CNN-LOH analysis:* D.P., A.D.S., T.M. *Haplotagging*: M.G., D.P., A.D.S., T.M. *Bulk genomic DNA sequence analysis:* T.R., B.R. *T-ALL clinical and classical cytogenetic data:* P.R-P., J.B.K., M.S., A.K., B.B., J.-P.B. *T-ALL gene expression measurements:* P.R-P., J.B.K., S.J., B.B., J.-P.B., A.K *Allele-specific RNA-seq analyses*: H.J., B.R., J.O.K. *Supervision*: T.M., J.O.K. *Manuscript*: A.D.S., T.M., J.O.K *wrote the initial version of the manuscript, which was then edited and approved by all authors*.

## Methods

### Data Availability

Sequencing data from this study can be retrieved from the European Genome-phenome Archive (EGA), and the European Nucleotide Archive (ENA) [accessions: PRJEB30027, PRJEB30059, PRJEB8037, EGAS00001003248, EGAS00001003365]. Access to human patient data is governed by the EGA Data Access Committee.

### Code Availability

The computational code of our analytical framework is hosted on github (see https://github.com/friendsofstrandseq/pipeline, and https://github.com/friendsofstrandseq/mosaicatcher). As an important note for reviewers, while some portions of our code are already available for download from github, others have not yet been posted to github owing to a pending patent application (all code will be made available freely for academic research upon acceptance of the paper). To ensure our code is available during peer review process, we have uploaded *all* code that is not yet available at github as a *Related Manuscript ZIP file*.

### Cell Lines and Culture

hTERT RPE-1 cells were purchased from ATCC (CRL-4000) and checked for mycoplasma contamination. BM510 cells were generated using the CAST protocol and derived from the RPE-1 parental line (as previously-described in Mardin et al. 2015, and further detailed in the **Supplementary Information**). C7 cells were acquired from Riches et al 2001. Cell lines were maintained in DMEM-F12 medium supplemented with 10% fetal bovine serum and antibiotics (Life Technologies).

### Ethics Statement

The protocols used in this study received approval from the relevant institutional review boards and ethics committees. The T-ALL patient samples were approved by the University of Kiel ethics board, and obtained from clinical trials ALL-BFM 2000 (P33; age: 14 years at diagnosis) or AIEOP-BFM ALL 2009 (P1; age: 12 years at diagnosis). Written informed consent had been obtained from these patients, and experiments conformed to the principles set out in the WMA Declaration of Helsinki and the Department of Health and Human Services Belmont Report. The in *vivo* animal experiments were approved by the veterinary office of the Canton of Zurich, in compliance with ethical regulations for animal research.

### Single cell DNA sequencing of RPE and T-ALL cells

RPE cells and PDX-derived T-ALL cells were cultured using previously established protocols^28,66^. We incorporated BrdU (40µM; Sigma, B5002) into growing cells for 18-48 hours, single nuclei were then sorted into 96-well plates using the BD FACSMelody cell sorter, and strand-specific DNA sequencing libraries were generated using the previously described Strand-seq protocol^21,67^. The BrdU concentration used was recently shown to have no measurable effect on sister chromatid exchanges^24^, a sensitive measure of DNA integrity and genomic instability^24^. To generate libraries at scale, the Strand-seq protocol was implemented on a Biomek FX^P^ liquid handling robotic system, which requires two days to produce 96 barcoded single cell libraries. Libraries were sequenced on a NextSeq5000 (MID-mode, 75bp paired-end protocol), demultiplexed and aligned to GRCh38 reference assembly (BWA 0.7.15). High quality libraries (obtained from cells undergoing one complete round of DNA replication with BrdU incorporation) were selected as described in^21,67^. Briefly, libraries showing very low, uneven coverage, or an excess of ‘background reads’ yielding noisy single cell data were filtered prior to analysis. In a typical experiment, ∼80% of cells yield high quality libraries reflecting BrdU incorporation in exactly a single cell cycle. Cells with incomplete BrdU incorporation or cells undergoing more than one DNA synthesis phase under BrdU exposure are identified during cell sorting and thus get only rarely sequenced during Strand-seq experiments^21,67^, typically contributing to less than 10% of sequenced cells. Such ‘unusable libraries’ hence do not palpably contribute to experimental costs.

### Chromosome-length haplotype phasing of heterozygous SNPs

Our SV discovery framework ‘MosaiCatcher’ phases template strands using StrandPhaseR^22^. The underlying rationale is that for ‘WC chromosomes’ (chromosomes where one parental homolog is inherited as W template strand and the other homolog is inherited as C template strand), heterozygous SNPs can be immediately phased into chromosome-length haplotypes (a feature unique to strand-specific DNA sequencing). To maximize the number of informative SNPs for full haplotype construction we aggregated reads from all single cell sequencing libraries and an internal 100 cell control and performed SNP discovery by re-genotyping the 1000 Genomes Project (1000GP) SNP sites^68^ using Freebayes^69^. All heterozygous SNPs with QUAL>=10 where used for haplotype reconstruction and single cell haplotagging (described below).

### Discovery of deletions, duplications, inversions and inverted duplications in single cells

We developed the core workflow of ‘MosaiCatcher’ to enable single cell discovery of Dup, Del, Inv, and InvDup SVs. Input data to the workflow are a set of single-cell BAM files from a donor sample, aligned to a reference genome. The core workflow performs binned read counting, normalization of coverage, segmentation, strand state and sister chromatid exchange (SCE) detection, and haplotype-aware SV classification. A brief description of each step is provided below, and for additional details see **Supplementary Information**.

#### Binned read counting

Reads for each individual cell, chromosome and strand were binned into 100kb windows. PCR duplicates, improper pairs and reads with a low mapping quality (<10) were removed to count only unique, high-quality fragments.

#### Normalization of coverage

Normalization was performed to adjust for systematic read depth fluctuations. To derive suitable scaling factors, we performed an analysis of Strand-seq data from 1,058 single cells generated across nine 1000GP lymphoblastoid cell lines made available through the HGSVC project (http://ftp.1000genomes.ebi.ac.uk/vol1/ftp/data_collections/hgsv_sv_discovery/working/20151203_strand_seq/), and pursued normalization with a linear model used to infer a scaling factor for each genomic bin.

#### Joint segmentation of single cells in a population

Segmentation was performed by jointly processing strand-resolved binned read depth data across all single cells of a sample, used as multivariate input signal with a squared-error assumption^70^. Given a number of allowed change points *k*, a dynamic programming algorithm was employed to identify the discrete positions of *k* change points with a minimal sum of squared error. Analyzing all cells jointly in this way rendered even relatively small SVs (∼200kb) detectable once these are present with sufficient evidence in the single cell dataset (*e*.*g*. seen in enough cells). The number of breakpoints was chosen separately for each chromosome as the minimal *k*, such that using *k*+1 breakpoints would only yield a marginal improvement, operationalized as the difference of squared error terms being below a pre-selected threshold (**Supplementary Information**).

#### Strand-state and SCE detection in individual cells

The interpretation of strand-specific binned read counts relies on the knowledge of the underlying state of template strands for a given chromosome (WW, CC, or WC). These “ground states” stay constant over the length of each chromosome in each single cell, unless they are altered through SCEs^21,71^. To detect SCEs, we performed the same segmentation procedure described above in each cell *separately* (as opposed to *jointly* across all cells, as for the segmentation). We then inferred putative SCEs by identifying changes in strand state in individual cells that are otherwise incompatible with breakpoints uncovered by the joint segmentation (**Supplementary Information**). Leveraging these putative SCEs, we then assigned a ground state to each segment (**Supplementary Information**). To facilitate haplotype-resolved SV calling, we employed StrandPhaseR^72^ to distinguish segments with ground state WC, where Haplotype 1 is represented by Watson (W) reads and Haplotype 2 by Crick (C) reads, from ground state CW, where it is *vice versa*.

#### Haplotype-aware SV classification

We developed a Bayesian framework to compute posterior probabilities for each SV diagnostic footprint, and derive haplotype-resolved SV genotype likelihoods. To this end, we modeled strand-specific read counts using a negative binomial (NB) distribution, which captures the overdispersion typical for massively-parallel sequencing data ^54^. The NB distribution has two parameters, *p* and *r;* the parameter *p* controls the relationship of mean and variance and was estimated jointly across all cells, while *r* is proportional to the mean and hence varies from cell to cell to reflect the different total read counts per single-cell library. After estimating *p* and *r*, we computed haplotype-aware SV genotype likelihoods for each segment in each single cell: For a given ground state (see above), each SV diagnostic footprint translates into the expected number of copies sequenced in W and C orientation contributing to the genomic segment (**Table S1**), which gives rise to a likelihood with respect to the NB model. The fact that our model distinguishes WC from CW ground states (see *Strand-state and SCE detection* above) renders our model implicitly whole-chromosome haplotype-aware - a key feature not met by any prior approach for somatic variant calling in single cells. In addition to this, we also incorporated the count of W or C reads assignable to a single haplotype via overlapping SNPs in the likelihood calculation, and refer to this procedure as “haplotagging” (since it involves reads “tagged” by a particular haplotype). We modeled the respective counts of tagged reads using a multinomial distribution (**Supplementary Information)**. The output is a matrix of predicted SVs with probability scores for each single cell.

#### SV calling in a cell population

Our workflow estimates VAF levels for each SV and uses them to define prior probabilities for each SV (Empirical Bayes). In this way, the framework benefits from observing SVs in more than one cell, which leads to an increased prior and hence to more confident SV discoveries. Our framework adjusts for the tradeoff between sensitively calling subclonal SVs, and accurately identifying SVs seen consistently among cells. We parameterized this tradeoff into a ‘strict’ and ‘lenient’ SV caller, whereby the ‘strict’ caller optimizes precision for SVs seen with VAF ≥5%, and the ‘lenient’ caller targets all SVs including those present in a single cell only. Unless stated otherwise, SV calls presented in this study were generated using the ‘strict’ parameterization, to achieve a callset that minimises false positive SVs (**Supplementary Information)**. We explored the limits of these parameterizations using simulations, by randomly implanting Dels, Dups and Invs into single cells *in silico*. We analyzed 200 single cells per simulation, applying coverage levels typical for Strand-seq ^21^ (400,000 read fragments per cell). We observed excellent recall and precisions for SVs ≥1Mb in size when present with >40% VAF (**Fig. S5**). And while we detected a decrease in recall and precision for events present with lower VAF, we were able to recover smaller SVs and those with lower VAF down to individual cells (**Fig. S5**).

### Single cell dissection of translocations

We discovered translocations in single cells by searching for segments exhibiting strand-states that are inconsistent with the chromosomes these segments originate from, while being consistent (correlated, or anti-correlated) in strand-state with another segment of the genome (*i*.*e*., their translocation partner) (**Supplementary Information)**. To infer translocations, we determined the strand states of each chromosome in a homolog-resolved manner. In cases where strand states appeared to change across a haplotype (because this haplotype exhibited SVs or SCEs), we used the majority strand state (*i*.*e*. ‘ground state’, see above) to pursue translocation inference. We examined template strand co-segregation by generating contingency tables tallying the number of cells with equivalent strand states versus those not having equivalent strand states (see **Fig. 3B**). We employed Fisher’s exact test to infer the probability of the count distribution in the contingency table, followed by p-value adjustment^73^.

### Characterization of breakage-fusion bridge (BFB) cycles in single cells

To infer and characterize BFB cycles in single cells, we first employed our framework with lenient parameterization to infer InvDups flanked by a DelTer event on the same homolog/haplotype. We tested whether InvDup-DelTer footprints resulting from BFB cycles may arise in single cells by chance, by searching for structures where an InvDup on one haplotype would be flanked by a DelTer on the other haplotype (for instance, an InvDup (H1)-DelTer (H2) event, where H1 and H2 denote different haplotypes). No such structures were detected, and InvDup-DelTer footprints thus always occurred on the same haplotype, consistent with BFB cycle formation. To ensure high sensitivity of our single cell based quantifications shown in **Fig. S14**, we additionally performed manual inspection of the single cell data for evidence of at least one of the following rearrangement classes: (*i*) an InvDup, (*ii*) a DelTer resulting in copy-number=1 on an otherwise disomic chromosome. These cells were inspected for InvDup-DelTer patterns indicative for BFBs, based on the diagnostic footprints defined in **Fig. 1**.

### Single cell based CNN-LOH discovery

For CNN-LOH detection, our framework first assembles consensus haplotypes for each sample, by analyzing all single cell Strand-seq libraries available for a sample using StrandPhaseR^22^. Each single cell is then compared to these consensus haplotypes in a disomic context, to identify discrepancies matching the CNN-LOH footprint. To detect clonally present CNN-LOH events, we used the 1000GP^68^ reference SNP panel to re-genotype aggregated single cell libraries in each sample. These re-genotyped (observed) SNPs were then compared to the 1000GP reference sets to identify genomic regions showing marked depletion in heterozygous SNPs indicative for CNN-LOH. To this end, we downsampled the 1000GP reference variants to the SNP numbers observed in the single cell data, and subsequently merged both data sets (observed and reference variants), sorting all SNPs by genomic position. We performed a sliding window search through these sorted SNPs, moving one SNP at a time, and compared the number of observed and reference SNPs in each window by computing the ratio *R*=observed SNPs/reference SNPs. In heterozygous disomic regions, *R* values of ∼1 will be expected, whereas deviations are indicative of CNN-LOH. Window sizes (determined by the number of SNPs in a window) were defined as the median SNP count per 500kb window. We employed circular binary segmentation (CBS)^74^ to detect changes in *R*, and assigned each segment a state based on the mean value of *R*. Segments ≥2Mb in size exhibiting mean values *R*≤0.15 were reported as CNN-LOH.

### Bulk genomic DNA sequencing

Genomic DNA was extracted using the DNA Blood Mini Kit (Qiagen, Hilden, Germany). 300 ng of high molecular weight genomic DNA was fragmented to 100 – 700bp (300bp average size) with a Covaris S2 instrument (LGC Genomics) and cleaned up with Agencourt AMPure XP (Beckman Coulter, Brea, USA). DNA library preparation was performed using the NEBNext Ultra II DNA Library Prep Kit (New England Biolabs, Ipswich, USA). We employed 15ng of adapter ligated DNA and performed amplification with 10 cycles of PCR. DNA was size selected on a 0.75% agarose gel, by picking the length range between 400 and 500 bp. Library quantification and quality control was performed using a Qubit 2.0 Fluorometer (Thermo Fisher Scientific, Waltham, USA) and a 2100 Bioanalyzer platform (Agilent Technologies, Santa Clara, USA). WGS was pursued using an Illumina HiSeq4000 (Illumina, San Diego, USA) platform, using 150 bp paired-end reads. Mate-pair sequencing with large insert size (∼5kb) was pursued as described previously^75^. SV detection in bulk DNA sequence data was pursued using Delly2^31^. RPE-1 WGS data was sequenced to 32× coverage.

### Bulk RNA-seq

Total RNA was extracted from RPE cells using the RNeasy MinElute Cleanup kit (Qiagen, Hilden, Germany). RNA quality control was performed using the 2100 Bioanalyzer platform (Agilent Technologies, Santa Clara, USA). Library preparation was pursued with a Beckman Biomek FX automated liquid handling system (Beckman Coulter, Brea, USA), with 200 ng starting material using TruSeq Stranded mRNA HT chemistry (Illumina, San Diego, USA). Samples were prepared with custom 6 base pair barcodes to enable pooling. Library quantification and quality control were performed using a Fragment Analyzer (Advanced Analytics Technologies, Ames, USA). RNA-Seq was pursued on an Illumina HiSeq 2500 platform (Illumina, San Diego, USA), using 50 base pair single reads. For RNA sequencing in T-ALL, total RNA was extracted using TRIzol (Invitrogen Life Technologies). The RNA was than treated with TURBO DNase (Thermo Fisher Scientific, Darmstadt, Germany) and purified using RNA Clean&Concentrator-5 (Zymo Research, Freiburg, Germany). We required a minimal RIN (RNA Integrity Number) of 7 as measured using a Bioanalyzer (Agilent, Santa Clara, CA) with the Agilent RNA 6000 Nano Kit. Cytoplasmic ribosomal RNA was depleted by Ribo-Zero rRNA Removal Kit (Illumina, San Diego, CA) and the libraries were prepared from 1 µg of RNA using TruSeq RNA Library Prep (Illumina, San Diego, CA). These samples were sequenced on a Illumina HiSeq 2000 lane as 75 bp single ends. Fusion junctions were detected using the STAR aligner^76^.

### Quantitative real time PCR (qPCR)

RNA from PDX-derived T-ALL samples was extracted using a RNeasy Mini kit according to manufacturer’s instructions (cat 74106, Qiagen, Hombrechtikon, Switzerland), and cDNA was generated using High Capacity cDNA Reverse Transcription Kit (Applied BioSystems, Foster City, USA). qPCR was performed using a TaqMan Gene Expression Master Mix (Applied BioSystems) in triplicate using an ABI7900HT Analyzer with SDS2.2 software. Threshold cycle values were determined using the 2-ΔΔCT method, normalized to human-GAPDH (Hs02786624_g1, Applied BioSystems).

