## Supplemental Information for "Single cell tri-channel-processing reveals structural variation landscapes and complex rearrangement processes"

### Index:

This Supplementary Information is divided into Figures, Tables, Experimental Procedures, and References. The Experimental Procedures section is organized in the following way: In Part 1 we provides details about the computational framework 'MosaiCatcher', which we devised to enable systematic SV detection in single cells based on the principles of scTRIP. In Part 2, we describe a number of genomic analyses in greater detail than was possible in the main text, including the construction of the BM510 cell line and extensive SV verification experiments.


**Supplementary Figures (page 1)**

**Supplementary Tables (page 19)**

**Supplementary Experimental Procedures (page 22)**

*1.1 Core computational framework for SV discovery in single cells*

*1.2 Simulations to evaluate our framework for single cell SV discovery*

*1.3 Comparing our scTRIP-based framework with a CNA detection tool*

*2.1 Single cell dissection of complex inter-chromosomal rearrangements*

*2.2 Characterizing altered ploidy in RPE cells by single cell sequencing*

*2.3 Construction of BM510: an RPE cell line showing genomic instability*

*2.4 Temporal ordering of SVs in single cells using the infinite sites assumption*

*2.5 Verification of SVs and karyotypes inferred by single cell sequencing in PDX-derived T-ALL samples*

**Supplementary References (page 36)**

### Supplementary Figures

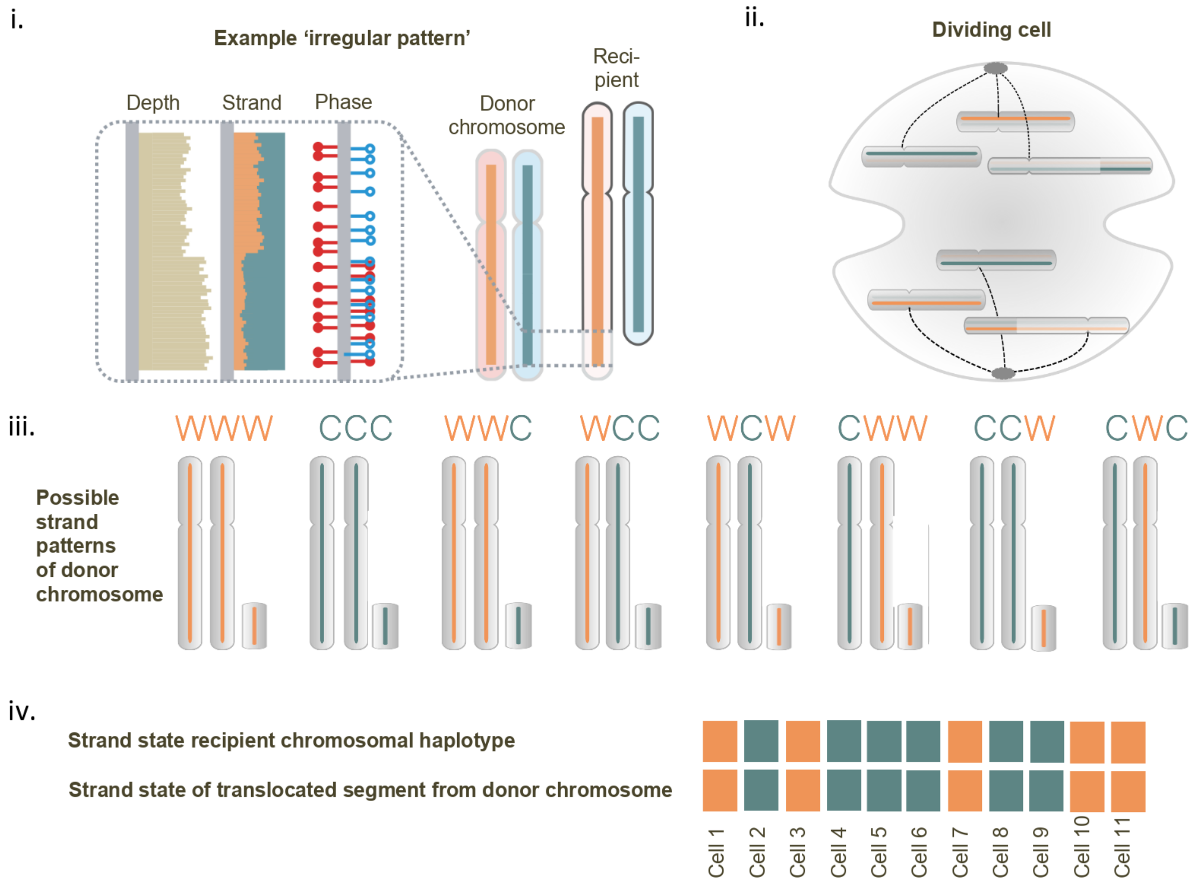


**Figure S1. SV diagnostic footprint of an unbalanced translocation.** In the event of an unbalanced translocation, the translocated segment of the donor chromosomal haplotype will exhibit a 'irregular' pattern, whereby read depth and strand-state is altered relative to the remainder of the donor chromosome (i.). This irregular pattern is due to the fact that the translocated chromosomal segment independently segregates during cell division (ii.), leading to eight different strand patterns with respect to the donor chromosome in the example shown here (iii.). Since the translocated segment co-segregates with the recipient chromosome during cell division (ii.), the segment will correlate in its strand state with the recipient chromosomal haplotype (see mock data for an example translocation shown in (iv.), with orange for W, and green for C; note: **Fig. 3** shows actual examples of translocations identified in RPE cells).

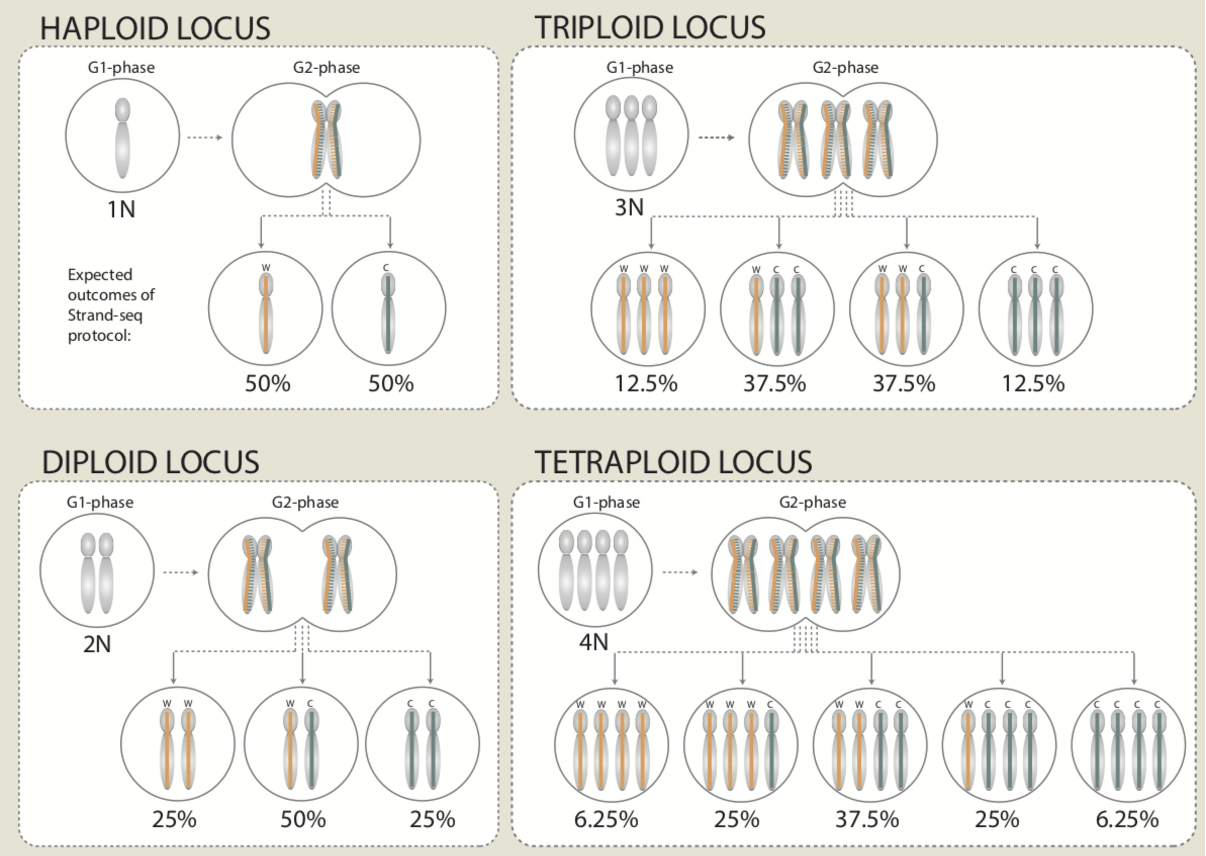


**Figure S2. The origin of diagnostic footprints for ploidy.** Examples are shown for haploidy, diploidy, triploidy, and tetraploidy, respectively. Abbreviations: W, Watson strand; C, Crick strand; G1/2-phase, phases of the cell division cycle; H1 - H4, chromosomal homologs 1 to 4 (with a tetraploid cell, for example, carrying four individual homologs during G1-phase). As part of the Strand-seq protocol incorporation of BrdU into the non-template strand will yield, for each autosome, a pair of sister chromatids (one pair per replicated homolog) with labeled non-template strand during G2-phase. After faithful cell division, daughter cells will yield exactly one chromosomal copy per homolog, each of which has a 50% chance of being inherited as a W or C strand from the mother cell[1,2]. The probability of observing a certain configuration of W and C template strands (referred to as autosomal strand pattern) in Strand-seq data depends on the ploidy state *N*, leading to characteristic expected frequencies for each observed autosomal strand pattern (here indicated as percentages), which can be computed according to a binomial distribution (see **Methods**). This implies that each ploidy state shows a characteristic diagnostic footprints, which enables detection of ploidy alterations in single cells (see **Table S4**).

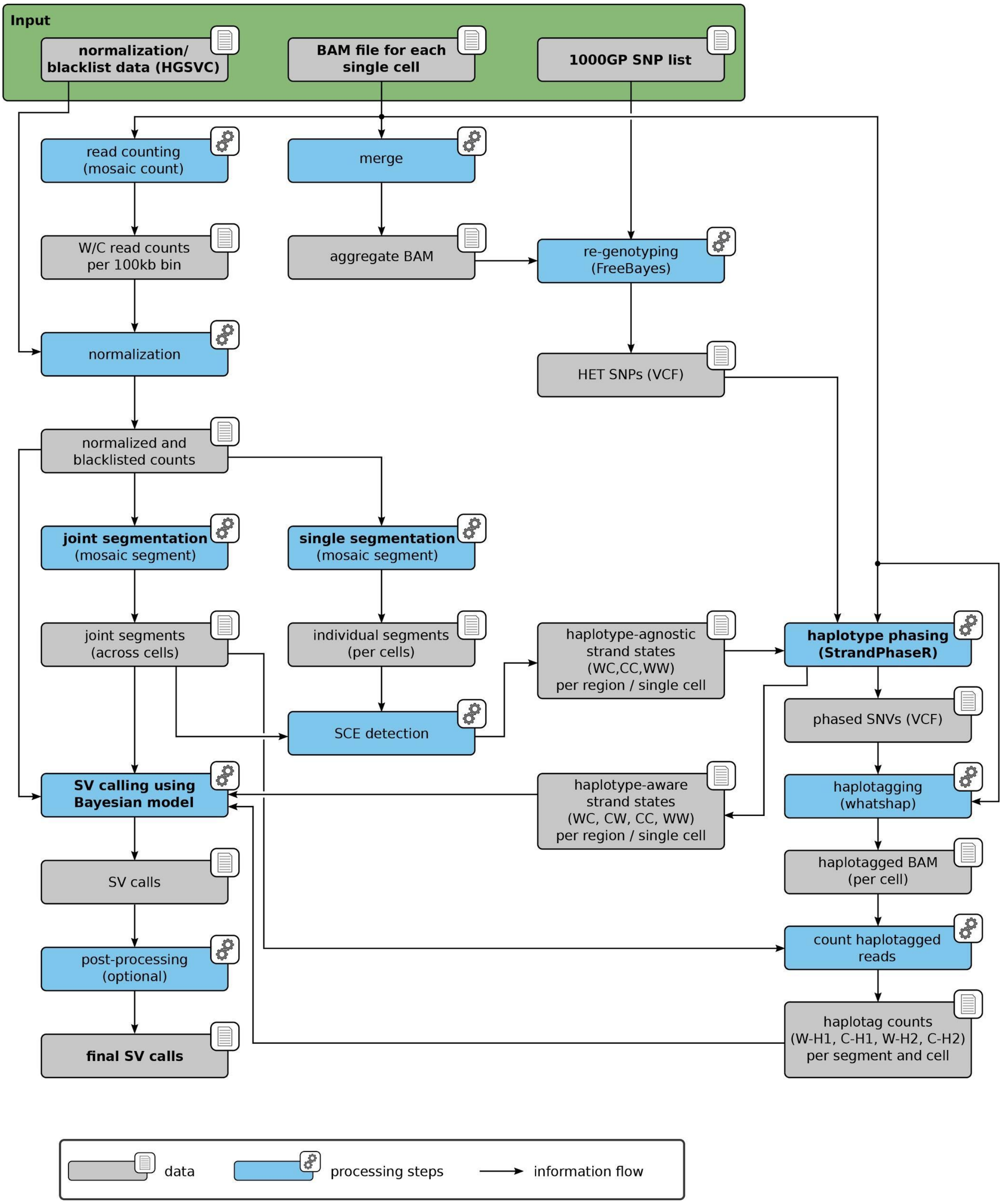


**Figure S3**. Core workflow used for calling SVs in single cells.

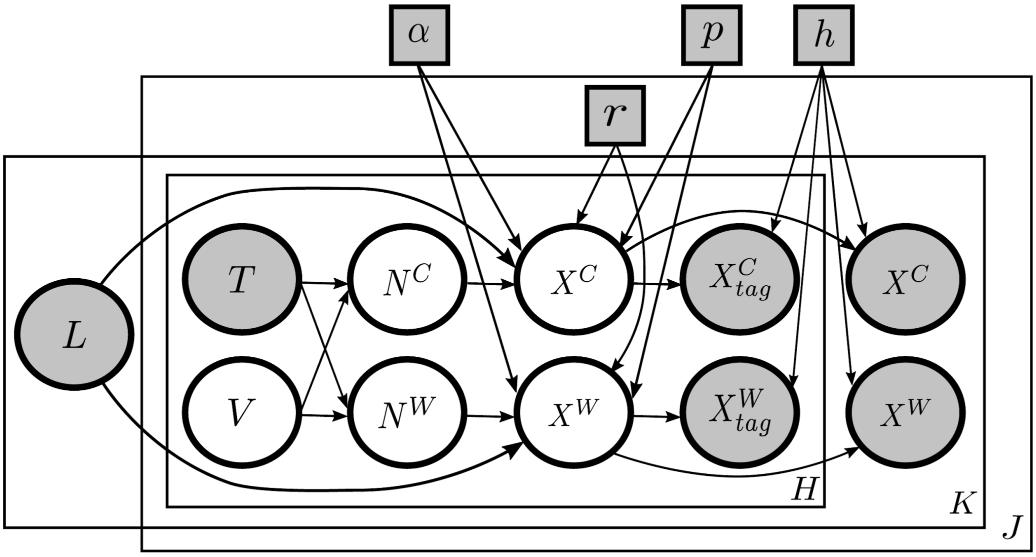


**Figure S4. Bayesian graphical model for haplotype-aware SV classification.** Model shown used to enable haplotype-aware SV discovery in single cells. This graphical model adopts the common plate notation: Circles represent random variables, squares show the model parameters, gray (white) objects show observed (latent) variables, arrows indicate dependencies, and large rectangles indicate that the enclosed variables exists multiple times. The model describes $J$ single cells, $K$ segments, and $H = 2$ haplotypes. Random variables: segment length $L$, ground state $T$, haplotype SV status $V$ (to be inferred), copy numbers of W/C reads $N^{W/C}$, read counts in W/C direction $X^{W/C}$, and read counts in W/C direction tagged by haplotype $X_{tag}^{W/C}$. Note that the read counts are *not observed* by their haplotypes (*white* circles inside the H box), but they are *observed* with no haplotype information (*gray* circles outside the H box). The fraction of reads that overlap with a heterozygous SNP are observed by haplotype (tagged gray read count variables inside the H box). Model parameters: the fraction of background reads $\alpha$, negative Binomial parameter $p$ and $r$, and the heterozygosity rate $h$.

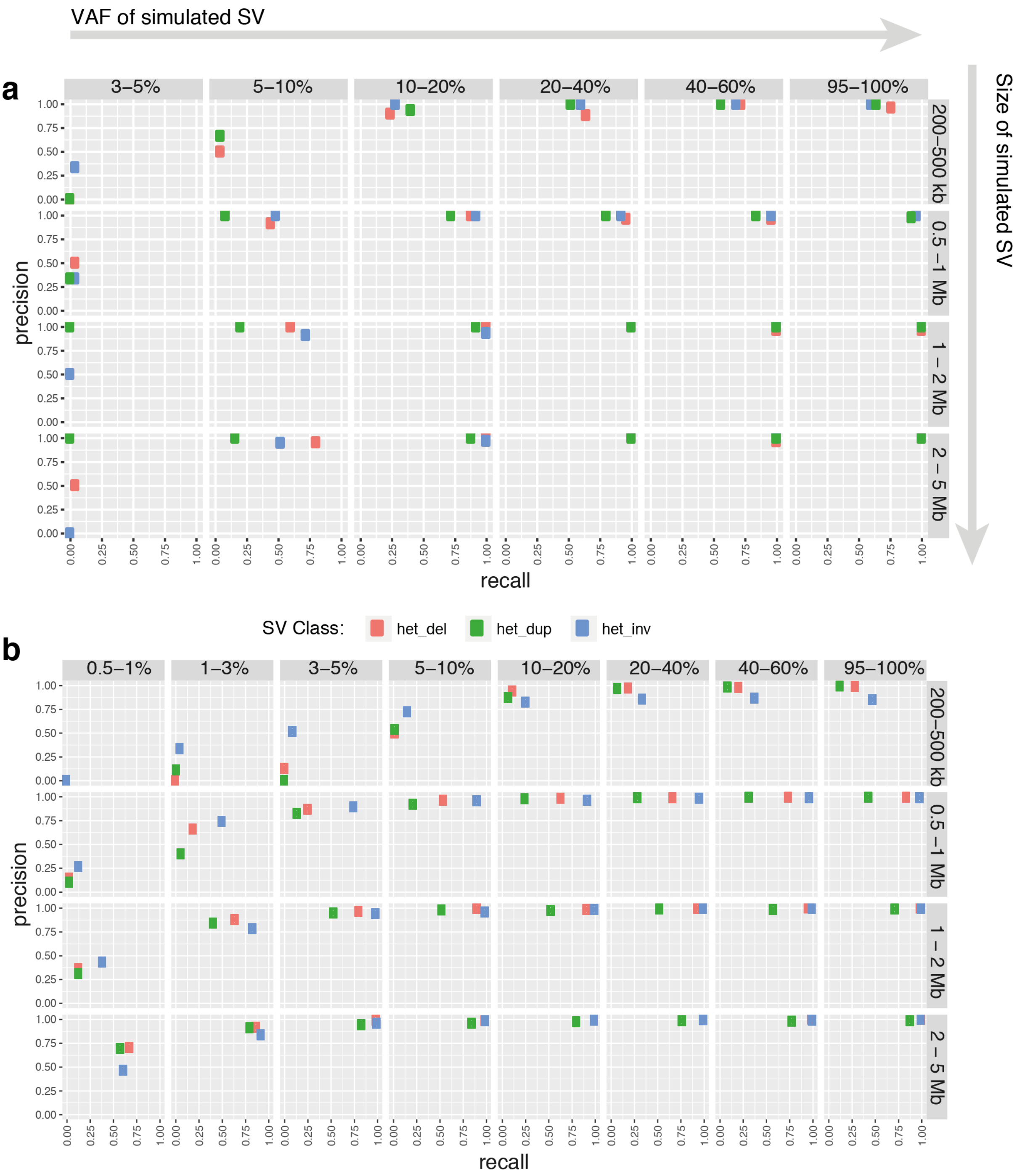


**Figure S5. MosaiCatcher - recall and precision on simulated data.** (**a**) SV calling using the 'strict' parameterization, on simulated data (see Supplementary Experimental Procedures). (**b**) SV calling using the 'lenient' parameterization, on simulated data. Percentage values given indicate simulated somatic variant allele frequencies (VAF) in a given simulation, and rows indicate simulated SV size bins.

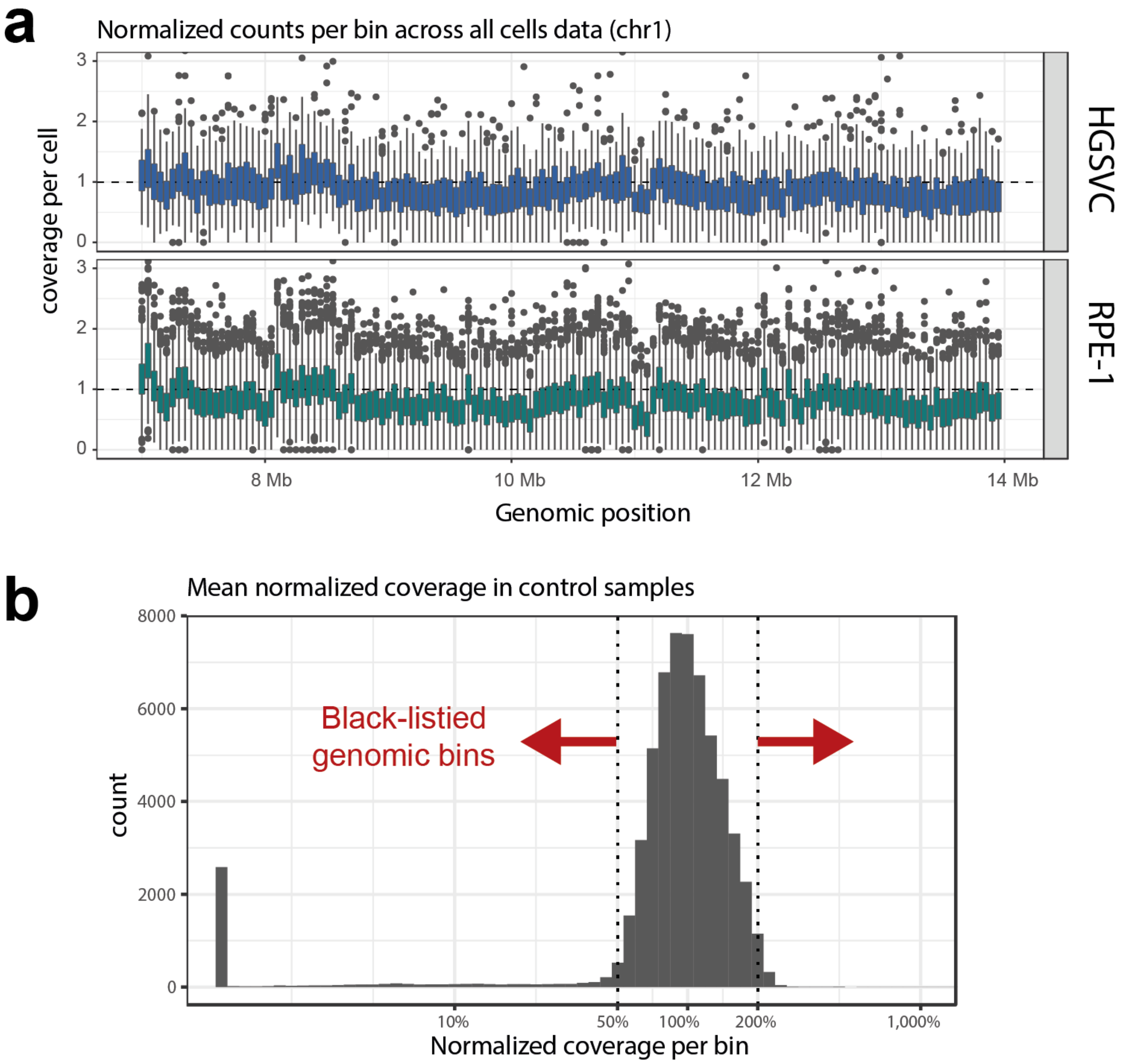


**Figure S6. Normalization of coverage pursued when using MosaiCatcher in conjunction with single cell sequencing data. (A)** Read coverage, summarized in 100 kb windows across sequenced cells, is shown for a region of chromosome 1 (for cumulative read coverage, *i.e.* W + C reads) for HGSVC (Human Genome Structural Variation Consortium) lymphoblastoid cell lines (upper) and RPE-1 cells (lower). **(B)** Bins with a skewed mean coverage in the reference dataset are consistently over or under-represented and were thus blacklisted.

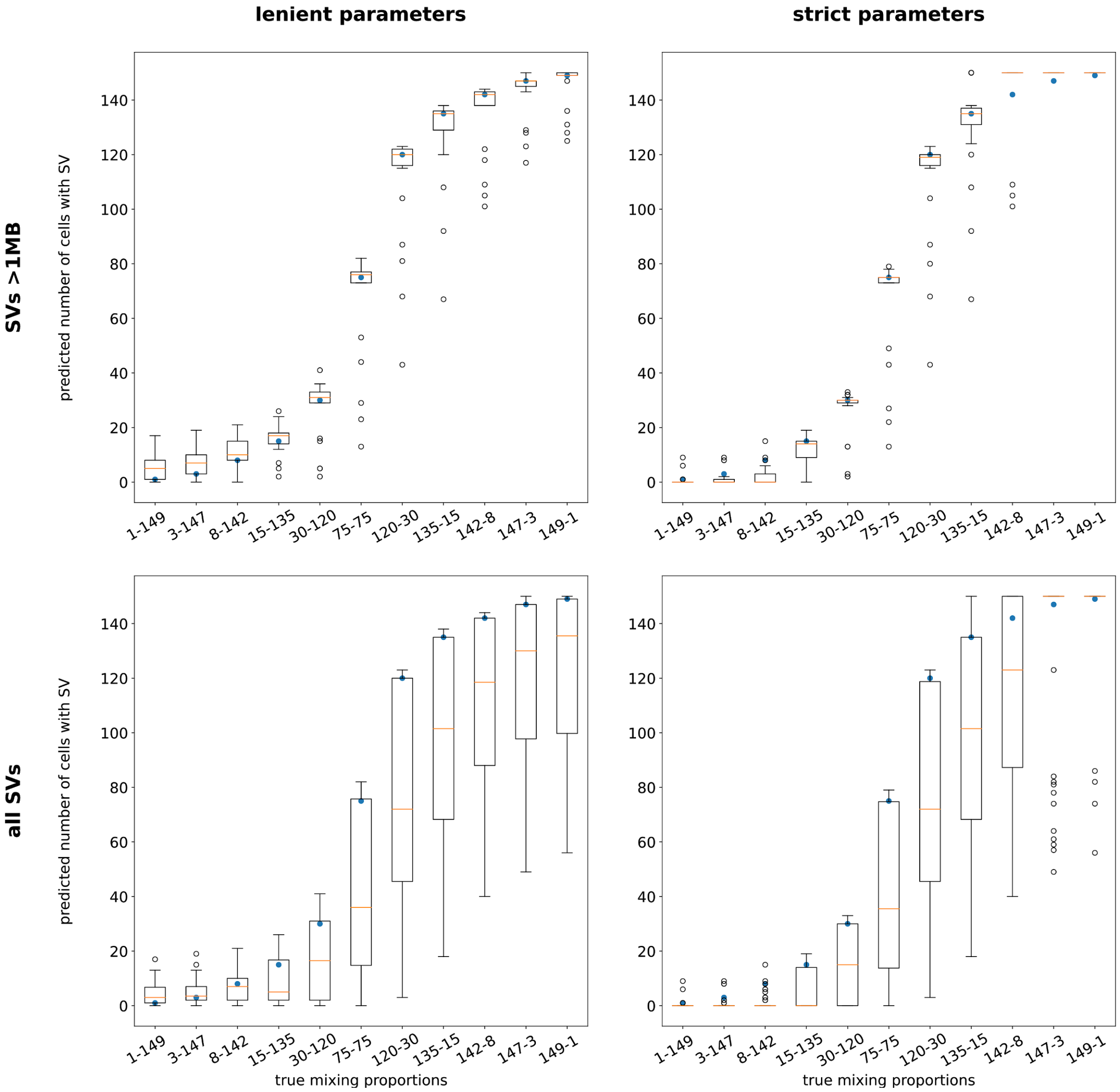


**Figure S7. Evaluation by *in silico* cell mixing.** SV calling performance when mixing single cell libraries from RPE-1 and C7 *in silico*. Each experiment was pursued by randomly sampling 150 single cell libraries with replacement. Eleven different proportions of cell from RPE-1 and C7, ranging from 1-149 to 149-1 were tested and for each proportion, five repetitions were run. We evaluated those SVs that had been independently validated by WGS or mate pair sequencing and called with VAF ≥0.9 in the original scTRIP call sets. For these SVs, boxplots depict the number of cells in which an event was recalled (y-axis) for each true mixing proportion (x-axis). A perfect SV caller would yield results corresponding to the blue circles. Top row: evaluation of SVs >1Mb in size; bottom row: SVs of all sizes; left column: lenient parameterization; right column: strict parameterization.

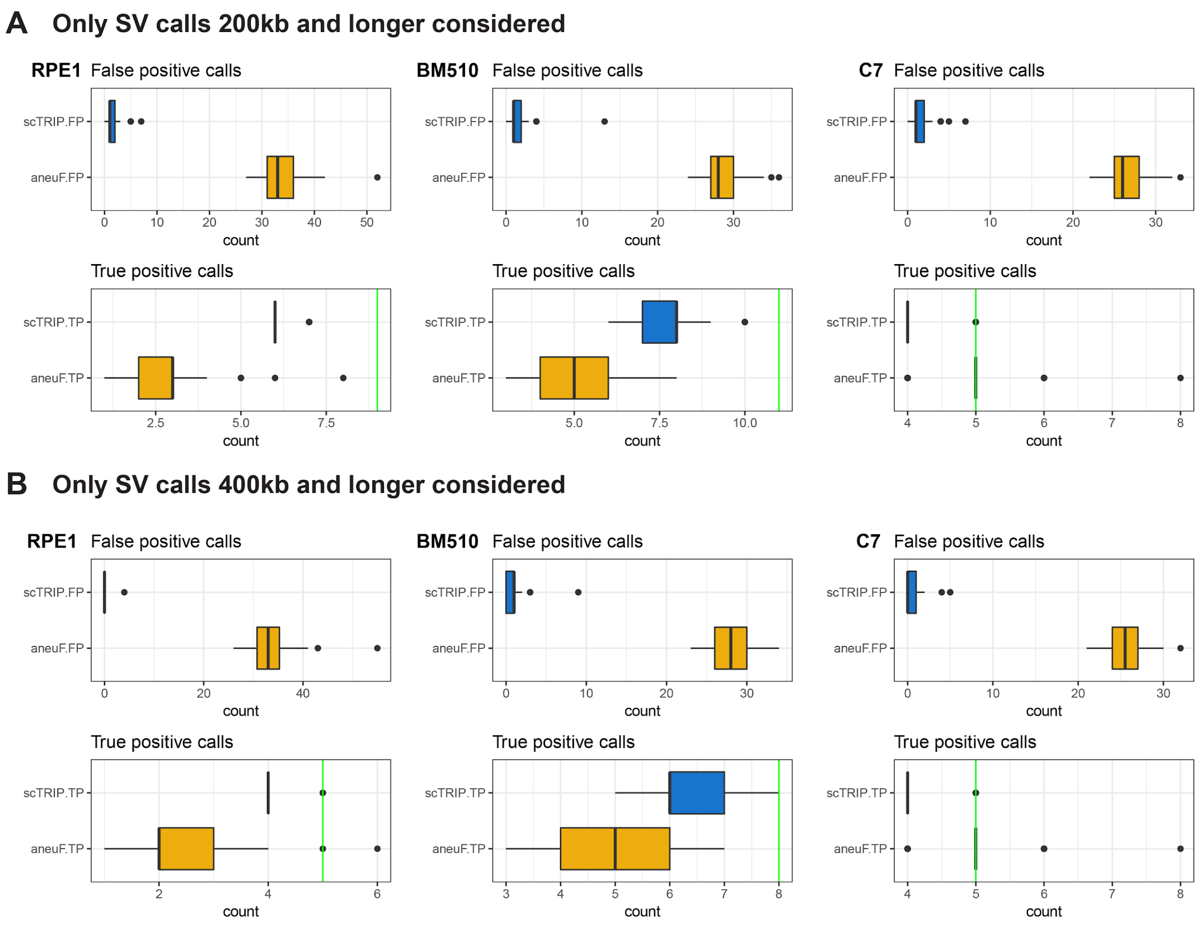


**Figure S8. Comparison of scTRIP with a single cell CNA detection method (AneuFinder).** CNAs (deletions and duplications) discovered separately by scTRIP (blue) and AneuFinder (yellow) were compared against a set of presumed true positive, clonal CNAs (**Table S5**) discovered by using bulk WGS or using mate-pairs (the 'ground truth set'). Boxplots show the distribution of false positive (FP) and true positive (TP) CNA calls across single cells. In (**A)** we considered all CNA calls, while in **B** we considered CNA calls ≥400kb (**Table S3**). The total number of true positive calls we compared against is marked by green vertical lines separately for each tested cell line. In some cases owing to 'over-segmentation', AneuFinder generated a larger number of calls overlapping with the ground truth set, which were scored as true positives. SV calls made by scTRIP did not show such over-segmentation. Even though the analysis shown in this figure focused only on CNAs, scTRIP generated more true positive SV calls and less false positive SV calls than the single cell CNA detection tool.

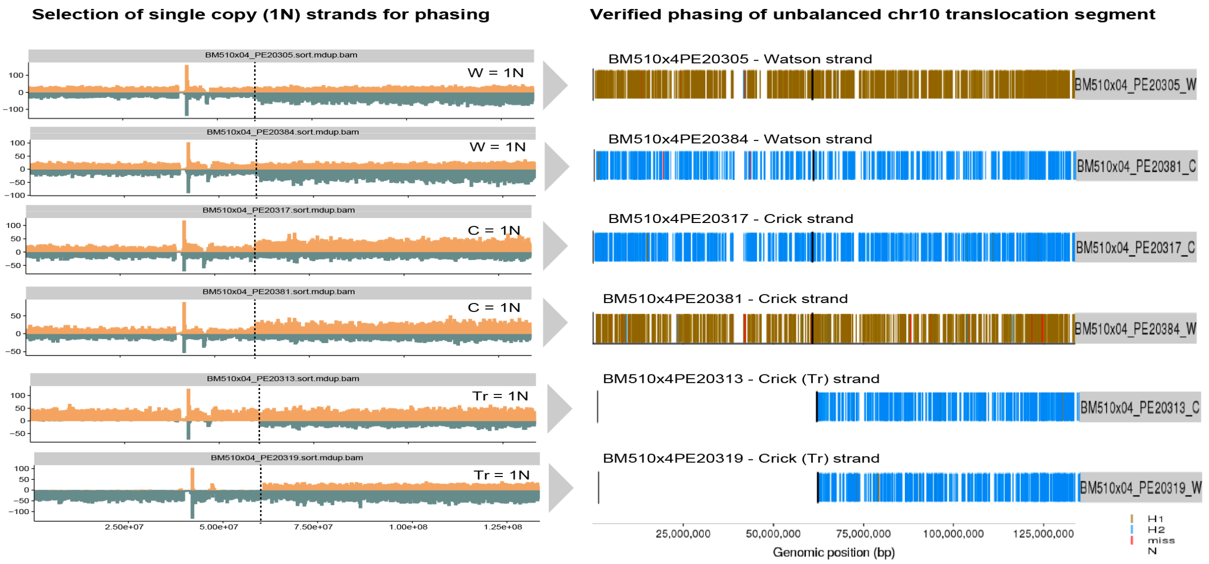


**Figure S9. Phasing of unbalanced translocation using BM510 single cell data.** Six BM510 cells are shown carrying the unbalanced t(X;10) translocation. Binned read depth data for chromosome 10 are depicted on the left column (W reads - orange, C reads - green). A dashed vertical line demarks the 10q translocation breakpoint. For each cell the strand represented as a single copy along the whole length (i.e. not containing the 10q gain) is listed as 1N. This strand was used for unambiguous phasing, to overcome any haplotype mixing that can occur at triploid regions. Phasing outcomes of the single copy strands are shown to the right. Vertical lines depict SNPs assigned to either haplotype 1 (H1, brown) or haplotype 2 (H2, blue). The shorter duplicated haplotype in the lower two cells (labeled with Tr) corresponds to H2, confirming the H2 haplotype of the translocated 10q segment. Tr, translocation.

**A**

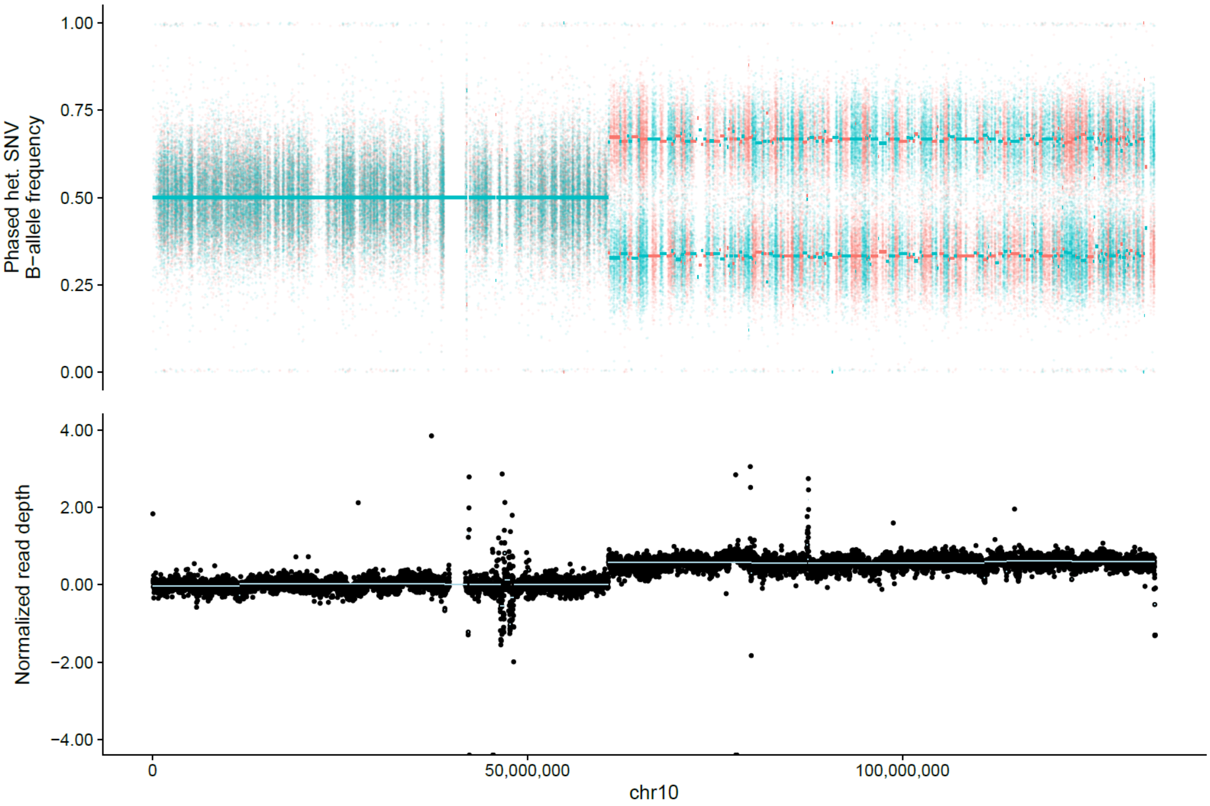


**B**

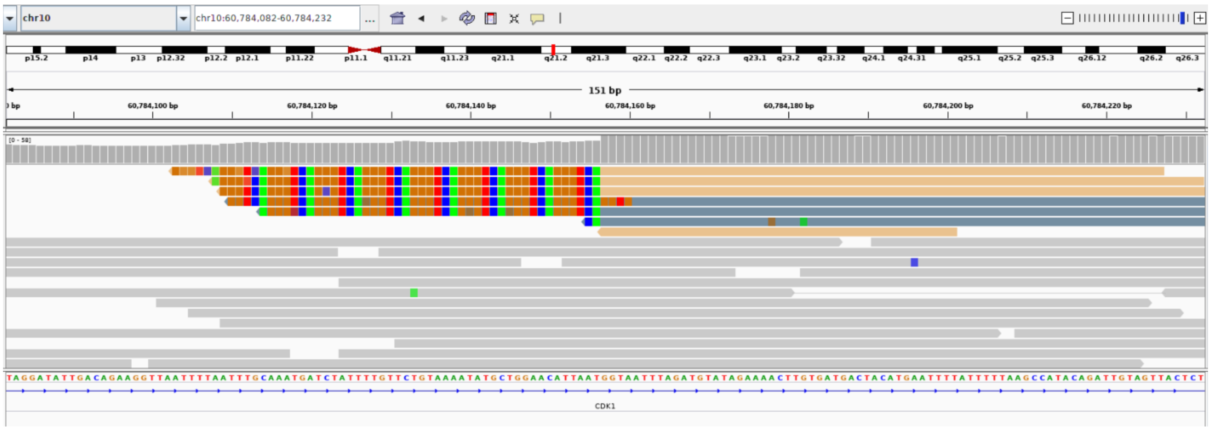


**Figure S10. RPE-1 unbalanced translocation escapes detection in bulk WGS data.** (**A**) Read depth plot of the affected region on chromosome 10, that in RPE-1 cells is fused to the end of chromosome X as shown by spectral karyotyping and detectable by three-channel processing (see *e.g.* **Fig. 3**). (**B**) IGV[3] plot with split reads centered at the chromosome 10 breakpoint of this unbalanced translocation. The other breakpoint junction of this SV maps to highly repetitive DNA, and thus remains unresolved by bulk Illumina bulk WGS, with this breakpoint junction lacking uniquely aligned reads that map to chromosome X.

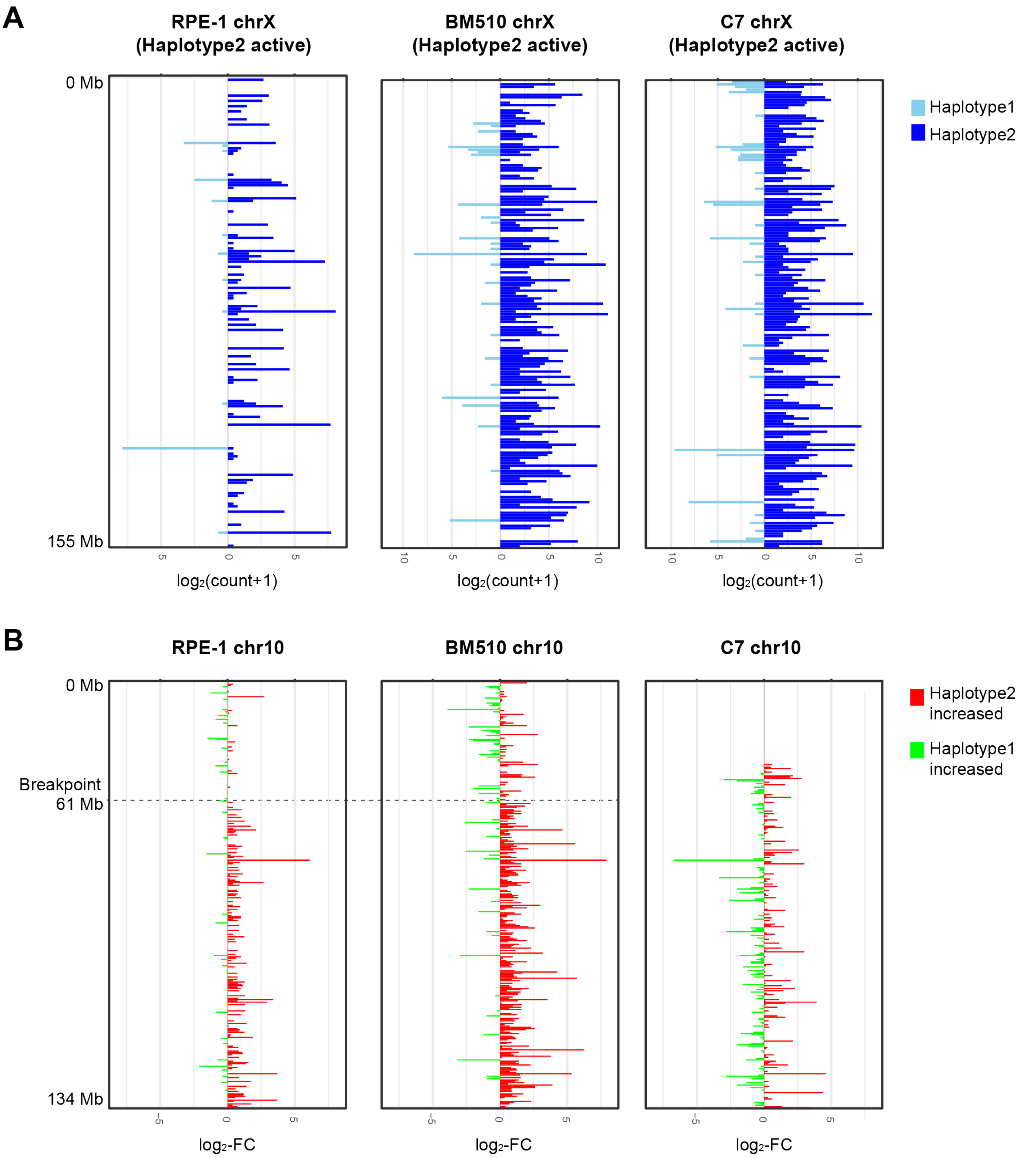


**Figure S11. Allele-specific expression analysis reveals involvement of the active X in the t(X;10) unbalanced translocation.** Allelic RNA expression analysis of chromosomes X and 10 in RPE cells pursued to investigate the consequences of an unbalanced translocation involving chromosomes 10 and X present in RPE-1 and BM510. scTRIP generates haplotype-resolved SV calls, whereby the two haplotypes (homologs) existing for each chromosome in a diploid cell are arbitrarily named H1 and H2 (for haplotype/homolog 1 and haplotype/homolog 2, respectively). In this figure, 'Haplotype 1' of RPE-1, BM510 and C7 (and also 'Haplotype 2') exhibit the same genotype (and hence correspond to the same chromosome homolog). Haplotype 2 is involved in der(X) t(X;10) translocation in RPE-1 and BM510. (**A**) Allelic read count plots of chromosome X based on scTRIP haplotype information and RNA-seq data of the corresponding RPE cells, which shows involvement of the active (rather than the inactive) X chromosome in the t(10,X) translocation. (**B**) Fold change plots of chromosome 10 comparing allelic read count of H1 and H2. RNA expression on the gained and translocated haplotype is, as expected, increased compared to the non-translocated haplotype in the context of the unbalanced t(10,X) translocation - further corroborating scTRIP's haplotype assignments. Each bar represents a gene; genes are sorted by genomic position.

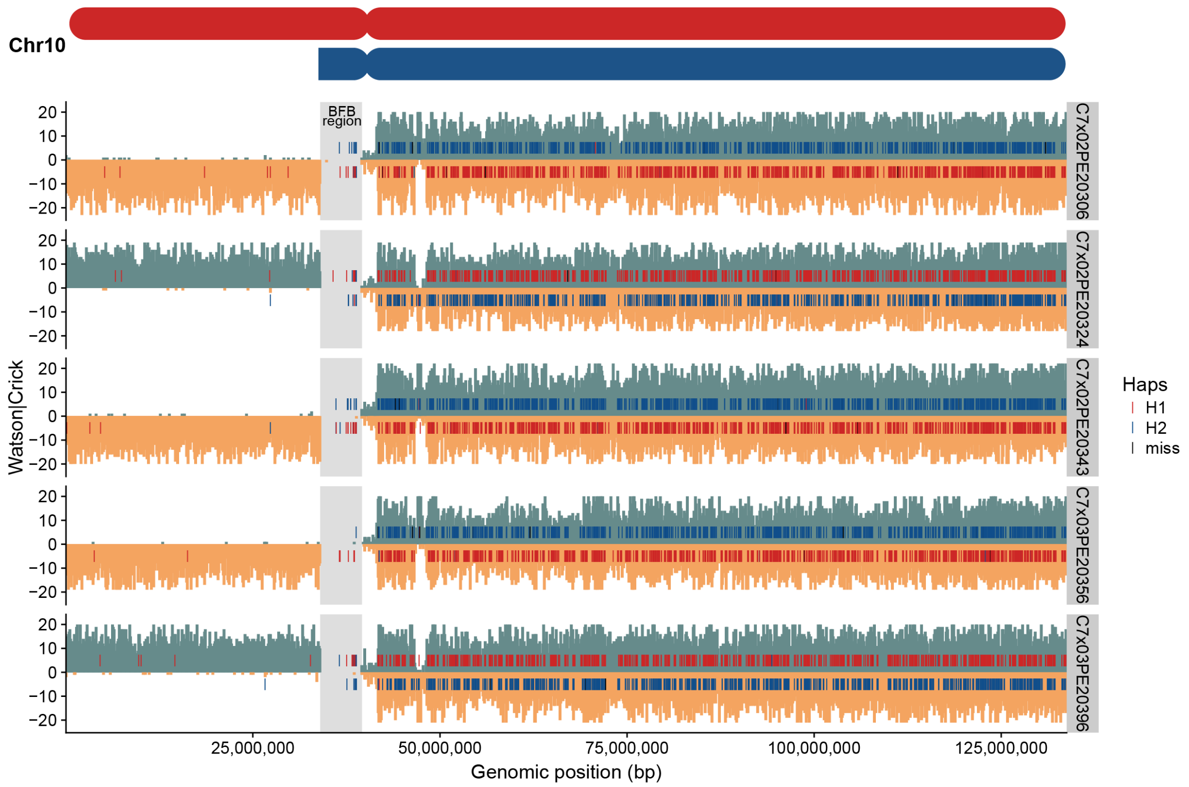


**Figure S12. Phasing of a homologue inferred to have undergone breakage-fusion-breakage cycles (BFB).** DelTer rearrangements seen at 10p in C7 were in each case located on the same haplotype, including cells with and without the 10p amplicon inferred to be formed by BFB cycles. To illustrate the haplotype analyses pursued we picked 5 cells randomly (shown above). For each cell we plotted reads mapping to the reference genome in plus directionality (C - green) above the zero line and those that map in minus directionality (W - orange) below the zero line. In each single cell, heterozygous SNPs represented separately in the C and W portion of reads were compared to the consensus haplotypes assembled from all haplotype informative cells. SNPs that matched haplotype 1 (H1) are colored in red and those that match haplotype 2 (H2) are colored in blue. SNPs that do not match any of the consensus haplotypes ('miss') are colored in black. Grey rectangles highlight the inferred BFB amplicon region (reads that map into this 10p segment have been removed from this plot, to enhance overall visibility). The pattern observed can be explained by the DelTer events always being present on the same haplotype (H2 - blue) in all C7 cells, including cells with the inferred BFB amplicon and cells without the inferred BFB amplicon. Haps, haplotypes.

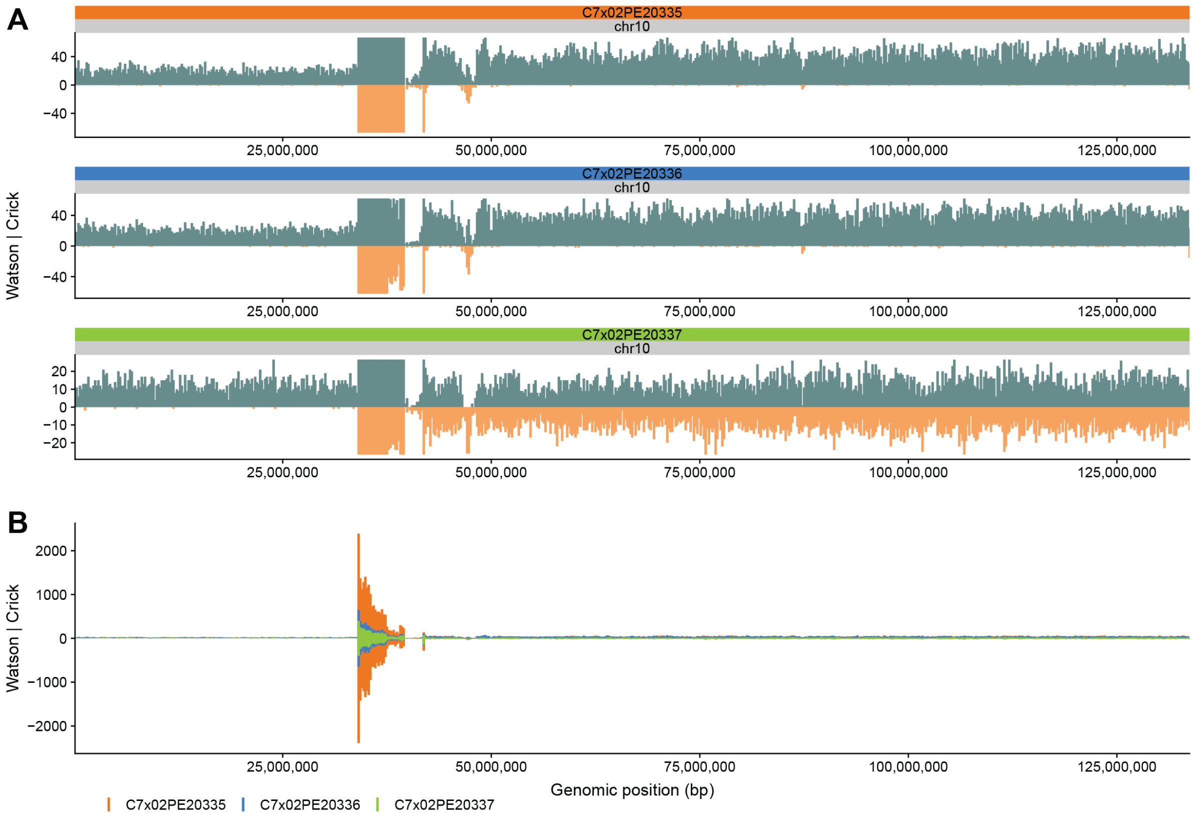


**Figure S13. C7 single cells with intermediate and high copy-number at 10p plotted using different scales.** Cell 335, showing excessive copy-number at 10p in C7 compared to other C7 cells, is depicted along with two cells showing intermediate copy-number at that 10p chromosomal location. Marked copy-number increase in cell 335 (cell id: C7x02PE20335; inferred copy number of 400-500) is only detectable for the BFB-associated amplicon region - *i.e.* is specific to the amplicon. The q-arm of cell 335 retains disomy, and the terminal segment of 10p is monosomic in this cell (DelTer) - as seen for all other C7 single cells exhibiting the BFB-associated amplicon. (**A**) Coverage of selected cells with BFB is shown in a scale that allows to focus on the disomic and monosomic regions. (**B**) The same cells are shown at constant linear scale, whereby this time the scale was chosen to visualize differences in 10p amplicon height.

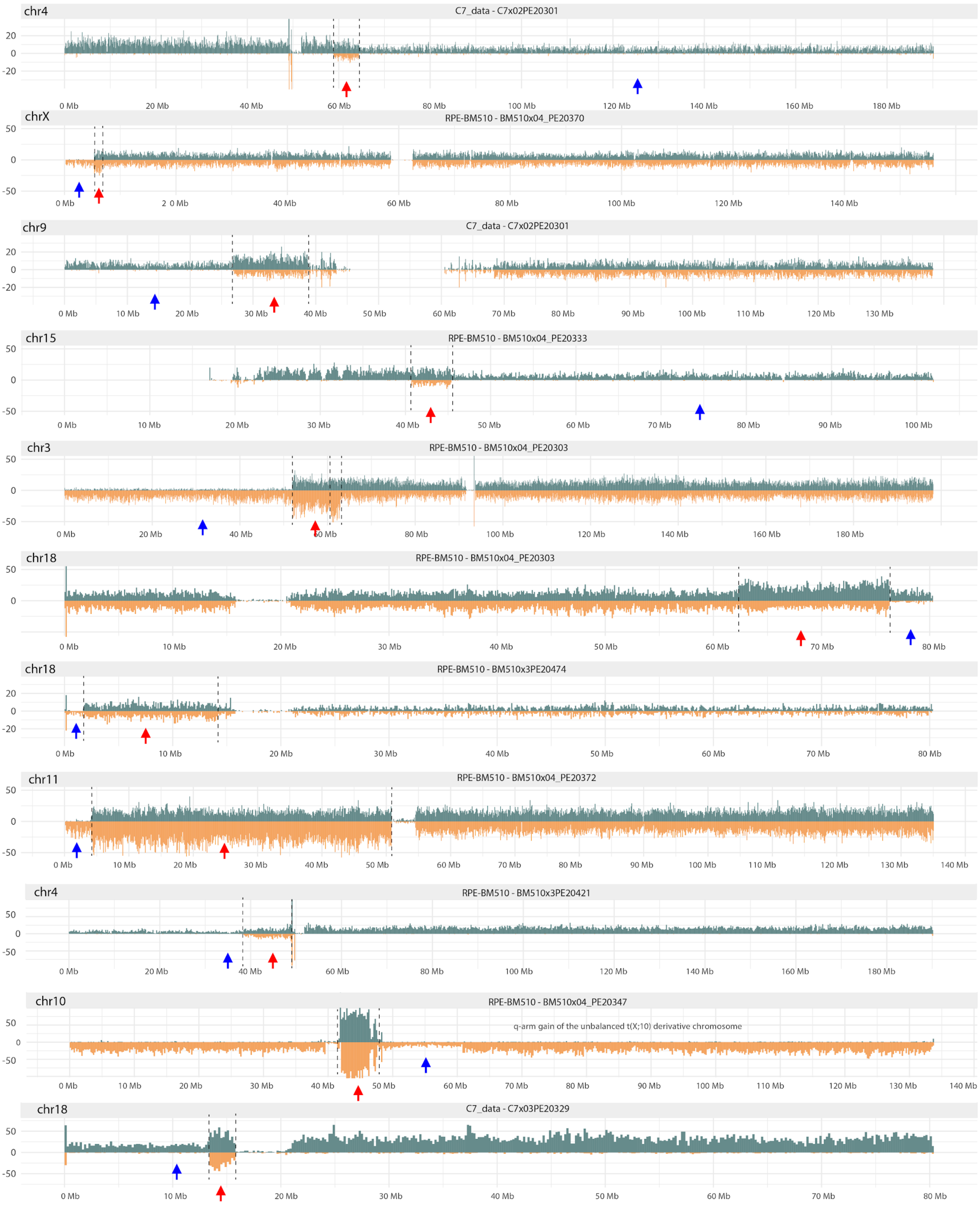


**Figure S14. Clustered rearrangements in single cells from C7 and BM510.** Whole chromosome plots highlighting individual Breakage-fusion-bridge (BFB) events located in transformed RPE cells. For each event, InvDup rearrangements (red arrows) are immediately flanked by terminal chromosome segment deletions (DelTer, blue arrows) arising on a single haplotype. Note this display item shows all cells having a 'classic' BFB signature with no other SVs present on the same homolog. Homologs containing BFB events along with additional SVs (*e.g.* deletions) are depicted in **Fig. S15**. Binned W reads are shown in orange (below each chromosome ideogram); binned C reads are show in green (above)

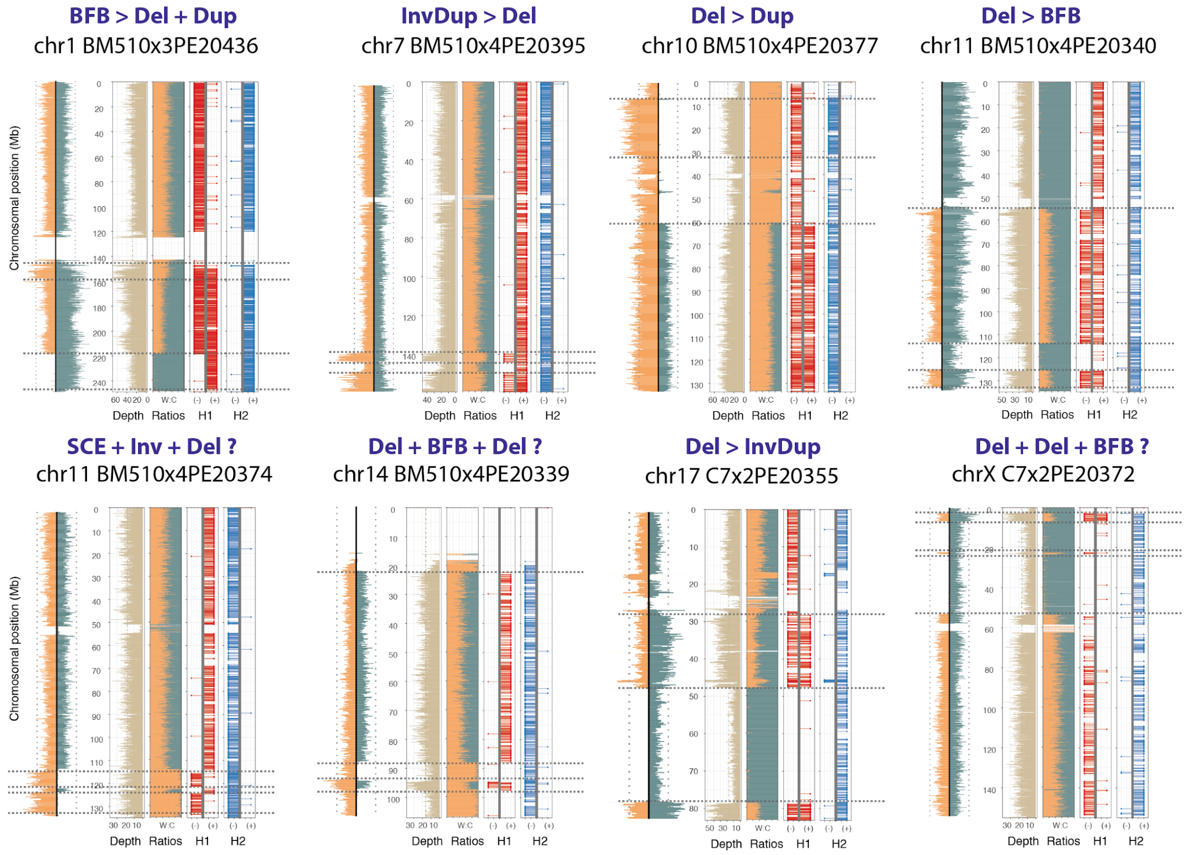


**Figure S15. Single cell based mapping of haplotype-resolved complex/clustered SVs in RPE cells.** Single cell plots of chromosomes identified as having multiple SVs clustered on a single haplotype. For each example the template strand plots are shown to the left, with the three-channels analyzed by scTRIP illustrated on the right. Depth is shown as total binned read counts. Strand ratios are shown as the fraction of W:C reads in each bin. Haplotype phasing data is separated into H1 and H2 with orientation of each phased SNP (shown as a lollipop) indicated by placement on ideogram (SNPs in W reads are shown on the left, SNPs on C reads are on right). Breakpoints located in each cell are indicated by dotted lines. For these clustered SVs we employed the infinite sites assumption[4] to infer a plausible temporal ordering of formed SVs, with the predicted ordering shown above each plot. A ">" symbol is used when one event was estimated to proceed the next, whereas an "+" symbol indicates the temporal sequence was unclear. For instance, the clustered SVs located on chr1 of BM510x3PE20436 (top left cell) was inferred to have undergone a BFB event *followed* by Del and Dup (denoted 'BFB > Del + Dup'), given that we observed CN steps of size 1 (see **Methods**). This was distinguished from chr11 of BM510x4PE20340 (top right cell) where the Del was seen in conjunction with a CN step size of 2, and was hence inferred to occur *prior* to BFB formation (denoted 'Del > BFB'). The SV pattern here suggests the Del first caused a complete loss of a segment on the H1 haplotype. H1 subsequently underwent a BFB cycle forming an InvDup with an interstitial loss (coinciding with the Del event). A lost H1 segment can obviously not be "regained" via subsequent inverted duplication[4], and the subsequent InvDup hence resulted in a CN step size of 2 (see **Methods**). In one case we predicted a sister chromatid exchange event (SCE) to be included in the ordering, as this represented the most parsimonious explanation for the SV profile observed. Use of '?' indicates uncertainty about the temporal ordering of events in a few cells.

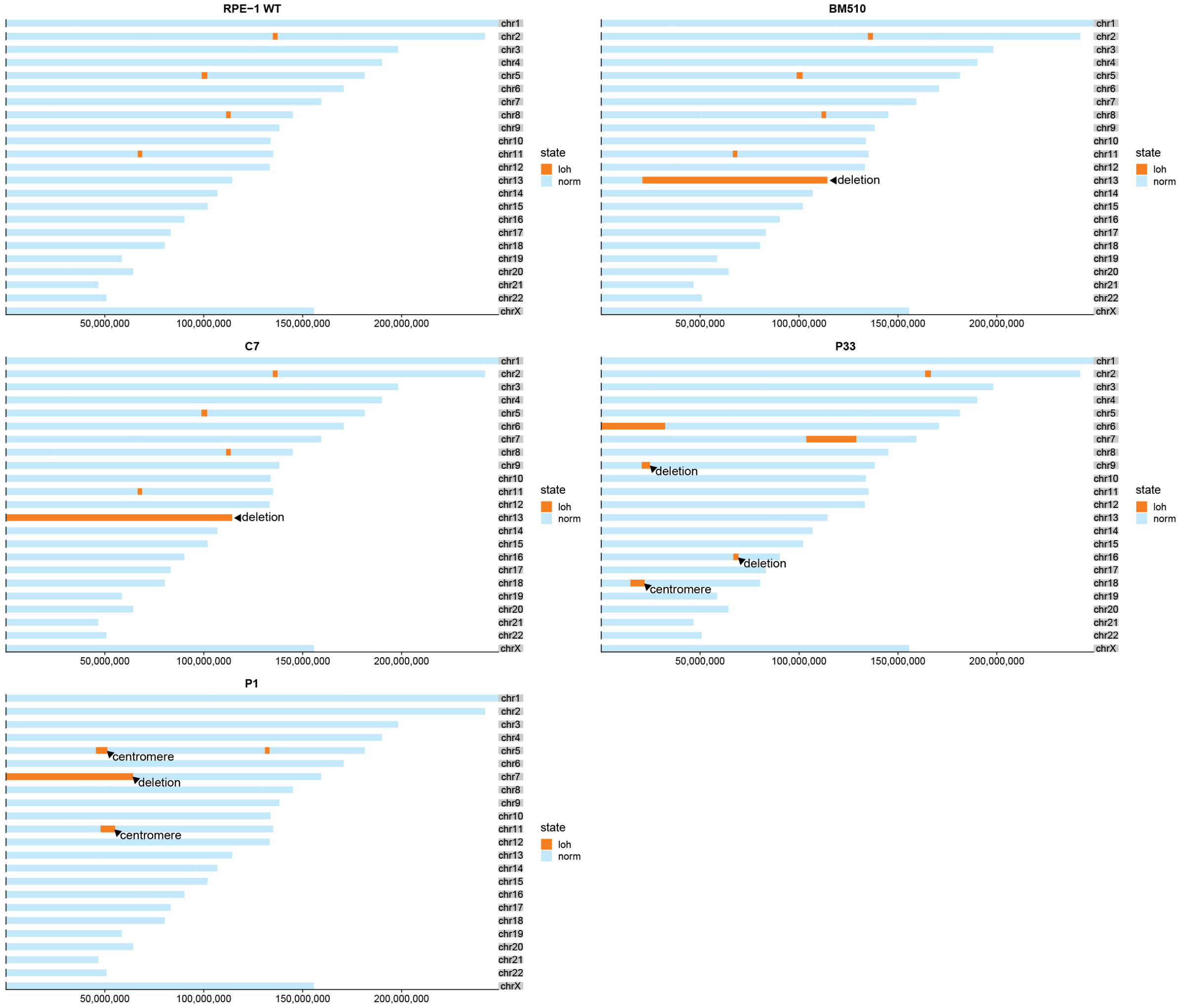


**Figure S16. Detection of regions with loss of heterozygosity.** Genome-wide ideograms showing regions of loss in heterozygosity (LOH) across the samples of our study (RPE-1, BM510, C7, P33 and P1). LOH regions depicted with orange color. Norm, normal.

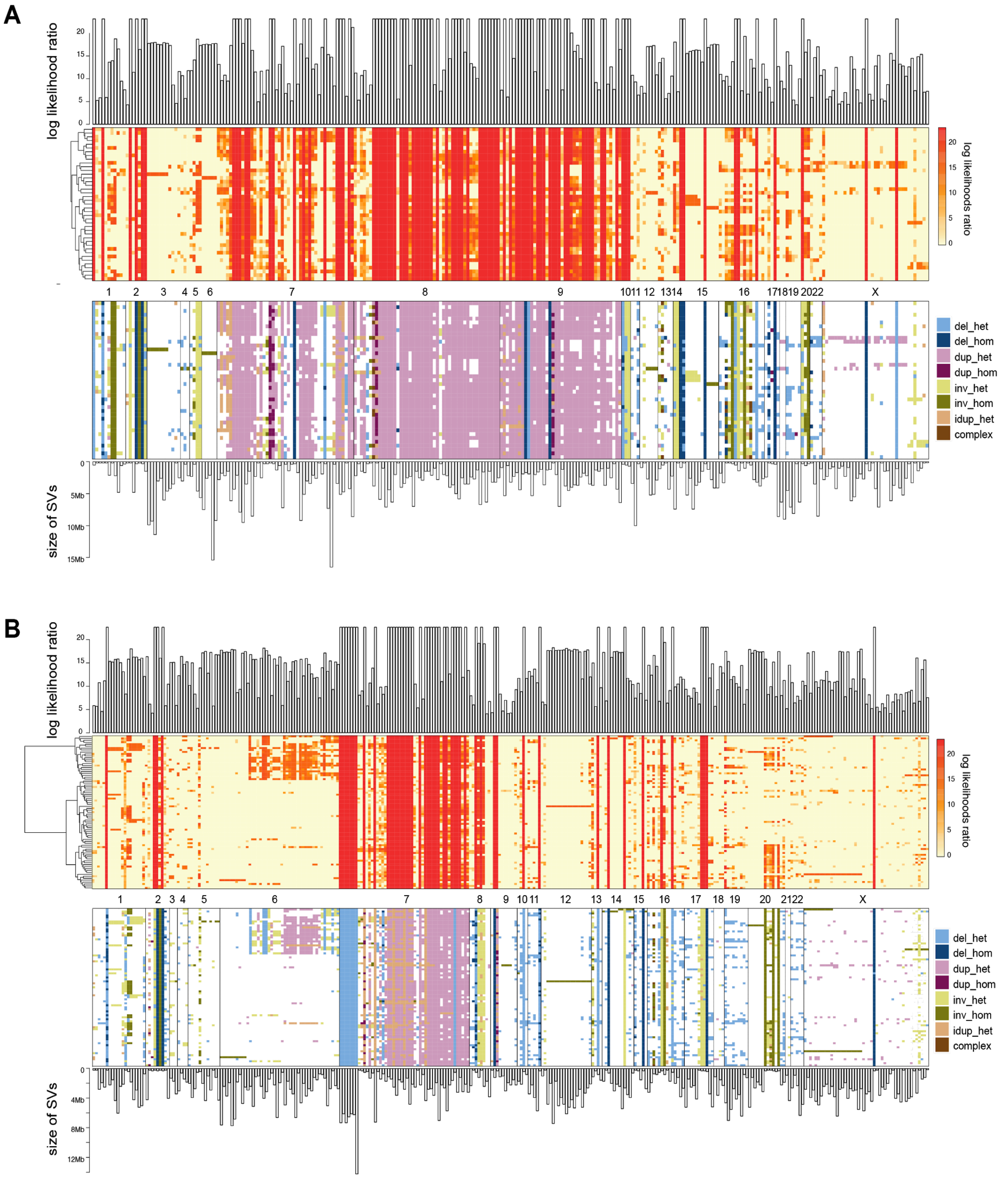


**Figure S17. Leukemic single cell SV landscapes in P33 and P1.** Each column corresponds to scTRIP based SV calls, and each row to single cells. A yellow/red color scale (upper heatmaps) depicts SV genotype likelihoods, and the heatmap below indicates the various SV classes contributing to these SV landscapes. Homozygous inversions can be unambiguously detected on WW or CC chromosomes, but are not readily distinguishable from the reference state in WC or CW chromosomes (**Table S1**). Thus, if more than 40% of the cells in a sample showed homozygous inversions (inv_hom) calls in certain column, other cells in the same column were imputed to contain the same homozygous inversion if in a WC or CW ground state (imputed SVs shown in light green color). Based on the same rationale, we also imputated translocations into each individual cell. Bar graphs above the heatmaps indicates the mean log likelihood ratio (computed across all single cells, for a certain SV); bar graphs below show SV size. Heatmaps were generated using Ward's hierarchical clustering of genotype likelihoods to arrange single cells based on SV call patterns from P33 **(A)** and P1 **(B)**. Hom, homozygous. Het, heterozygous.

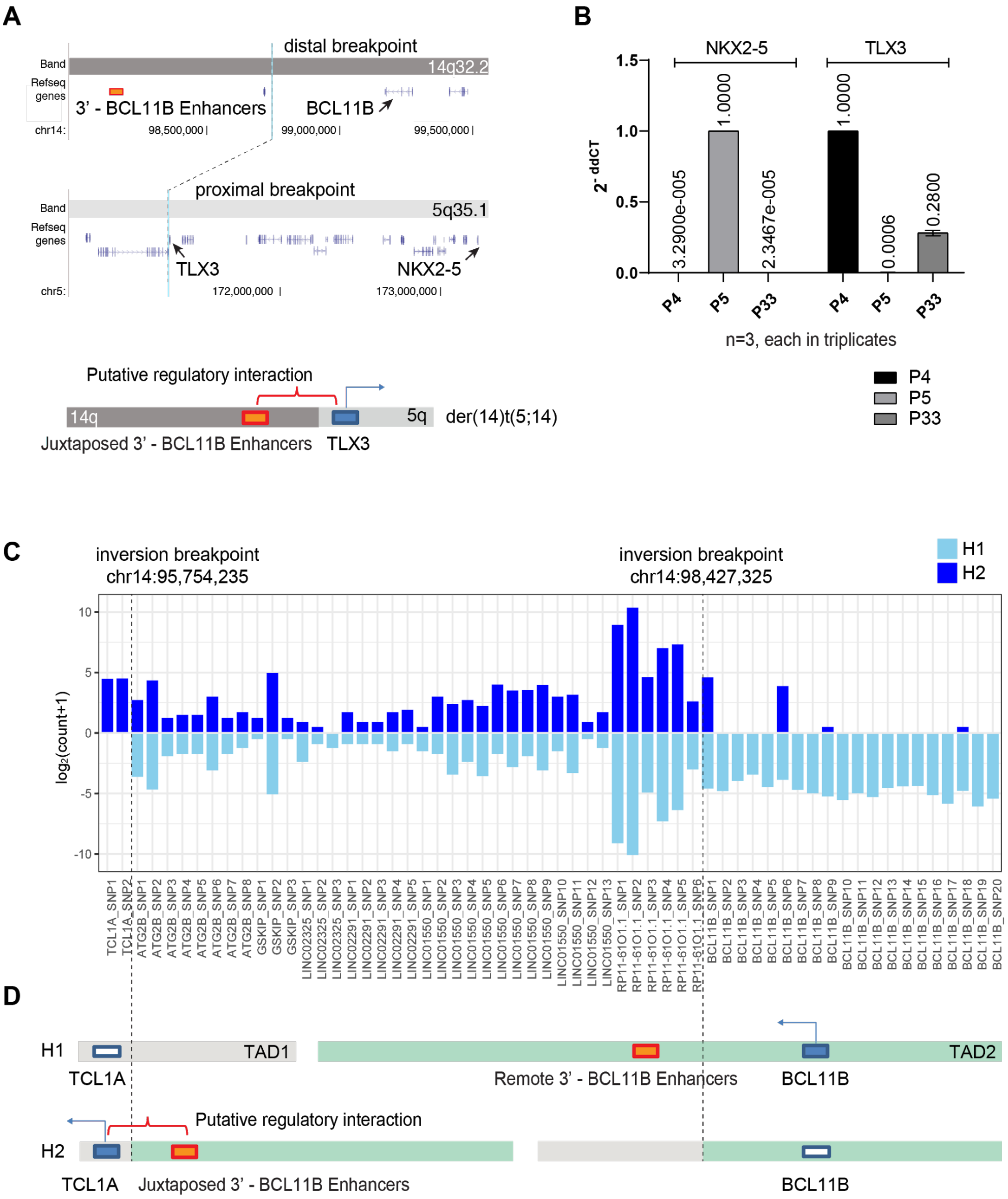


**Figure S18. T-ALL oncogenic dysregulation in conjunction with balanced SVs in 3' of *BCL11B*.** (**A**) The t(5;14) translocation in P33 brings the *TLX3* at chr5q35.1 and enhancers in 3' of *BCL11B* at chr14q32.2 into proximity. (**B**) Quantitative real time PCR (qPCR) validation of *TLX3* dysregulation, and comparison to control samples with high and low *TLX3* levels. (**C**) The plot depicts allelic read counts of the RNA-seq (P1) detected from heterozygous germline SNPs within gene loci residing in two previously reported topologically associating domains (TADs)[5], both of which are affected by the inversion identified in P1. (**D**) Rearrangements of TADs in the context of the 14q32 inversion in H2. Following 14q32 inversion, the *TCL1A* and nearby enhancers are not separated by a TAD boundary any more, which may have facilitated long-range regulatory interactions between the loci involved[6,7]. Allele-specific expression measurements revealed significantly increased expression of the *TCL1A* H2 allele compared to the H1 allele (FDR = *6.68E-21*). The red box indicates the genomic position of remote 3'-*BCL11B* enhancer elements thought to mediate oncogene overexpression[8]. TAD boundary information was obtained from the literature[5]. The dashed lines depict the inversion breakpoints, which we verified by mate-pair sequencing of the P1 sample[9].

### Supplementary Tables

**Table S1.** ***Overview of SV diagnostic footprints for different strand state inheritance patterns***
The diagnostic footprints described by scTRIP consider three data layers - read depth, read orientation and phase. Classifying SVs using scTRIP depend on the underlying strand state of a chromosome, and whether the SV is homozygous or heterozygous. The below Table introduces this principle for WC, CW, WW and CC ground states. While SV classes occasionally can not be unambiguously called in a single cell alone (*, e.g. homozygous inversions), all SVs can be unambiguously resolved when examining subclonal SVs at the cell population level.

[1] Cannot be distinguished from a reference state in WC/CW chromosomes* (yet is resolved in CC and WW chromosomes, when assessing subclonal SVs in a cell population, or when using haplotags)

[2] Cannot be distinguished from a heterozygous duplication in WC/CW chromosomes* (yet is resolved in CC and WW chromosomes, when assessing subclonal SVs in a cell population, or using haplotags)

[3] Cannot be phased in WW or CC chromosomes* (yet is resolved for WC/CW chromosomes, when assessing subclonal SVs in a cell population, or when using haplotags)

**SV diagnostic footprints in a WC chromosome**

| SV state | Coverage | W. frac. | Haplotype tags W | Haplotype tags C | W cov | C cov |
|---|---|---|---|---|---|---|
| Reference state | 2N | 50% | H1 | H2 | 1N | 1N |
| Deletion of H1 | 1N | 0% | - | H2 | 0N | 1N |
| Deletion (homozygous) | 0N | - | - | - | 0N | 0N |
| Duplication of H1 | 3N | 66% | 2xH1 | H2 | 2N | 1N |
| Duplication (homozygous) | 4N | 50% | 2xH1 | 2xH2 | 2N | 2N |
| Inversion of H1 | 2N | 0% | - | H1+H2 | 0N | 2N |
| Inversion (homozygous)[1] | 2N | 50% | H2 | H1 | 1N | 1N |
| Inverted duplication of H1[2] | 3N | 33% | H1 | H1+H2 | 1N | 2N |

**SV diagnostic footprints in a WW chromosome**

| SV state | Coverage | W. frac. | Haplotype tags W | Haplotype tags C | W cov | C cov |
|---|---|---|---|---|---|---|
| Reference state | 2N | 100% | H1+H2 | - | 2N | 0N |
| Deletion of H1[3] | 1N | 100% | H2 | - | 1N | 0N |
| Deletion (homozygous) | 0N | - | - | - | 0N | 0N |
| Duplication of H1[3] | 3N | 100% | 2xH1+H2 | - | 3N | 0N |
| Duplication (homozygous) | 4N | 100% | 2xH1+2xH2 | - | 4N | 0N |
| Inversion of H1[3] | 2N | 50% | H2 | H1 | 1N | 1N |
| Inversion (homozygous) | 2N | 0% | - | H1+H2 | 0N | 2N |
| Inverted duplication of H1[3] | 3N | 67% | H1+H2 | H1 | 2N | 1N |

**SV diagnostic footprints in a CC chromosome**

| SV state | Coverage | W. frac. | Haplotype tags W | Haplotype tags C | W cov | C cov |
|---|---|---|---|---|---|---|
| Reference state | 2N | 0% | - | H1+H2 | 0N | 2N |
| Deletion of H1[3] | 1N | 0% | - | H2 | 0N | 1N |
| Deletion (homozygous) | 0N | - | - | - | 0N | 0N |
| Duplication of H1[3] | 3N | 0% | - | 2xH1+H2 | 0N | 3N |
| Duplication (homozygous) | 4N | 0% | - | 2xH1+2xH2 | 0N | 4N |
| Inversion of H1[3] | 2N | 50% | H2 | H1 | 1N | 1N |
| Inversion (homozygous) | 2N | 100% | H1+H2 | - | 2N | 0N |
| Inverted duplication of H1[3] | 3N | 33% | H1 | H1+H2 | 1N | 2N |

**Table S2. Overview of Strand-seq libraries included in the study**
Metrics of the single cell sequencing data for RPE-1, C7, BM510, P33 and P1 samples, with total number of high-quality mapped fragments per library listed.
*This table is provided as an external data file.*

**Table S3. SV calls generated with our framework and using external methodologies**
Overview of the single cell SV calls generated for RPE-1, C7, BM510, P33 and P1 samples, with variant allele frequencies and orthogonal validation notes included
*This table is provided as an external data file.*

**Table S4. Diagnostic footprints characteristic for ploidy states**.
A binomial distribution can be used to compute expected frequencies of autosomal strand patterns for alternative ploidy states. W, Watson. C, Crick.

| **Ploidy state** | **Strand patterns observed** | | | | |
|---|---|---|---|---|---|
| **Haploid** | C | W | | | |
| Strand-ratios: 1:0 | 50% | 50% | - | - | - |
| **Diploid** | CC | CW | WW | | |
| Strand-ratios: 1:1, 2:0 | 25% | 50% | 25% | - | - |
| **Triploid** | CCC | CCW | WWC | WWW | |
| Strand-ratios: 2:1, 3:0 | 12.50% | 37.50% | 37.50% | 12.50% | - |
| **Tetraploid** | CCCC | CCCW | CCWW | CWWW | WWWW |
| Strand-ratios: 4:0, 3:1, 2:2 | 6.25% | 25% | 37.50% | 25% | 6.25% |

**Table S5**. **Presumed clonal CNA events in RPE cells detected by genomic sequencing**
Data shown for RPE-1, C7 and BM510. WGS: whole genome sequencing. MP: mate-pair sequencing. Only regions of 2Mb and longer are reported.
*This table is provided as an external data file.*

**Table S6: Summary of detected translocations**

Summary of translocations located using scTRIP. der: derivative. t: translocation. chr: chromosome. D: direct orientation. I: inverted orientation. H1: haplotype 1. H2: haplotype 2. BFB: breakage-fusion bridge. Mb: megabase.

| Sample | Derivative chromosome | Translocation Type | Orientation | correlation p-value (adj) | Derivative: haplotype | breakpoint | Partner: haplotype | breakpoint | Notes |
|---|---|---|---|---|---|---|---|---|---|
| C7 | der(10; t(10;15) | unbalanced | D | *p=3.45e-22* | chr10p-H2 | ends BFB | chr15q-H1 | 93.2Mb | provides telomere cap for BFB on chr10p |
| RPE-1 | der(X)t(X;10) | unbalanced | I | *p*=2.9e-33 | chrXq-H2 | Telomere | chr10-H2 | 60.8Mb | cytogenetically validated |
| BM510 | der(X)t(;10) | unbalanced | D | *p*=2.26e-32 | chrXq-H2 | Telomere | chr10-H2 | 60.8Mb | originated from RPE-1 cell line |
| BM510 | der(13)t(13;22) | unbalanced | D | *p*=5.52e-41 | chr13p-H2 | 19.4Mb | chr22q-H2 | 37.9Mb | rest of chromosome 13 is monosomic |
| BM510 | der(15)t(15;17) | reciprocal | I | *p=4.75e-29* | chr15q-H2 | 88.3Mb | chr17p-H2 | 19.6Mb | contains an inversion originating on chr17p; places TP53 upstream of NTRK3 |
| BM510 | der(17)t(15;17) | reciprocal | I | *p=3.93e-30* | chr17p-H2 | 19.6Mb | chr15q-H2 | 88.3Mb | |
| P33 | der(5)t(5;14) | reciprocal | D | *p=4.52e-5* | chr5q-H1 | 171.3Mb | chr14q-H1 | 98.7Mb | cyrptic (not seen in cytogenetic karyotype) |
| P33 | der(14)t(5;14) | reciprocal | D | *p=1e-4* | chr14q-H1 | 98.7Mb | chr5q-H1 | 171.3Mb | places BCL11B enhancer upstream of TLX3; cryptic |

*derivative assigned to chromosome containing centromere, translocation partners listed in numerical order
D = Direct (correlated segregation); I = indirect (anti-correlated segregation)

**Table S7**. **Inferred clonally present LOH events**

Summary of all LOH regions located in this study. CEN: spans centromere

| Sample | chrom | start | end | width | allele ratio | ploidy |
|---|---|---|---|---|---|---|
| RPE-1 | chr2 | 134959410 | 137376936 | 2,417,527 | 0.04268 | 2N |
| RPE-1 | chr5 | 98973704 | 101810758 | 2,837,055 | 0.03360 | 2N |
| RPE-1 | chr8 | 111323642 | 113738614 | 2,414,973 | 0.05549 | 2N |
| RPE-1 | chr11 | 66663975 | 68915891 | 2,251,917 | 0.12251 | 2N |
| C7 | chr2 | 134959410 | 137399019 | 2,439,610 | 0.05205 | 2N |
| C7 | chr5 | 98804837 | 101868951 | 3,064,115 | 0.04564 | 2N |
| C7 | chr8 | 111309080 | 113722108 | 2,413,029 | 0.08229 | 2N |
| C7 | chr11 | 66574908 | 68850811 | 2,275,904 | 0.10331 | 2N |
| C7 | chr13 | 1 | 114343387 | 114,343,387 | 0.04158 | N |
| BM510 | chr2 | 134883904 | 137364444 | 2,480,541 | 0.05813 | 2N |
| BM510 | chr5 | 98804837 | 101851871 | 3,047,035 | 0.04813 | 2N |
| BM510 | chr8 | 111347382 | 113731095 | 2,383,714 | 0.08725 | 2N |
| BM510 | chr11 | 66591291 | 68932307 | 2,341,017 | 0.11950 | 2N |
| BM510 | chr13 | 20839485 | 114342742 | 93,503,258 | 0.02428 | N |
| P33 | chr2 | 163723021 | 166686477 | 2,963,457 | 0.04431 | 2N |
| P33 | chr6 | 1 | 32348638 | 32,348,638 | 0.11968 | 2N |
| P33 | chr7 | 103695439 | 129082286 | 25,386,848 | 0.05444 | 2N |
| P33 | chr9 | 20511642 | 24727948 | 4,216,307 | 0.08911 | N |
| P33 | chr16 | 66813810 | 69469464 | 2,655,655 | 0.05423 | N |
| P33 | chr18 | 14791983 | 22033191 | 7,241,209 | 0.13534 | CEN |
| P1 | chr11 | 47840958 | 55257823 | 7,416,866 | 0.13867 | CEN |
| P1 | chr5 | 45508861 | 51382336 | 5,873,476 | 0.10649 | CEN |
| P1 | chr5 | 130961561 | 133293129 | 2,331,569 | 0.11553 | 2N |
| P1 | chr7 | 1 | 64381904 | 64,381,904 | 0.11537 | N |

CEN = centromere

### Supplementary Experimental Procedures

#### 1.1 Core computational framework for SV discovery in single cells

Our core computational framework (termed MosaiCatcher), described here in further detail, has been developed for detecting Dup, Del, Inv, InvDup, and 'other/complex SV' classes in single cells, based on scTRIP's SV diagnostic footprints. CNN-LOH events, altered cellular ploidy, and translocations are detected through separate modules defined around the core framework.

***Input data.*** Input data required by the framework are a set of single-cell (Strand-seq) BAM files from the same donor sample. In our study, these data were aligned to build GRCh38 of the human reference genome (GCA_000001405.15_GRCh38_genomic.fna). To later enable haplotype phasing and haplotype-resolved SV assignments, our framework performs re-genotyping of SNPs provided by the 1000 Genomes Project (1000GP; phase 3) to detect heterozygous sites from the single-cell input data. When using our framework a VCF file with these 1000GP SNP sites is to be provided as input. Alternatively, the MosaiCatcher pipeline is able to call SNPs directly from the single-cell data or to use externally generated SNP calls for a given sample, *e.g.* based on bulk WGS. Additionally, a tab-separated file with normalization factors (see below) per bin across the genome is used as input to the framework.

***Workflow management.*** Our core framework is implemented as a Snakemake workflow[31], to facilitate reproducibility and scalability. Source code is available at github (https://github.com/friendsofstrandseq/pipeline) and an overview of different steps is shown as **Figure S3**. The software requirements are described as a Bioconda environment[29], again to facilitate reproducibility by allowing for easy installation of all dependencies by executing a single command. To ensure computational efficiency, a number of functionalities inside the core workflow have been implemented in C++ and are hosted in a separate repository (https://github.com/friendsofstrandseq/mosaicatcher); we refer to the corresponding executable as "`mosaic`" in the following. Most additional functionalities, such as the Bayesian model for SV classification are implemented in R and are distributed as part of the core workflow.

***Binned read counting in single cells.*** At first, reads in all individual cells are binned, for each strand (`mosaic count` command). Bins have a fixed width (default: 100kb), starting from position 0 up the end of the chromosome. Mapped reads were assigned to bins based on their start position and filtered according to the following criteria: non-primary and supplementary alignments are excluded; alignments with the QC failure flag are excluded; PCR duplicates are excluded; reads with mapping quality <= 10 are excluded. In case of paired-end data only the first read of each pair (based on the BAM flag 0x40) was used to avoid double-counting. Cells with too little coverage (median count per bin of 3 or less) were removed by default. The parameters $p$ and $r$ of the NB distribution were determined in the same manner as for SV classification (see respective section below). During parameter estimation, bins were excluded from the parameter estimation process if their mean coverage across all cells was very low (<0.1, where coverage was previously normalized to 1) or if they showed a highly abnormal WC/(WC+CC+WW) fraction ($WC_{frac}$) across cells. Bins were deemed abnormal if exhibiting either $WC_{frac}$<0.05 or $WC_{frac}$> 0.95, reflecting bins that either never showed WC status, or those that exhibited always WC status, as for example often seen in regions within or near centromeres[32].

***Coverage normalization in single cells.*** Our framework pursues normalization of read coverage prior to SV calling (**Fig. S5**). To estimate suitable parameters for normalization, we analyzed Strand-seq data recently generated by the Human Genome Structural Variation Consortium (HGSVC) comprising 9 lymphoblastoid cell lines from the 1000 Genomes Project (1000GP) (*i.e.*, samples NA19238, NA19239, NA19240, HG00731, HG00732, HG00733, HG00512, HG00513, and HG00514)[33]. We utilised 1058 cells from these HGSVC samples sequenced via Strand-seq, obtained from ftp.1000genomes.ebi.ac.uk/vol1/ftp/data_collections/hgsv_sv_discovery/ working/20151203_strand_seq/, and subjected these cells to the same binning scheme described above. Analysis of several of these 1000GP samples showed that these do not carry any germline copy number variants (CNVs) ≥200kb[34]. To identify a scaling factor for normalization we aggregated these HGSVC Strand-seq data, and first masked regions using any of the following 'exclusion criteria': observed mean coverage <50%, observed mean coverage >200% (**Fig. S5**), or observed standard deviation larger than the mean coverage. Then, using the remaining bins, we modeled the observed mean bin coverage in our test samples assuming a linear relationship to the mean HGSVC bin coverage, which explained 66% of the variance with a slope of ~0.6. We used this linear relationship to derive a scaling factor for each bin, which subsequently was applied to all cells of our study.

***Blacklist construction.*** We created a "blacklist" of regions exhibiting strong sequencing/mapping abnormalities to avoid false positive somatic variant calling. To construct our blacklist, we started from the 'masked regions' with unusual coverage in the independent HGSVC samples (see previous paragraph). We then progressively merged such intervals if they exhibited a distance of 500kb or less (which avoided generation of a highly fragmented blacklist). Lastly, we ensured that no known polymorphic inversion[32] was accidentally masked by removing all intervals from our blacklist that overlapped with a germline inversion larger than 100kb in size reported by the HGSVC[33]. The resulting blacklist was used in all following analyses, which considered regions outside of the blacklisted intervals for single cell SV calling.

***Joint segmentation of single cells.*** We followed the strategy suggested by Huber *et al.* to perform segmentation on a multivariate input using a squared-error assumption[35]. Therefore, the binned read count data for all single cells of a sample were simultaneously used as input, with the rationale that SVs that recur in multiple cells can reinforce each other. Given a number of allowed change points *k*, a dynamic programming algorithm finds the discrete positions of the *k* change points with minimal sum of squared error (SSE). The change points at level *k* are computed using knowledge about a set of *k-1* optimal change points through dynamic programming[35]. This algorithm uses a cost matrix, to determine the cost (summed squared error) of every possible consecutive segment. While the same direction of change was assumed in all samples in the original implementation of Huber *et al.*, we adapted the algorithm to calculate this cost matrix for each cell and strand separately. We additionally adapted the cost matrix to penalize segments which are below 200kb in size, as a means of avoiding over-segmentation. The segmentation procedure (`mosaic segment`), performs the segmentation separately for each chromosome and outputs the resulting change points up to a maximum number of allowed change points. We selected appropriate segmentation parameters by assessing the benefit of increasing the number of change points (k) in terms of the summed squared errors (SSE) of the piecewise constant function compared to the actual count data. Let $SSE_k$ be the residual error associated to partitioning a chromosome into k segments. We then select the smallest number *k* such that $SSE_k$ - $SSE_{k+1}$ is below a user-set parameter (default: 0.1, which is used in this study) to adjust the number of change points *k* for a chromosome.

***Strand state and SCE detection in single cells.*** Detecting SV diagnostic signatures depends on whether the corresponding segment in a single-cell followed a WW, CC, WC, or CW pattern of mitotic segregation (**Table S1**). We refer to the underlying baseline distribution of W and C reads along a chromosome as the "ground state" (see **Methods**). While the ground state usually stays the same along the length of a chromosome, it can be altered by sister chromatid exchanges (SCEs), which underlie mitotic patterns of recombination unrelated to structural variation[2]. Changepoints in Strand-seq data that result from mitotic recombination events/SCEs represent a source of "noise" that MosaiCatcher is able to correct for. Fortunately, SCEs happen independently in each single cell[2], and unlike SVs, SCEs are not transmitted clonally to daughter cells (*i.e.* are only detectable in the cell they occur in[2]). Hence, changepoints resulting from SCEs are very unlikely to recur at the same position in >1 cell of a sample[1,2]. MosaiCatcher uses changepoint recurrence as a key criterion for distinguishing SCEs from SVs. To identify SCEs, we employed the same segmentation strategy as described above, but to each single cell *separately* rather than jointly. To do so, the threshold to select the number of breakpoints k (see above) was set to 0.5. We assigned an *observed state* to each resulting segment by computing the fraction $f_{WC} = W/(W + C)$ and assigning state WW if $f_{WC} > 0.8$, state CC if $f_{WC} < 0.2$ and state WC/CW otherwise. The states of neighbouring segments were compared to each other and if the states were unchanged the intervening changepoint was discarded, while the remaining changepoints were subsequently further considered as putative SCEs. Note that we write "WC/CW" to indicated that we are not making a distinction between these two states in this step, distinguishing the two happens in the subsequent StrandPhaseR step (see **Fig. S3**).

An important consideration is that in some cases, changepoints detected in this way may correspond to SVs rather than SCEs. We thus employed the following strategy to select a high confidence list of SCEs: We first select those changepoints far away (>500kb) from any breakpoint identified during the joint segmentation (see previous paragraph); these changepoints are likely to represent true SCEs. With this provisional set of candidate SCEs, we considered each of the three ground states WW, CC, WC/CW to determine a plausible "ground state". We employed the assumption that a given state at the beginning of a chromosome and a set of SCE positions (which change the state) uniquely determine the state for every segment on the chromosome. To assess which of the three ground states (WW, CC, or WC/CW) at a chromosome start to pick, we computed the *discordant length*, defined as the total length of genomic intervals for which the observed state differs from the predicted ground state. Although highly unlikely, in rare occasions, an SCE changepoint may appear to coincide with an SV breakpoint. In order to enable MosaiCatcher to recover such rare SCEs, all putative SCEs closer than 500kb to a breakpoint in the joint segmentation were analyzed. If adding one of these putative SCEs reduces the *discordant length* by 20Mb or more, MosaiCatcher assigns these SCE status. Doing so, MosaiCatcher is able to avoid that missed SCEs result in an incorrectly assigned ground state along larger parts of a chromosome. Note that adding at most one such additional SCE precludes masking most true SVs, which have two breakpoints, whereas SCEs lead typically only to a single "switch" (changepoint) in W and C states along a chromosome [2]. Also, it should be noted that since SCEs never associate with copy-number alteration, the chance that SCEs are confused with SVs is near "zero" for many SV classes - that is for, Del, Dup, InvDup, and complex rearrangements - even if these SV are present only in a single cell. Thus, in reality, SCEs are only very rarely incorrectly assigned SV status (as also evidenced by our experimental validation data).

***Chromosome-length haplotype phasing using single-cell sequencing data***. To facilitate haplotype-aware SV calling, we phased all available chromosomes using StrandPhaseR[36]. While building whole-chromosome haplotypes for a sample, we assigned regions represented by both W and C strands as either WC or CW for each cell. That is, we used reads overlapping heterozygous SNPs to determine whether haplotype H1 was represented by W reads and H2 by C reads (a situation we denote as WC), or vice versa (denoted CW) (see **Methods)**. In addition to this refined characterization of the ground state, StrandPhaseR outputs the chromosome-wide haplotypes as a VCF file, which we later utilized in the "haplotagging" step. This phasing step of our framework requires at least a few dozen SNPs per chromosome. To ensure availability of enough SNPs, we re-genotyped germline variants previously identified in the 1000GP[27], using Freebayes[26] with options "`-@ <1000GP-snps.vcf> --only-use-input-alleles <input.bam> --genotype-qualities`". We retained all heterozygous SNPs with QUAL>=10. Alternatively, our framework can use externally provided SNPs. To boost the usable coverage for SNP calling, we performed a cell sorting experiment, independently sorting 100 cells (termed the '100 cell control') in each sample, followed by short-read whole genome sequencing to 1.9x mean coverage.

***Estimating Negative Binomial parameters.*** The number of high throughput sequencing reads mapped to genomic windows (or bins) were previously shown to be in agreement with a negative binomial (NB) distribution[37], which can account for overdispersion. We employed the NB distribution as the basis for our Bayesian framework. The NB distribution has two parameters, $p$ and $r$, which are estimated from the observed read counts as follows. Let us denote the value $n$ as the number of single cells analyzed in a sample. We assume that the number of reads sampled from each single-cell at a fixed bin size is an NB random variable. In reality, the coverage of single cells will be varying resulting in different NB parameters for each cell. Key for parameter estimation is that not only the coverage of individual single cells, but also the total coverage of all single cells together, are derived from an NB distribution. This implies that all single cells should have the same $p$, therefore there are $n$+1 free parameters to estimate (one $p$ parameter and $n$ dispersion parameters).

In an NB distribution, the ratio of the mean to the variance is equal to 1-$p$. Having the same $p$ parameter over all single cells implies that the ratio of mean to variance is constant across all single cells. Consequently, the mean and variance of binned read counts among single cells share a linear relationship in which the line connecting these mean-variance points for single cells passes the origin coordinate with a slope determining the $p$ parameter. This relationship allows estimation of the shared $p$ parameter: for each single-cell, we compute the empirical mean and variance of the observed read counts in fixed-sized bins across the genome. If we denote the set of empirical mean-variance pairs by $(m_1, s^2_1), (m_2, s^2_2), \ldots,$ and $(m_n, s^2_n),$ the $p$ parameter is estimated as follows:

$$p = \frac{\sum_{i=1}^{n} m_i}{\sum_{i=1}^{n} s^2{}_i}$$

After obtaining $p$, we estimate the dispersion parameter $r_j$ of each single cell $j$ by setting the distribution mean to the average read count per bin of that single cell. We employed a trimmed mean for estimating the dispersion parameters (with trim parameter set to 0.05), to remove the effect of abnormally high or zero read counts (e.g. seen in regions of low mappability).

***SV diagnostic footprints***. Each SV diagnostic footprint (**Fig. 1**) can be translated into the expected number of copies sequenced in W and C orientation contributing to the genomic segment under consideration. **Table S1** shows this relationship for each SV class, both for chromosomes where both haplotypes are represented by different template strands (here referred to as 'WC/CW chromosomes') and for such where both haplotypes are represented by the same template strand ('WW chromosomes' and 'CC chromosomes'). Every haplotype-resolved SV implies a particular segment strand pattern in WC, CW, WW, and CC chromosomes, respectively. For example, if the ground state of a single cell in a chromosomal region is WW and the SV status in a segment in that region is 'inverted duplication of the paternal haplotype represented on the W strand', the observed segment strand pattern will be WWC in this given single cell. By comparison, if the ground state is WC (W for the H1 haplotype), and the SV status is deletion of the H1 haplotype, the observed segment strand pattern is C (see **Table S1**). These expectations are formalized in our Bayesian model, which we describe in the following.

***Bayesian model to compute haplotype-aware SV genotype likelihoods in single cells.*** We utilized a Bayesian model (**Fig. S4**) to compute haplotype-resolved SV genotype likelihoods for each segment in each single cell. We model $V$, the SV type to be inferred, as a pair $(\overrightarrow{C}, \overleftarrow{C})$, where $\overrightarrow{C}$ gives the number of copies of that segment in forward direction and $\overleftarrow{C}$ gives the number of copies of that segment in reverse direction (i.e. when an inversion is present). That is, the pair (1,0) encodes the reference state of a haplotype (one forward copy and zero inverted copies). As illustrated in **Fig. S4**, each segment $k \in K$ and haplotype $h \in H = \{h_1, h_2\}$ in single cell $j \in J$ comes with a variable $V$ for this SV state, which we refer to as $V_{j,k,h}$. Together with the ground state $T$, each SV state $V$ deterministically leads to a corresponding "copy number" observed in Crick direction $N^C$ and in Watson direction $N^W$, as explained in the previous section on SV diagnostic signatures (also see **Table S1**). Conditional on the sum of Crick and Watson copy numbers of both haplotypes, the corresponding coverages $X^C$ and $X^W$ are assumed to follow a negative binomial (NB) distribution

$$X^W_{j,k} \mid (N^W_{j,k,h_1} + N^W_{j,k,h_2}) \sim NB(r^W_{j,k}, p)$$
$$X^C_{j,k} \mid (N^C_{j,k,h_1} + N^C_{j,k,h_2}) \sim NB(r^C_{j,k}, p)$$

for each single cell $j$ and segment $k$. Here, $p$ is the estimated common $p$-parameter of the NB distribution (see *Estimating Negative Binomial parameters* above), and $r^W_{\mathrm{J},}$ and $r^C_{\mathrm{J},}$ are proportional to the estimated parameter $r_j$ (also see above), the segment size $L_k$ and the Watson and Crick segment copy numbers ($N^W_{j,k} = N^W_{j,k_{h1}} + N^W_{j,k_{h2}}$ and $N^C_{j,k} = N^C_{j,k_{h1}} + N^C_{j,k_{h2}}$) and hence are computed as follows (for $d \in \{W, C\}$):

$$r^d_{j,k} = \begin{cases} \frac{1}{2}\alpha r_j L_k & \text{if } N^d_{j,k} = 0 \\ \frac{1}{2}(1-\alpha) r_j L_k N^d_{j,k} & \text{otherwise} \end{cases}$$

In this formula, $\alpha$ is a parameter in our model indicating the fraction of "background reads", which represents noise in Strand-seq data (for example due to regions with incomplete BrdU incorporation or removal)[1,2]) These background reads are taken into account by assuming $\alpha = 0.1$, which reflects an upper bound for the abundance of such background reads observed in practice. Note that the $\frac{1}{2}$ coefficients in the above formula serve to scale the dispersion parameter to copy number 1 ($r_j$ is estimated above to reflect a diploid state of copy number 2). In summary, every haplotype-resolved SV class ($V$) in a segment together with the ground state ($T$), define a Watson and Crick copy number

($N$) used to compute the NB likelihood of observed read counts. Through this mechanism, we obtain likelihoods for all diagnostic signatures in **Table S1**.

***Incorporating haplotype-specific sequencing reads ('haplotagging')***. One of the key advantages of scTRIP is the ability to utilize haplotype information made available through strand-specific sequencing. In the base model described in the previous paragraph, this haplotype-awareness is brought forth by distinguishing WC from CW ground states (also see *Chromosome-length haplotype phasing using single-cell sequencing data)*. Our framework is additionally able to make use of reads not directly assigned to a haplotype (i.e. those in WW and CC regions) owing to their overlap with a haplotype-phased SNP. This feature can further facilitate validation and falsification of putative SVs seen only in few or even individual cells. We utilize the whole-chromosome haplotypes generated using StrandPhaseR [36] to tag reads by haplotype using the 'haplotag' command of WhatsHap [38,39], resulting in one 'haplotagged' BAM file per single cell library. These BAM files are then used to compute the number of Watson/Crick reads that could be tagged by haplotype H1/H2, respectively, for each segment and each single cell. The resulting happlotagged read counts are incorporated in the Bayesian model as random variables $X^W_{tag}$ and $X^C_{tag}$ (see **Fig. S4**). We employed a multinomial distribution to model the conditional distribution of these tagged read counts given the (haplotype- and strand-specific) copy numbers $N^C$ and $N^W$. More precisely, we defined parameters of the multinomial distributions $p^C_{j,k,h_1}$, $p^C_{j,k,h_2}$, $p^W_{j,k,h_1}$, and $p^W_{j,k,h_2}$, for each segment $k$ and single cell $j$, such that they are proportional to the corresponding copy numbers:

$$p^d_{j,k,h} \propto \max(\alpha, N^d_{j,k,h})$$

where $d \in \{W, C\}$ as before. Here, $\alpha$ is again a rate of background reads (set to $\alpha = 0.1$) and the $p^d_{j,k,h}$ are normalized to sum up to one. Given the total number of reads and the (haplotype- and strand-specific) copy numbers $N^C$ and $N^W$, the tagged reads are multinomially distributed:

$$(X^C_{j,k,h_1,tag}, X^C_{j,k,h_2,tag}, X^W_{j,k,h_1,tag}, X^W_{j,k,h_2,tag}) \sim \text{Multinomial}(p^C_{j,k,h_1}, p^C_{j,k,h_2}, p^W_{j,k,h_1}, p^W_{j,k,h_2})$$

***Employing the Bayesian model for SV calling.*** To utilize our Bayesian model for SV calling, we defined prior probabilities and combined them with the model-based likelihoods for each single cell and segment. We started by regularizing the raw likelihoods, adding a small constant (set to $10^{-6}$) to all likelihoods and renormalizing afterwards. This ensures that very small values (or hard zeros) are avoided and corresponds to the error assumption that every SV genotype is possible with this given small probability, no matter what the data suggests. Then, we used two forms of priors. First, we captured biological knowledge on the plausibility of observing certain event types. To do this, we defined the priors to be proportional to a pre-specified constant per SV type and chose these constants as follows: ref=200, del/inv/dup=100, invdup=90, other/complex=1. While this choice is somewhat arbitrary, it encourages the SV calling process to prefer the reference state (ref) over canonical SVs (del/inv/dup/invdup) over more exotic SV classes, for example involving an inversion on one haplotype and a deletion on the other haplotype (other/complex) - unless the model observed sufficient evidence to overwhelm these priors. Thus, we required the caller to gather more evidence for SV classes deemed implausible. The second type of priors we applied acts on each segment separately and uses the raw likelihoods computed by the model across all cells to compute a probability distribution over all SV types. That is, for each segment we summed up the likelihoods per SV type across all cells and normalized to one, which corresponds to estimating the frequency of each

SV genotype for that segment. The intuition behind this procedure is that we need to encourage the SV caller to prefer SV types present in many cells over those SV types present only on few cells - unless the evidence inherent to the genotype likelihoods is strong enough to overwhelm these priors. Before applying these priors, we set the prior of each SV genotype to zero if the estimated frequency of that genotype was below a threshold, which we term GTCUTOFF (set to 0.05 for the strict call set and set to 0 for the lenient call set). Effectively, this means that the strict parameterization only considers an SV genotype if the likelihoods across all cells suggest it to be present in the cell population at an expected frequency of at least 5%. The lenient call set, in contrast, disables this cutoff by setting it to zero and hence readily permits SV genotypes present in individual cells only (for more details see *Strict and lenient parameterizations of our single cell SV discovery framework*). Lastly, we used the resulting posterior probabilities to compute log odds ratios (of an SV genotype vs. the reference state), and accepted an SV call if the log odds ratio was at least 4. SV calls in segments with >20% blacklisted bins were discarded (see *Blacklist construction*).

***Call set post-processing:*** *Filtering:* We developed a filtering routine to be used only in conjunction with the strict parameterization, the main goal of which is to arrive at a high confidence SV callset for all SVs with VAF greater than 5%. This filtering routine removes rare inversions seen in only 1 or 2 cells, since rare inversions may occasionally correspond to SCEs. This routine further removes SV calls exhibiting particular biases, most importantly, those biased to occur largely in the context of a certain ground state. In particular, while SVs can be detected in the context of all four ground states (WW, CC, WC and CW; see **Table S1**), we noticed during the development of MosaiCatcher that artifactual SV calls can occasionally arise on WW or CC chromosomes, where the ability of the caller to measure gains or losses in read depth is reduced. Calling deletions or duplications on WW or CC chromosomes is indeed conceptually related to previously developed copy-number profiling methodology; *i.e.*, SVs called on WW or CC chromosomes will not benefit from the ability of scTRIP to call these SVs based on strand-specific read depth gain or loss (**Fig. 1**, **Table S1**).

*The following hard filters were implemented to be used with the strict parameterization:*

(i) Removal of inversions seen in less than 3 cells.

(ii) Removal of deletions seen in multiple cells, if these show a bias towards occurring mostly in WW and CC chromosomes with less than a third seen in WC or CW regions (deletions with log odds ratio ≥50 will not be removed by this hard filter). As reasoned further above, we implemented this filter since deletions that are repeatedly seen in the WW or CC ground state, but not or only rarely in the WC ground state, are (according to our experience) of lower confidence.

(iii) Removal of duplications seen in multiple cells, if these show a bias towards occurring in WW and CC chromosomes, with less than a third seen in WC or CW chromosomes (duplications with log odds ratio ≥50 will not be removed by this hard filter). As reasoned further above, we implemented this filter since according to our experience duplications that are repeatedly seen in the WW or CC ground state, but not or only rarely in the WC ground state, are of lower confidence.

(iv) Removal of SVs overlapping UCSC annotated segmental duplications in the genome (file: segDups_hg38_UCSCtrack.bed.gz) by more than 50% (we found such SV calls to be of lower confidence).

*Merging:* We also developed a merging routine to be used in conjunction with the strict parameterization, which groups adjacent SVs with a similar VAF (where VAF ≥ 0.1) into a single SV call to avoid over-segmentation and produce a final high confidence somatic SV sites list. To this end, we considered VAFs of adjacent SVs to be similar if $VAF_{SV1}/VAF_{SV2} \geq 0.75$ (for cases where $VAF_{SV2}>VAF_{SV1}$) or $VAF_{SV2}/VAF_{SV1} \geq 0.75$ (for cases where $VAF_{SV1}>VAF_{SV2}$), and grouped all immediately neighboring SVs selected by this similarity criterion. In our experience, SVs merged by this routine will nearly always correspond to a single structural variation event in validation experiments.

***Strict and lenient parameterizations of our single cell SV discovery framework.*** As alluded to above, our framework comes with the ability to adjust for the tradeoff between sensitively calling SVs present at low VAF, and accurately identifying SVs consistently seen among cells. We parameterized this tradeoff into a 'strict' and 'lenient' SV caller, whereby the 'strict' caller optimizes precision for SVs seen with VAF≥5%, while the 'lenient' caller targets all SVs including such present only in a single cell. These parameterizations differ in three settings: the GTCUTOFF (see *Using the Bayesian model for SV calling*), whether or not haplotagged reads counts are incorporated (see *Incorporating haplotype-specific sequencing reads)*, and whether filtering is enabled (see *Call set post-processing*). The strict caller uses GTCUTOFF=0.05, while the lenient caller uses GTCUTOFF=0. For the strict caller, we disabled the haplotagging feature, while we enabled haplotagging for the lenient call set - with the reasoning that haplotagging is mostly valuable to resolve putative SVs with low VAF. Lastly, we used the filtering described in the previous paragraph for the strict caller, while we proceed with the unfiltered set for the lenient caller. We recommend use of the strict caller to enable reliable detection of subclonal SVs down to a VAF of 5%. The lenient caller should be used for analyzing SVs across the whole VAF spectrum down to the individual cell.

#### 1.2 Simulations to evaluate our framework for single cell SV discovery

We devised two procedures to perform simulations enabling evaluation of our framework for single cell SV discovery. The first procedure - referred to in the main text and in **Fig. S7** and not further detailed in this Supplementary Material section - employed *in silico* mixtures of cells from clonal RPE cell lines. The second procedure, outlined below, inserted randomly picked SVs of arbitrary type and size into subclonal cell fractions. While naturally simulations represent idealized conditions, they closely reflect several essential properties of Strand-seq data, and thus can facilitate methods development.

In particular, we devised a subcommand of MosaiCatcher to simulate Strand-seq experiments on a cell population by sampling binned read counts from a negative binomial (NB) distribution. At first, the basic read coverage of two homologues is generated via sampling from a NB distribution. Each haplotype was covered either with W or with C reads during the simulation, as seen in regular Strand-seq libraries. To do this, we set the bin size to 100kb, the NB parameter $p$ to p=0.28301 (corresponding to the lowest value observed for the datasets used in this study), and the expected number of reads per library to 300,000-500,000 reads (sampled uniformly to reach 300,000-500,000 reads) per cell. SVs were implanted into individual chromosomal homologs based on the expected consequence these events have on a haplotype-resolved genome. In the case of a heterozygous deletion, for example, the coverage of one homologue was set to 0 with a randomly assigned fraction of background reads per bin, by default 0.05, to simulate noise. Inversions were introduced by flipping orientation of counts in the affected region. In each simulation, MosaiCatcher implants 25 SVs of a given SV class and with given size and VAF, by randomly placing them along the genome (with a minimum distance of 1Mb between variants). We sampled SV sizes uniformly on a log scale (*i.e.*, preferring smaller variants over larger ones), and placed SVs randomly in a bin-unaware manner (i.e. without requiring that the start and end align with the boundaries of bins). Variants with a clonal fraction *f* <1 were incorporated into subsets of cells, chosen with a probability *f* for each cell. Finally, after implanting SVs into the genome, we additionally simulated on average 3.78 SCEs into each cell (reflecting the average number of SCEs typically seen in RPE-1 data using Strand-seq). VAF values shown in **Fig. S5** were computed based on the number of cells in which the SV was placed during the simulation, divided by the total number of cells simulated. We evaluated the precision and recall of our framework for detecting different classes and sizes of subclonal SVs, as shown in **Fig. S5**.

### 1.3 Comparing the scTRIP framework with another CNA detection tool

We additionally compared our framework's Del and Dup calls with Aneufinder [40] (version 1.8.0; https://bioconductor.org/packages/release/bioc/html/AneuFinder.html), a CNA detection tool suitable for operating on sparse single cell data. The exact settings used to run Aneufinder are listed below:

```
Aneufinder(inputfolder = <bam.dir>, outputfolder = <output.dir>, reuse.existing.files = TRUE, binsizes = 100000, pairedEndReads = TRUE, chromosomes = paste0('chr', c(1:22, 'X')), remove.duplicate.reads = TRUE, min.mapq = 10, use.bamsignals = TRUE, method = 'edevisive', strandseq = TRUE, blacklist = 'mosaicather_specific_blacklisted_regions')
```

We analysed our RPE single cell data. To set a ground truth CNA set for our comparison, we made use of CNA regions inferred using Delly2[9], by selecting events with ≥200kb in size that showed a paired-end signal and displayed a read-depth shift compared to flanking regions (this ground truth dataset is available as **Table S5**). For deletions and duplications we required a minimum read-depth shift of 0.8 and 1.2, respectively. Any CNA region detected either by scTRIP or Aneufinder that overlapped with our truth set was considered as true positive call, or was considered as a false positive if missed. Sporadic (single-cell specific) Aneufinder calls that did not reach ≥30% VAF were not counted as false positives. VAF was calculated as a fraction of gains and losses detected across cells. We performed two independent analyses, (1) evaluating CNA calls ≥200kb and (2) evaluating for CNA calls ≥400kb. In both settings, scTRIP performed better than Aneufinder (*e.g.* yielding 26.1% more true positive calls for CNAs ≥200kb, and 13.8% more true positive calls then Aneufinder, while at the same time producing less false positives than Aneufinder; see **Fig. S8**).

### 2.1 Single cell dissection of complex inter-chromosomal rearrangements

To infer translocation partners by scTRIP, we subjected candidate translocation segments to template strand co-segregation analysis. Candidate segments were identified based on recurrent breakpoints that flagged regions failing to co-segregate with the chromosome they originated from. Co-segregation analysis involved investigating pairwise correlations in template strand identity between candidate translocation segments and potential partner segments across the genome. These principles, which allowed us to reconstruct complex derivative chromosomes by single cell strand sequencing, are detailed with several examples below.

***Single cell sequencing based dissection of an "unsequenceable" translocation in BM510.***

We first aimed to assess and verify the herein introduced diagnostic footprints for translocations in RPE-1, in which a well-documented balanced translocation involving chromosomes X and 10 (der(X) t(X;10)) was previously characterized microscopically using spectral karyotyping[10] - albeit not yet resolved with DNA sequencing. We initially analyzed BM510 Strand-seq single cell data to identify segments with an 'irregular' strand-pattern representing 'candidate translocation segments' (see **Fig. 3A**). In an unbalanced translocation, according to our diagnostic footprints (**Fig. S1**), only one of the translocated segments (*i.e.* the segment exhibiting a read depth increase) will show an 'irregular' strand-pattern (that is, represent a candidate translocation segment). This segment, while being 'inconsistent' in terms of strand-state with the remaining regions of the chromosome the segment originated from, will exhibit a strand-pattern correlating with the strand-pattern of its translocation partner.

Analysis of BM510 using our framework identified a large segment on chromosome 10 as 'irregular', and thus representing a 'candidate translocation segment' (**Fig. 3A**). We performed single cell-based haplotype analysis to infer the haplotype of this irregular chromosome 10 segment. We only considered those cells in which a single haplotype could be unambiguously separated as a single copy either represented by a Watson (W) or Crick (C) template strand (see **Fig. S9**). We subsequently performed chromosome-scale haplotyping of these selected single strands (**Methods**) to derive consensus haplotypes for chromosome 10. The identity of the translocated chromosome 10 haplotype was established by comparing single cell haplotypes inferred for the 'candidate translocation segment' to both consensus haplotypes derived from all haplotype informative single cells generated for BM510. We detected a single chromosome X haplotype (H2) that correlated in its strand states with this chromosome 10 segment - indicating that the segment co-segregated with chromosome X H2 haplotype during mitosis (**Fig. 1B**, **Fig. 3B)**. The false discovery rate (FDR)-adjusted p-value, in an analysis comprising 118 cells from BM510 that showed consistent full-length (or majority) strand states on chromosome X (*i.e.* not affected by SCEs), was $p$=2.9e-33 (Fisher's exact test). None of the other FDR-adjusted p-values, obtained by assessing co-inheritance of the 'irregular' chromosome 10 segment with other genomic regions, reached significance ($p$>0.01). Thus, our framework was able to readily and unambiguously identify the previously microscopically detected der(X) t(X;10) in the RPE-1 clone BM510, and to fully haplotype resolve this event, despite being incomplete accessible to WGS (**Fig. S10**). We repeated this analysis for RPE-1 and detected the same event with significant adjusted p-value (p<0.01), consistent with this translocation being present in the RPE-1 parental line[10].

***Karyotyping a derivative chromosome with scTRIP: placement of the unbalanced der(X) t(X;10) translocation to chromosome X q-arm.***

It can be reasoned that the translocated chromosome 10 segment would, likely, be fused either to the Xp-arm or the Xq-arm in the context of this inferred unbalanced translocation. To precisely order and orient the translocated chromosome 10 segment with respect to chromosome X H2, we made use of sister chromatid exchange events (SCEs) detectable by Strand-seq[2] (see "*Strand state and SCE detection*" below). We selected all regions in BM510 with a change in strand state corresponding to an SCE (CC/WW to WC, or WC to CC/WW), keeping only those that mapped to either end of the chromosome – a condition true in 24 cases. In these cases we gathered the strand state at the end of chromosome X haplotype 2 (p-arm and q-arm) and the translocated chromosome 10 segment in two separate contingency tables. Next, we calculated the FDR-corrected p-value for both tables in the same way as described above, obtaining a p-value of 1.7e-05 for the q-arm (Fisher's exact test). The p-arm received an FDR-adjusted p-value of 0.69. Cytogenetics (*i.e.* spectral karyotyping[10]) shows that the translocation is indeed attached to the end of the q-arm of chromosome X, consistent with our scTRIP based inference.

***Allele-specific analysis places the tr(X;10) translocation to the active X chromosome.***

The two X chromosomes in females function differently, with one being transcriptionally active and the other silenced via X inactivation[11]. Taking advantage of the haplotype-resolution of scTRIP, we generated bulk RNA-seq data and characterized RPE-1, C7 and BM510 for patterns of X inactivation using allele-specific gene expression analyses. In order to analyze allele-specific expression, we aligned raw RNA-seq data from BM510 and RPE-1 to the human reference genome (GRCh38) using STAR aligner (2.5.3 version). Allelic read counts at heterozygous SNP sites were obtained using the ASEReadCounter, a method provided by the GATK package[12], using the following parameters:

```
GenomeAnalysisTK.jar –R <reference.fasta> -T ASEReadCounter -o
<output.csv> -I <input.bam> -sites
<chrX_phased_from_MosaiCatcher.vcf> -U ALLOW_N_CIGAR_READS --
minMappingQuality 10 --minBaseQuality 2 -drf DuplicateRead
```

We assigned allelic read counts to either H1 (haplotype 1) or H2 (haplotype 2) based on our framework's whole-chromosome phasing information. Every SNP site was annotated with gene locus information using the Homer annotate Peak tool[13]. Intergenic RNA-seq reads were excluded, while intronic reads were kept for haplotype-specific expression analyses (these can reveal the level of nascent transcript before splicing[14,15]). Read counts for heterozygous SNPs within the same gene locus were aggregated into gene-level read counts. Differential expression of the H1 and H2 allele of each gene was evaluated using the likelihood ratio test, followed by FDR adjustment using the Benjamini-Hochberg procedure[16], as provided by EdgeR[17]. Allelic counts revealed fusion of the duplicated chromosome 10 haplotype to the active, rather than the inactive X chromosome, in RPE-1 and BM510 (**Fig. S11**). Genes residing on the duplicated 10q haplotype, furthermore, showed specific increases in allele-specific expression when compared the non-duplicated 10q haplotype (**Fig. S11**), corroborating our haplotype assignments.

***Single cell discovery of an unbalanced translocation in BM510 involving chromosomes 13 and 22.***

We also detected another unbalanced translocation, connecting chromosomes 13 and 22 in BM510. A duplicated chromosome 22 segment correlated significantly in strand states with a proximal segment of chromosome 13 ($p$=5.52e-41, Fisher's exact test, FDR-adjusted), consistent with a der(13) t(13;22) translocation. WGS data generated for BM510 verified this scTRIP-discovered translocation (**Table S5**).

***Single cell dissection of a complex translocation t(15;17) mediating a TP53-NTRK3 gene fusion.***

We also more closely analyzed a reciprocal translocation involving chromosomes 15 (haplotype 2, H2) and 17 (haplotype 2, H2) that our scTRIP based framework revealed in BM510. The involved segments showed perfectly inverse correlations in terms of strand states with the recipient chromosomes (FDR-adjusted $p$=4.75e-29 [chr15/chr17tr] and $p$=3.93e-30 [chr15tr/chr17], respectively; Fisher's exact test). This can be explained by the fusion of both chromosomal segments occurring with one chromosome being inverted with respect to the translocation partner (**Fig. 3BC**) - a relative 'reorientation' of chromosomal segments that makes intuitively sense, allowing the derivative chromosomes to retain telomeric sequence at the chromosome's tips (**Fig. 3C)**. scTRIP based analysis, notably, revealed this inter-chromosomal rearrangement to be complex, involving an additional ~12Mb balanced inversion on chr17p (placed to 7.7-19.6Mb) (**Fig. 3BC)**. By pooling the BM510 single cell sequencing data, we carefully analyzed the breakpoints of the copy-balanced intra-

and inter-chromosomal rearrangements of 15q and 17p. This revealed that the proximal breakpoint of the chr17p inversion disrupted *TP53* and physically placed the 5' coding region of *TP53* to the distal translocation breakpoint (*i.e.* re-locating it to 19.6Mb of 17p). As a consequence of the additional translocation arising between chr15q and chr17p, the 5' end of *TP53* was thus juxtaposed to the 3' end of the oncogene *NTRK3* (located at the 88.3Mb chr15q breakpoint). This complex rearrangement resulted in a candidate gene fusion event between *TP53* and *NTRK3*. Fusion products leading to the overexpression or activation of genes from the *NTRK* gene family have been previously observed in different cancers[18]. We confirmed the over-expression of *NTRK3* in BM510 by performing RNA-seq (see main text, **Fig. 3E**). To the best of our knowledge, our study provides the first description of a *TP53-NTRK3* oncogenic fusion, and represents the first oncogenic gene fusion discovery made via single cell genomic sequencing.

#### 2.2 Characterizing altered ploidy in RPE cells by single cell sequencing

Our analyses of RPE cells, presented in Figure 2, revealed diagnostic footprints of altered chromosomal ploidy, namely monosomic and trisomic regions (**Figure 2C, Figure 2D**). Random and independent mitotic segregation of sister chromatids to daughter cells during anaphase (**Fig. 1B, Fig. S2**) implies that ploidy states can be predicted in strand-specific sequencing based solely on the relative fraction of W and C reads along a chromosome. A binomial distribution can be used to compute expected frequencies of template strand patterns for different ploidy states (**Table S4**). The principle of ploidy footprints is detailed as follows: For example, in the case of diploidy, 50% of all autosomes will show a characteristic pattern where one homolog is sequenced on the minus strand (W) and the other homolog is sequenced on the plus strand (C) – termed WC-pattern[2]. This produces a balanced strand orientation signal, or 1:1 strand ratio, of W and C reads for the autosome. The remaining autosomes are sequenced either only on the C strand (25%; CC-pattern), or only on the W strand (25%; WW-pattern), respectively (**Fig. S2**). Conversely, in the context of triploidy a CCC-pattern (all reads of an autosome map to the C-strand) and a WWW-pattern (all reads map to the W-strand) will be seen for 12.5% of all autosomes, respectively. The CWW-pattern and the CCW-pattern, resulting in a 1:2 (or 2:1) strand ratio, will each be seen for 37.5% of autosomes (**Table S4**). Notably, triploid segments will never exhibit a balanced 1:1 strand ratio. Tetraploidy and haploidy similarly result in their own readily discernible strand inheritance patterns, producing their own unique diagnostic footprints in scTRIP (**Table S4, Fig. S2**). Of note, in contrast to currently available methods, these diagnostic footprints do not require additional data for ploidy assignments, such as the detection of additional SVs on non-disomic chromosomes or comparison of depth-of-coverage values with a control - both of which are challenging to obtain in single cells. These footprints may be leveraged to study chromosome-specific aneuploidies as well as cellular ploidy.

We present the following evidence for the accuracy of these diagnostic footprints: When applied to RPE cells, our framework identified a chromosome arm-level CNA, particularly loss of 13q in C7, and additionally detected a gain of a large portion of 10q in RPE-1. As can be clearly seen in **Fig. 2CD**, the 13q-arm in C7 showed a 1:0 strand ratio consistent with monosomy, whereas the gained 10q region in RPE-1 exhibited 2:1 and 3:0 strand ratios consistent with trisomy (**Table S4**). All other autosomes showed prevalence of 1:1 and 2:0 strand ratios consistent with the near-diploid karyotype of the RPE lines[10,19]. The previously published karyotypes[10,19] of C7 and RPE-1 confirmed monosomy 13 and gain of a large region on 10q, respectively (**Table S3**).

#### 2.3 Construction of BM510: an RPE cell line showing genomic instability

To demonstrate the power of scTRIP to detect diverse classes of somatic SV we generated the BM510 cell line. To this end, we employed the CAST approach, which we previously established to transform parental RPE-1 cells[20]. Here, we subjected *TP53-/-* (knock-out generated using zinc finger nucleases[20]) hTERT-RPE1 cells[20] to siRNAs against the mitotic-spindle-associated protein Astrin[21] (Ambion) for 72 hours. Transfected cells were sorted using a MoFlo Legacy cell sorter (Beckman Coulter Inc.) equipped with a 100µm nozzle. After single cell sorting into 96-well plates, cells were grown into colonies, and then subjected to soft agar culture treatment (consisting of 0.5% bottom layer agar and 0.35% top layer agarose) to assay for *in vitro* transformation. The anchorage-independently growing cells were recovered from 96-wells, and re-cultured on 6-well plates. Single colonies were then isolated and grown, one of which yielded the BM510 cell line. In the context of our CAST screen, we subjected BM510 to low-coverage bulk WGS (<1x coverage) and verified the presence of somatic copy-number abnormalities. Subsequent single cell analysis of BM510 by scTRIP revealed a high level of genomic instability in this line, with many *de novo* formed and clustered SVs seen in individual cells (see main text; and *e.g.* **Figs. 3, 4**).

#### 2.4 Temporal ordering of SVs in single cells using the infinite sites assumption

We used the infinite sites assumption[36] to infer the temporal ordering of SVs affecting the same haplotype in single cells. The assumption made is that the probability of two independent internal SV breakpoints occurring at the exact same genomic position, and on the same haplotype, is zero. The theory and practical applications of this assumption are detailed in Li *et al.* For example, copy-number (CN) steps >1 (where a CN step refers to the magnitude of change in copy number at each breakpoint location) are, according to the infinite sites assumption, resulting from Del and Dup events that overlap with previously formed SVs (for examples, see **Fig. S15**). By comparison, newly formed SVs result in CN steps of 0, -1 or 1. While previously developed for bulk WGS data[36], we employed infinite sites assumption to infer the temporal ordering of overlapping SVs falling onto the same chromosome-length haplotype in single cells.

Application of the infinite sites assumption enabled us to predict a plausible temporal ordering of SVs in most cells showing more than one SV on the same haplotype (**Fig. S15**). In one case, we uncovered the formation of multiple inverted and lost fragments resulting in 12 SV breakpoints on the same chromosome 4 homolog/haplotype (**Fig. 4G)**, indicative for a one-off rearrangement burst (chromothripsis[39]) in a cell that additionally exhibited SVs on chromosome 8, 9, and X. In several other cases, the clustered rearrangements appeared to have formed through successive rounds of SV formation affecting the same homolog **(Fig. S15)**. This included cells where additional SVs were inferred to precede and succeed BFB cycle formation on the same homolog. These analyses indicate that via its ability to resolve SVs by whole-chromosome haplotype, scTRIP enables inference of sequentially arising and one-off complex rearrangement processes in single cells.

#### 2.5 Verification of SVs and karyotypes inferred by single cell sequencing in PDX-derived T-ALL samples

To verify the clonal and subclonal SV landscapes identified by scTRIP in the T-ALL samples, we investigated orthogonal datasets. Classical karyotyping of T-ALL samples P33 and P1 was pursued during diagnosis, at German study centers in the cities of Kiel and Gießen, respectively. We additionally performed exome-capture sequencing and reanalyzed recently generated exome sequencing data[22] from P33 and P1. These included samples taken during initial diagnosis, remission (interpreted as 'normal'), and relapse[23], enabling verification of scTRIP based karyotyping. These data additionally enabled us to compare CNAs seen during relapse with those present at initial diagnosis as well as with germline copy-number variants[22], which confirmed the presumed somatic status of these CNAs.

P33 whole exome sequencing data (number of reads aligned on exonic sequence per exome capture experiment: 60,919,787-63,897,672) were generated as described previously[22], following alignment to the human reference (hg19) using bwa[24]. Alignment files were sorted and indexed using samtools[25]. Quality control was pursued using Alfred (https://tobiasrausch.com/alfred), requiring at least 80% of all exonic sequence targets seen with >20x coverage. We calculated the coverage for each exonic target region requiring a minimum phred-scaled mapping quality of 20 using the 'count_dna' subcommand of Alfred. Binned coverage values were GC-normalized and adjusted by their respective coverage in the remission sample. Besides the normalized read-depth signal, we also called SNPs using FreeBayes[26]. To de-noise these raw SNP calls and their respective allele frequency, we phased all heterozygous SNPs present in the remission sample against the 1000 Genomes Project[27] SNP reference panel using Eagle2[28]. Phased, heterozygous germline SNPs were annotated with their variant allele frequency in the matched tumor genome to corroborate read-depth based CNA calls. We then utilised read-depth signal and SNP variant allele frequencies to verify, or invalidate, CNA calls made by our scTRIP framework. *De novo* calling from the whole-exome data was pursued for large CNAs >1Mb in size. In order to verify smaller CNAs we plotted the read depth signal for the inferred variant site and for flanking regions allowing verification of scTRIP based CNA calls. Bioconda[29] was used to install the aforementioned tools. To verify DNA rearrangements in P1, we utilized mate-pair sequencing[30] to confirm a one-off complex SV on 6q as well as a copy-number balanced inversion at 14q32. We performed SV detection using Delly[9].
